# A Hierarchical Cascade of Organellar Silencing and their Regeneration under Anaesthetic Stress in Plants

**DOI:** 10.1101/2025.07.03.663012

**Authors:** Shilpa Chandra, Sakshi Chouhan, Bodhidipra Mukherjee, Abdul Salam, Chayan Kanti Nandi, Laxmidhar Behera

**Affiliations:** Indian Knowledge System and Mental Health Applications Centre, Indian Institute of Technology Mandi, HP-175005; School of Biosciences and Bioengineering, Indian Institute of Technology Mandi, HP-175005; School of Chemical Science, Indian Institute of Technology Mandi, HP-175005; School of Computer Sciences and Electrical Engineering, Indian Institute of Technology Mandi, HP-175005

**Keywords:** Anaesthesia, Plants, Nuclear reorganization, Root apex, Organelle recovery, Causal structure, Hierarchical cascade

## Abstract

Anaesthetics are pharmacological drugs that temporarily inhibit neural activity by acting on voltage-gated sodium channels and GABA receptors. Although their neurological mechanisms are well-defined, their wider cellular effects, especially in non-neuronal systems, are inadequately understood d. This study utilized *Solanum lycopersicum* plant’s root apex cells as a transparent model to examine anaesthetic-induced subcellular alterations via live-cell fluorescence imaging, immunostaining, and super-resolution microscopy. Our findings, first of its kind, demonstrate the initial comprehensive model of sequential causal structure and hierarchical cascade of organelle collapse of different but important organelles such as mitochondria, lysosome, vesicles trafficking and nuclear architectures under anaesthesia in plants. The nucleus is identified as the most important controller of recovery potential and cellular fate. In a time dependent experiment, we found that plant cells exposed to lidocaine for up to four hours could still recover mitochondrial potential, lysosomal function, and nuclear integrity when anaesthesia is removed. However, beyond four hours, the damage, especially to the nucleus, was irreversible, and cells proceeded to programmed cell death. Our data further demonstrate that organelle functions can recover after brief exposure to anaesthetic stress. However, prolonged exposure prevents recovery, resulting in the irreversible degradation of the nucleus leading to complete cell death. The results present novel targets for preventative therapies, such as mitochondrial antioxidants, lysosomal stabilizers, and nuclear structural protectants that could mitigate anaesthetic-induced damage.

## Introduction

Anaesthesia has long served as a cornerstone of modern medicine, enabling surgical and diagnostic procedures through reversible suppression of consciousness and neural activity ^1,2^. Pharmacologically, anaesthetics such as lidocaine, etomidate, and propofol are known to act on specific neuronal targets, blocking voltage-gated sodium channels, enhancing GABA *A* receptor activity, or modulating chloride ion flux to inhibit excitatory neurotransmission ^1–3^. While these mechanisms are well-characterized in the context of the central nervous system, the broader cellular and subcellular consequences of anaesthetic exposure remain poorly understood, particularly in non-neuronal systems. This knowledge gap is significant, as it prevents a full understanding of anaesthetic toxicity and limits the development of protective strategies for preserving cellular integrity during clinical interventions. Notably, existing literature overwhelmingly centres on mitochondria as the primary site of anaesthetic-induced cytotoxicity^4^. In animal and neuronal models, prolonged exposure to anaesthetics such as propofol and etomidate leads to mitochondrial membrane depolarization, inhibition of Complex I activity, and excessive generation of reactive oxygen species (ROS) ^5^. These events start a process that leads to cell death, known as apoptosis, by releasing cytochrome c and activating caspases, making mitochondria the main area where drugs cause cells to die ^6,7^. However, this mitochondria-centric view does not fully capture the coordinated collapse of cellular systems observed under conditions of anaesthetic overdose. Other organelles, including lysosomes, endomembrane vesicles, the nucleus, and chromatin, are often overlooked, despite their well-established roles in intracellular signaling, homeostasis, and programmed cell death.

One major limitation of current research is the difficulty of real-time, organelle-specific observation in live mammalian systems. Technical and ethical constraints make it challenging to dissect subcellular interactions during progressive stages of chemical stress, particularly at the onset of irreversible damage. In this context, plant systems, specifically root apex cells offer a powerful and underexplored model. These cells possess transparent tissues and spatially organized organelles that are amenable to live-cell imaging. Furthermore, the root apex is considered the sensory and integrative hub of the plant, functionally analogous to a “brain,” making it a biologically relevant system for studying integrative subcellular behavior ^8,9^. Our study leverages this plant-based model to investigate how commonly used anaesthetics particularly lidocaine, affect organelle function beyond the nervous system. Despite their distinct molecular targets in neurons, these lipophilic compounds share the ability to diffuse across cell membranes and accumulate within mitochondria ^10,11^. In *Solanum lycopersicum* root apex cells, we observed that anaesthesia results in mitochondrial membrane depolarization and oxidative stress ^5^. This disruption is followed by a cascade of failures in other organelles vesicular transport is impaired, lysosomal activity declines, and ultimately, nuclear chromatin organization breaks down. These convergent effects suggest that mitochondria serve as an initial trigger for broader cellular collapse, but they are not the sole organelles affected. A cascade of failures in other organelles follows this disruption. Lysosomal acidification declines, vesicle trafficking is impaired, and autophagic buildup increases ^12,13^. Remarkably, the nucleus, despite being exposed to the same treatment, retains its integrity and chromatin organization during the early phases of stress. It remains the last organelle to show irreversible damage, reinforcing its role as the central command system of the cell ^14^. Cell death becomes inevitable when the breakdown of nuclear structure, fragmentation of chromatin, and loss of NRF2-mediated transcription occur (NRF 2 is a transcription factor that regulates the expression of antioxidant and detoxification genes in response to oxidative and cellular stress). In a time dependent experiment (0 h, 1h, 2h, 4h, 6h, 8h post-exposure), we found that plant cells exposed to lidocaine for up to four hours could still recover mitochondrial potential, lysosomal function, and nuclear integrity upon washout. However, beyond four hours, the damage, especially to the nucleus, was irreversible, and cells proceeded to programmed cell death. This scenario defines a critical temporal threshold for functional recovery and highlights the nucleus as the gatekeeper of cellular viability.

This sequence provides a causal framework for understanding multiorganelle failure in response to anaesthetic overload. The timeline established by our study enables precise mapping of these cellular events, offering a temporal resolution rarely achievable in animal models. This model helps us understand how different parts of the cell stop working together when exposed to anaesthetics. Organelles like mitochondria, lysosomes, and the nucleus normally work as a team to maintain balance, handle stress, and support recovery. When anaesthetics are applied for too long, this teamwork breaks down in a specific order starting with energy loss, followed by problems in waste removal, communication, and finally, breakdown of the nucleus. Studying this step-by-step failure in plant cells, which are easy to image and don’t raise ethical concerns, gives us a clear view of how cells respond to stress. It also helps identify when damage becomes permanent and which parts of the cell could be targeted to prevent it. Furthermore, our findings have translational relevance. By highlighting the vulnerabilities of mitochondria, lysosomes, vesicles, and the nucleus under anaesthetic stress, we identify potential targets for protective interventions. Compounds such as mitochondrial antioxidants like MitoQ and SS-31 to preserve ETC function^15^, TFEB activators and lysosomal stabilizers to sustain degradation pathways ^16^, and nuclear protectants such as sirtuin activators ^17^ and HDAC inhibitors to maintain chromatin integrity ^18^. Furthermore, NRF2 pathway activators and autophagy inducers could offer organelle-specific protection during anaesthesia, reducing the risk of long-term cellular damage in clinical settings.

## Results and discussions

### Mitochondria as the Primary Target Initiating the Stress Cascade

Mitochondria are key regulators of energy metabolism, redox balance, and cell survival. To see how mitochondria are put together and how they work, we used MitoTracker Green (MTG) for mitochondria and Tetramethylrhodamine Ethyl Ester (TMRE) for membrane potential ^5,19–21^. In 0 hour cells, both probes showed strong, even signals, which showed that the mitochondria were still whole **(Figure 1Ia, Ib)**. With increasing anaesthetic concentration, TMRE intensity declined, indicating mitochondrial depolarization, ATP depletion, and early stress signaling **(Figure 1Ib–f)** ^5,22,23^. There was a big drop in the TMRE and MTG signals (\*\*\**p* < *0*.*001*, two-tailed paired t-test) in **Figure 1III–VI**, which showed that the mitochondria were breaking down. There was a rise in the Pearson correlation analysis **(Figure VII–IX)** between mitochondria and lysosomes at first, which showed mitophagy. Then there was a fall, which showed lysosomal stress and mitochondrial loss. Following mitochondrial depolarization, the cell initiates autophagy to eliminate damaged mitochondria. We used LysoTracker Red (LTR) to assess lysosomal activation^12,24^. LTR signal increased progressively **(Figure 1IIb–d)**, indicating autophagosome formation and mitophagy. Later on, though, persistent LTR fluorescence without corresponding recovery of TMRE/MTG signals **(Figure 1IIe–f)** showed that autophagic activity was no longer at work. Quantitative fluorescence analysis confirmed this mismatch, highlighting the fact that cells can’t fix themselves after a certain point.

**Figure 1:**
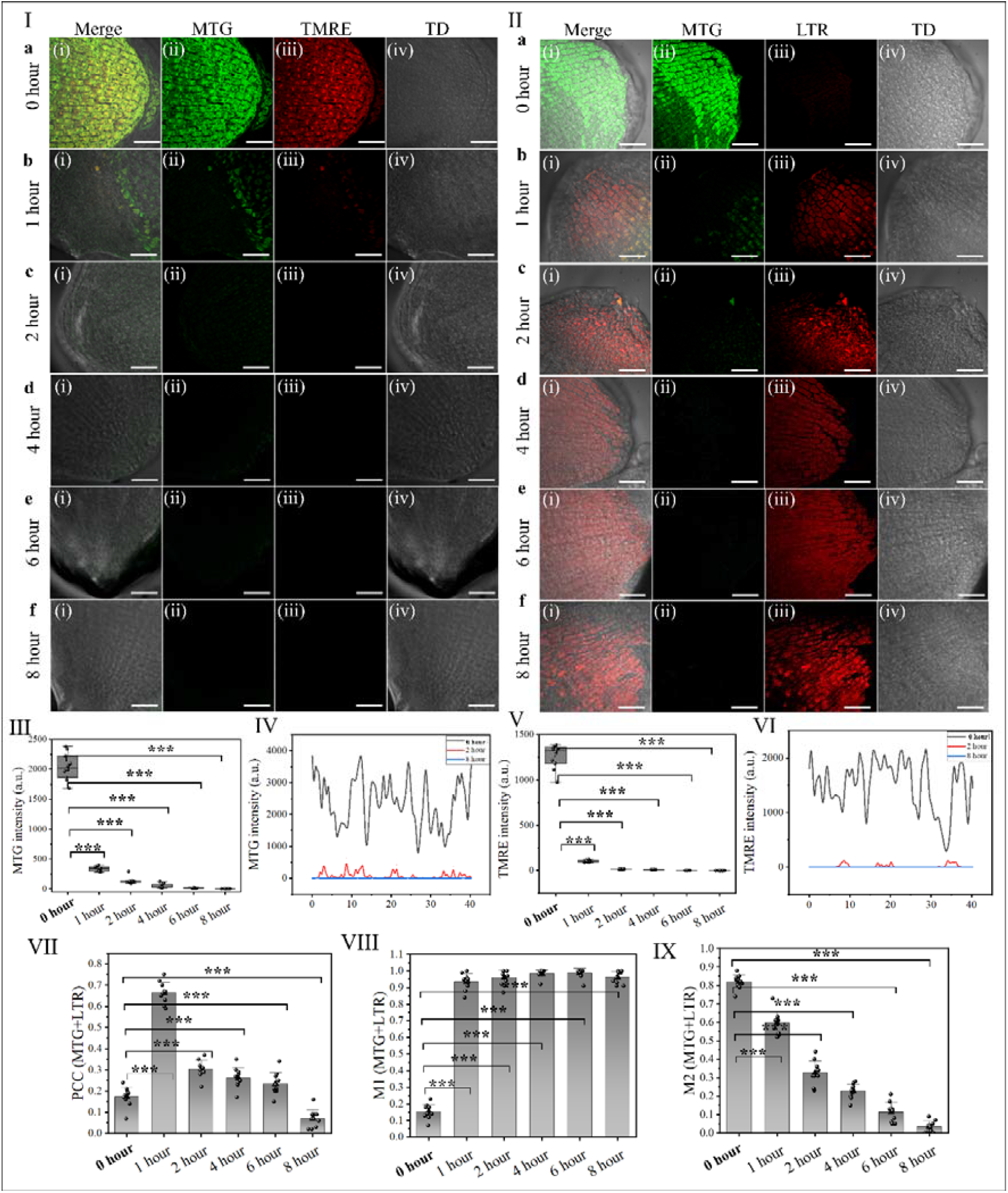
Effects of prolonged anaesthesia exposure on mitochondrial integrity and lysosomal activity. **(I)** Mitochondrial membrane potential analysis using TMRE (red) and mitochondrial structure visualization with MTG (green) over different time points. (i) Merged images, (ii) MTG staining, (iii) TMRE fluorescence. 0-hour cells (a) exhibit intact mitochondria, while TMRE and MTG fluorescence progressively decline from 1 to 8 hours (b–f), indicating mitochondrial dysfunction. **(II)** Lysosomal activity assessment using LysoTracker Red (LTR) and MTG. (i) Merged images, (ii) MTG staining, (iii) LTR fluorescence. 0-hour cells (a) display strong MTG and LTR signals, while lysosomal accumulation intensifies after 2 hours, suggesting stress-induced lysosomal activation. Quantitative analysis of MTG and TMRE fluorescence intensities shows a significant decline from 1 hour onwards, with a sharp drop after 2 hours, confirming mitochondrial deterioration **(III-VI)**.

Additionally, a recovery experiment was performed to assess whether mitochondrial integrity and function could be restored after anaesthesia exposure. Following 1 hour of anaesthesia treatment, mitochondria recovered within 1 hour **(Figure S1 Ia-b)**. It took 4 hours for cells treated for 2 hours to recover **(Figure S2 Ia–d)**, but 8 hours for cells treated for 4 hours to recover mitochondria **(Figure S3 Ia–g)**. However, for cells exposed to 6 hours of anaesthesia, no recovery was observed, indicating that mitochondrial damage was irreversible beyond this threshold **(Figure S4 Ia-f)**.

### ROS Generation Redox Collapse and Oxidative Damage Lead to Communication Breakdown

To be sure that mitochondrial failure caused systemic stress, we measured reactive oxygen species (ROS) using HlJDCFDA, a reliable dye that can detect ROS ^25^. ROS levels remained low in 0 hour cells but increased steadily with higher lidocaine exposure **(Figure 2Ia–f)**. Quantitative ROS intensity analysis **(Figure 2III–IV)** showed highly significant increases (\*\*\****p* < *0*.*001***) ^26^. Recovery experiments indicated that ROS levels returned to baseline only for exposures up to 4 hours, while longer exposures resulted in irreversible oxidative stress **(Figures S5–S8)**. We visualized endocytic vesicle trafficking using FM4-64, a fluorescent dye that stains plasma membranes and endocytic vesicles. With increasing anaesthesia exposure, vesicle area and fluorescence intensity significantly declined **(Figure 2IIa–f)**, indicating that membrane transport and communication pathways were disrupted by oxidative stress ^10,13,27^. Quantitative image analysis **(Figure 2V–VII)** confirmed this decrease with statistical significance (\*\*\****p < 0*.*001***). These results suggest that long-term ROS accumulation makes it harder for vesicles to form, which separates organelles and makes it even harder for cells to work together. Vesicle formation was irreversibly inhibited after 6 hours, even though it recovered in 2–8 hours for exposure up to 4 hours **(Figures S9–S11) (Figure S12)**.

**Figure 2:**
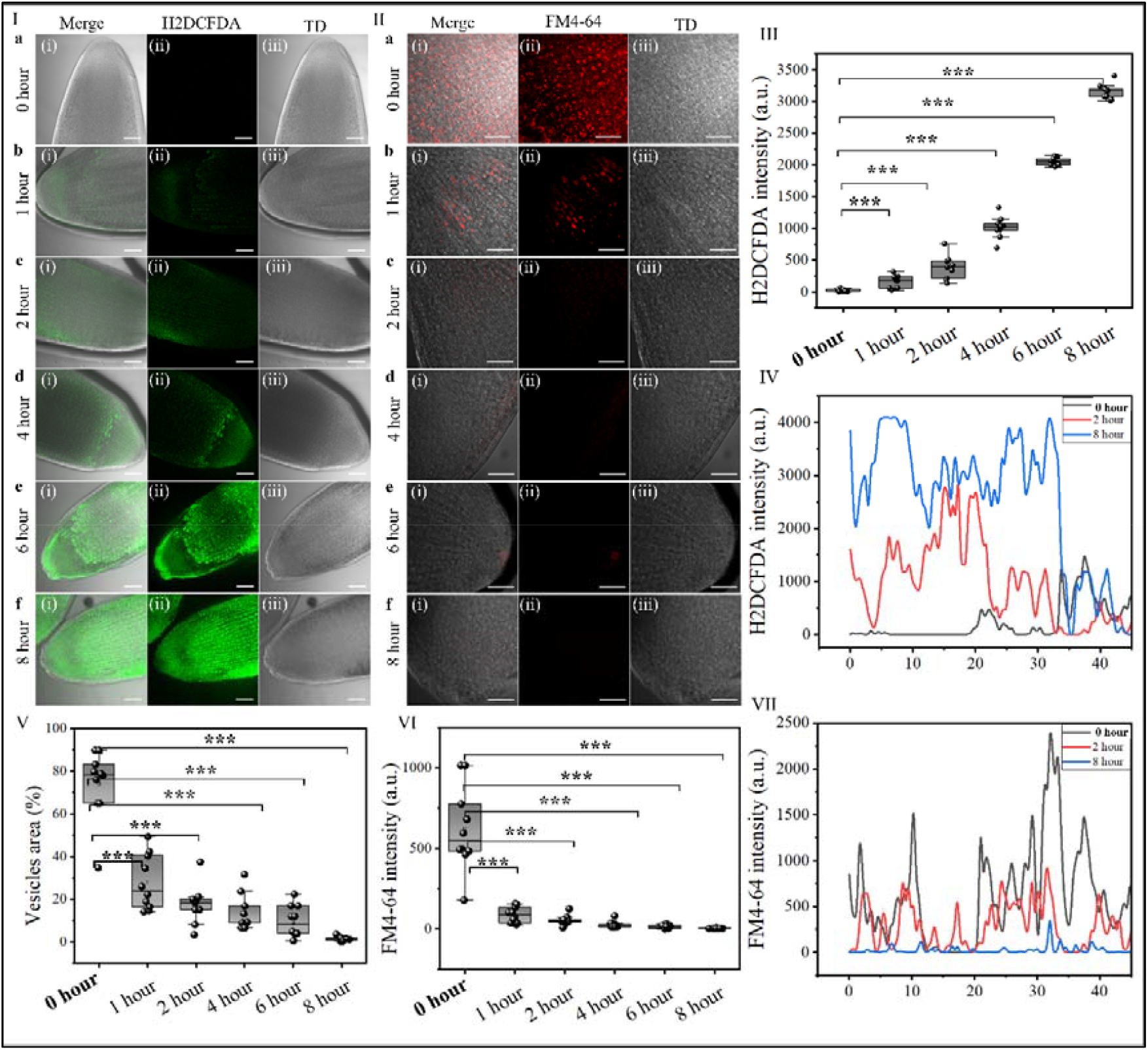
Effects of anaesthesia exposure on ROS accumulation and vesicle dynamics in plant root cells. **(I)** H□DCFDA staining for ROS detection over increasing exposure durations. (i) Merged images, (ii) H□DCFDA fluorescence indicating ROS accumulation, (iii) Transmitted light (TD) images. ROS levels increase progressively from 1 to 8 hours. **(II)** FM4-64 staining for vesicle dynamics assessment. (i) Merged images, (ii) FM4-64 fluorescence marking vesicles, (iii) TD images. Vesicle formation initially rises but declines after prolonged stress. **(III)** Normalized intensity of H□DCFDA fluorescence shows a significant increase over time. **(IV)** Intensity plot along a selected line highlights H□DCFDA signal variations in 0 hour, 2-hour, and 8-hour treatments. **(V)** Quantification of vesicle area percentage over time. **(VI)** FM4-64 fluorescence normalised intensity significantly decreases after 2 hours, indicating reduced vesicle activity. **(VII)** Intensity plot of FM4-64 fluorescence along a selected line in 0 hour, 2-hour, and 8-hour conditions. Statistical significance determined by two-tailed paired t-test (*p < 0.05, **p < 0.01, ***p < 0.001). Scale bars = 20 μm.

### Nuclear Stress Response and NRF2 Translocation

To determine whether cells initiate nuclear defences, we tracked NRF2, a master regulator of the oxidative stress response ^28–30^. Under stress, NRF2 translocates from the cytoplasm to the nucleus. Immunolocalization studies **(Figure 3IIa–c)** showed increased nuclear localization of NRF2 with higher anaesthetic concentrations, indicating an adaptive nuclear response. In later stages, though **(Figure 3IId–f)**, chromatin condensation happened at the same time as a persistent NRF2 signal ^28,31^. This suggests that the mechanisms that were protecting the cells have broken down. DAPI staining showed that the nuclei were getting smaller and more broken up **(Figure 3III)** ^32^, and image analysis **(Figures 3IV–IX)** demonstrated that the size and brightness of the nuclei had changed significantly **(\**p* < *0*.*05 to* \*\*\**p* < *0*.*001*)**. Nuclear integrity is restored 6–8 hours after brief anaesthesia exposure, according to recovery experiments **(Figures S13–S15)**. But after six hours, there is no nuclear recovery **(Figure S16)**, which results in irreversible chromatin damage and cell death ^33–35^. These results demonstrate that under prolonged oxidative stress, NRF2-mediated defence is inadequate, leading to nuclear collapse and cell death. We also examined nuclear organization under general anaesthesia (etomidate) and observed a nuclear arrangement pattern comparable to that seen with local anaesthesia (lidocaine), indicating a shared effect on nuclear architecture despite differing molecular targets **(Figure S17)**.

**Figure 3:**
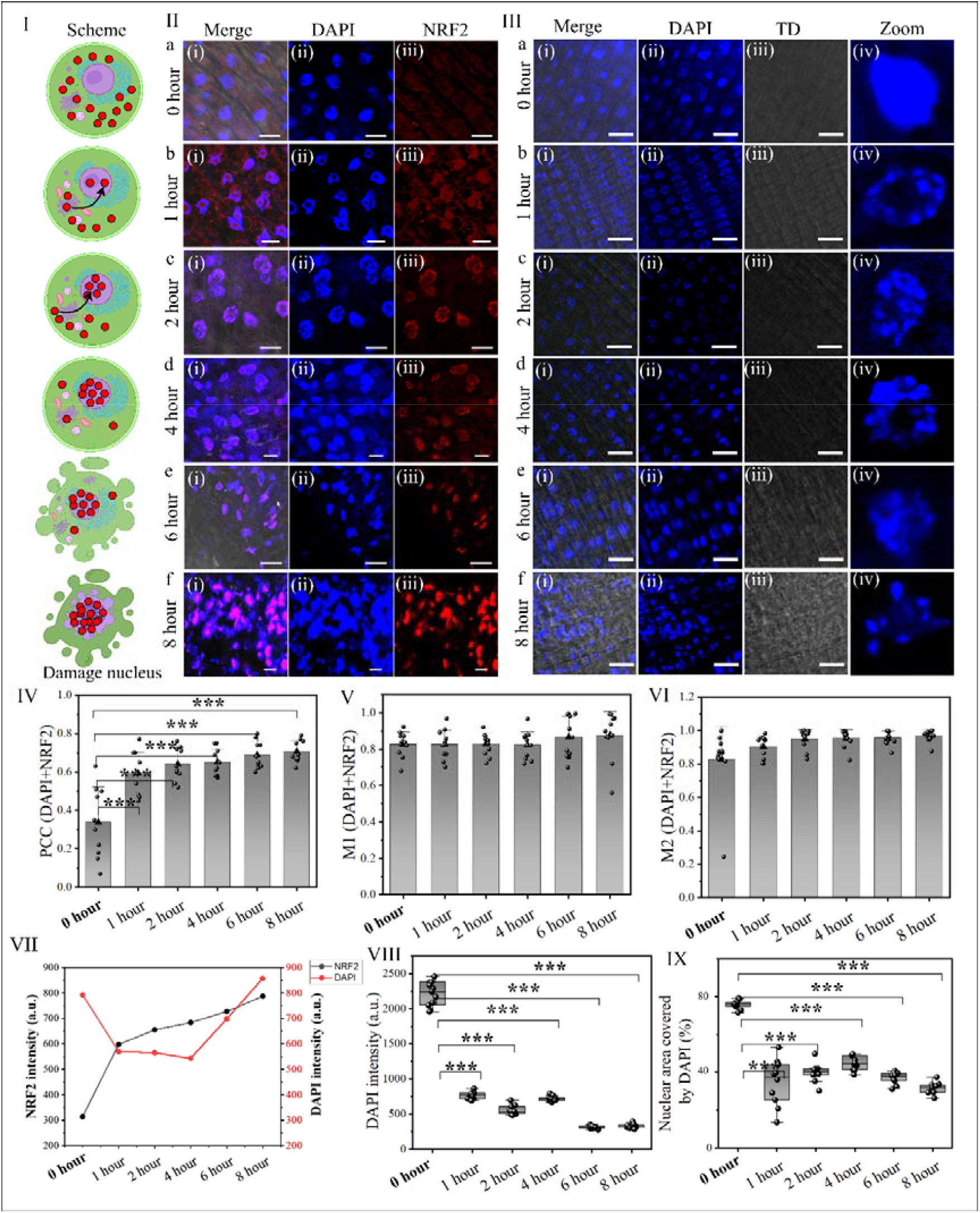
NRF2 (Red dot) Nuclear Translocation and Nuclear Integrity Under Anaesthesia Stress. **(I)** Schematic representation of NRF2 translocation into the nucleus following stress exposure. **(II)** Immunofluorescence analysis of NRF2 expression in plant cells over increasing durations of anaesthesia exposure. (i) Merged images, (ii) DAPI-stained nuclei, and (iii) NRF2 fluorescence. 0-hour cells (a) show normal NRF2 distribution, while nuclear accumulation of NRF2 increases from 1 to 8 hours (b–f). **(III)** Nuclear morphology assessment using DAPI and transmitted light (TD) imaging. (i) Merged images, (ii) DAPI-stained nuclei, (iii) transmitted light images, and (iv) zoomed-in nuclear regions. Prolonged exposure leads to nuclear deformation and chromatin condensation. **(IV–VI)** Quantification of NRF2 colocalization with DAPI using Pearson’s correlation coefficient (PCC), Manders’ coefficient 1 (M1), and Manders’ coefficient 2 (M2). **(VII– IX)** DAPI and NRF2 intensity measurements show a significant decrease in nuclear area covered by DAPI over time, indicating nuclear deformation and chromatin condensation. (*p < 0.05, **p < 0.01, ***p < 0.001). Scale bars = 20 μm.

### Chromatin Reorganization from Transcriptional Pause to Structural Collapse

To understand how this nuclear stress response change’s function, it is important to look at how chromatin is organized and the epigenetic changes that manage gene expression and genome stability. There were two changes we were interested in: H3K4me3, which is connected to active euchromatin and transcriptional activity, and H3K9me3, which is connected to heterochromatin that stops transcription ^36^. Stress signals sent by NRF2 can either cause an adaptive transcriptional response or happen at the same time as transcriptional shutdown and chromatin compaction. These are signs that a cell has made a decision about its fate that can’t be changed, such as programmed cell death (PCD) ^37–40^.

Staining 0-hour cells with H3K4me3 revealed clear, well-structured euchromatin domains. This meant that the nucleus was stable and actively transcribed **(Figure 4Ia)** ^23^. However, with increasing anaesthesia concentration, H3K4me3 fluorescence intensity and chromatin domain area progressively declined. Along with the drop, the perimeter and circularity got smaller, which showed a shift toward chromatin condensation and transcriptional silencing **(Figure 4Ib–f)** ^41–44^. Quantitative analysis showed a big drop in the area of euchromatin, the number of domains, the geometric features, and other things **(\**p* < *0*.*05 to* \*\*\**p* < *0*.*001)* (Figure 4II–IX)**. At the highest concentrations, there was a small increase in the number of domains, which was probably due to chromatin fragmentation rather than functional reorganization.

**Figure 4:**
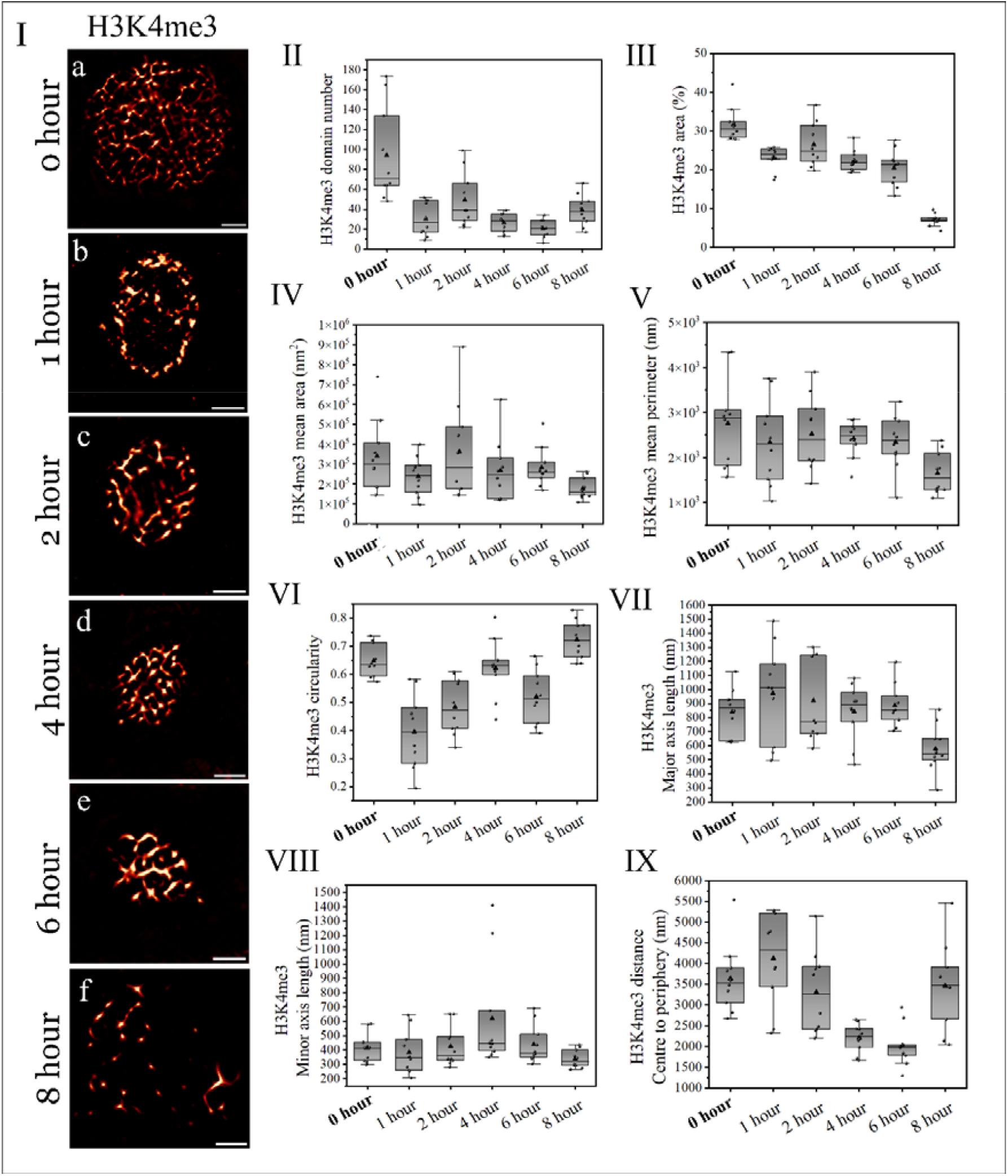
H3K4me3 Distribution and Nuclear Chromatin Organization Under Prolonged Anaesthesia Exposure. **(I)** Representative images of H3K4me3-stained nuclei from 0 hour **(a)** and anaesthesia-exposed root cells for 1–8 hours **(b–f)**. Scale bars: 2 μm. **(II)** H3K4me3 domain number decreases significantly with prolonged exposure, indicating chromatin reorganization. **(III)** H3K4me3-stained area (%) reduces over time, suggesting loss of active chromatin regions. **(IV)** H3K4me3 mean area declines, reflecting condensation and compaction of euchromatin. **(V**) H3K4me3 mean perimeter shows variability, indicating changes in chromatin domain morphology. **(VI)** H3K4me3 circularity fluctuates, suggesting alterations in chromatin structure. **(VII & VIII)** H3K4me3 major and minor axis length decrease, showing progressive chromatin condensation. **(IX)** H3K4me3 centroid-to-periphery distance significantly shortens, indicating a shift toward perinuclear chromatin localization. Independent t-test show all the data significant *p < 0.05, **p < 0.01, ***p < 0.001. Scale bars = 2 μm.

On the other hand, H3K9me3 staining in 0-hour nuclei showed a moderate spread. This means that heterochromatin is changing and spreading out **(Figure 5Ia)**. As anaesthetic concentration increased, H3K9me3 domains exhibited initial compaction followed by signal weakening, structural disorganization, and eventual nuclear occupancy loss **(Figure 5Ib–f)**. Significant decreases in domain area, count, and minor axis length **(\**p* < 0.05 to \*\**p* < 0.001) (Figure 5II–VIII)**, along with a shift of heterochromatin toward the nuclear centre **(Figure 5IX)**, indicated nuclear architectural collapse ^33^.

**Figure 5:**
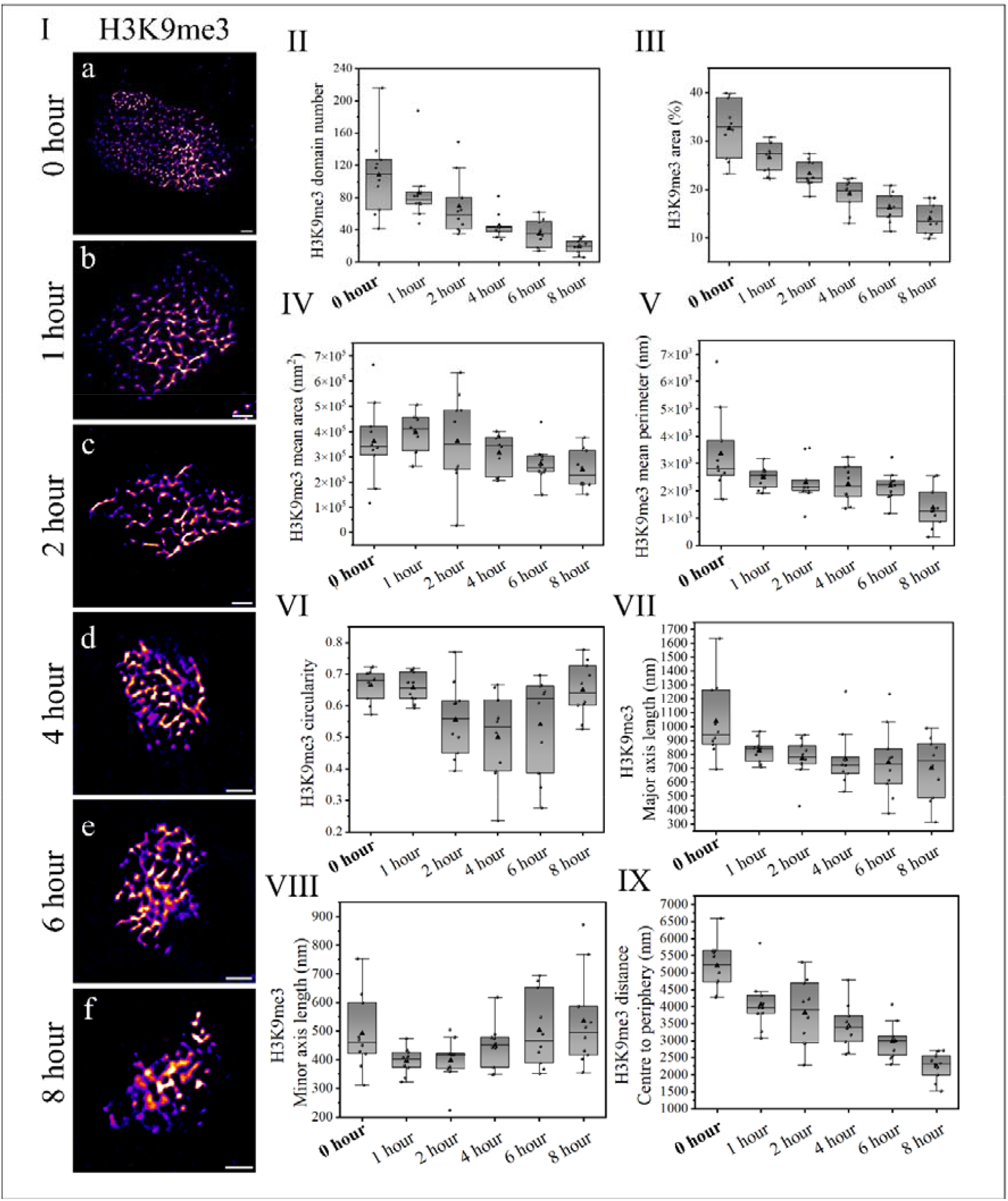
H3K9me3 Distribution and Heterochromatin Organization Under Prolonged Anaesthesia Exposure. **(I)** Representative images of H3K9me3-stained nuclei from 0 hour (a) and anaesthesia-exposed root cells for 1–8 hours **(b–f)**. Scale bars: 2 μm. **(II)** H3K9me3 domain number decreases significantly, indicating heterochromatin reorganization. **(III)** H3K9me3-stained area (%) reduces over time, reflecting chromatin compaction. **(IV)** H3K9me3 mean area shows a decreasing trend, indicating heterochromatin condensation. **(V)** H3K9me3 mean perimeter decreases with prolonged exposure, suggesting altered chromatin structure. **(VI)** H3K9me3 circularity fluctuates, reflecting nuclear structural changes. **(VII & VIII)** H3K9me3 major and minor axis length decline, showing progressive chromatin condensation. **(IX)** H3K9me3 centroid-to-periphery distance significantly shortens, indicating perinuclear localization of heterochromatin. Scale bars = 2 μm.

Correlation matrix analysis revealed a clear causal sequence in chromatin remodeling. In H3K4me3-labeled euchromatin, increasing anaesthetic stress disrupted positive correlations between domain area, perimeter, and axis length, reflecting a transition from chromatin loosening to condensation and transcriptional suppression **(Figure 6I)**. Conversely, H3K9me3 correlations showed a steady rise in structural order, followed by terminal compaction and rigidity, confirming repressive chromatin reorganization **(Figure 6II)**. This biphasic response highlights a central theme in the causal structure of nuclear stress: euchromatin initially undergoes disorganization, possibly to allow stress-responsive gene expression, but ultimately condenses and fragments under persistent stress. Heterochromatin, on the other hand, transitions from a dispersed to a condensed and transcriptionally inert state. These distinct epigenetic responses converge to a point of irreversible nuclear remodeling and loss of regulatory plasticity.

**Figure 6:**
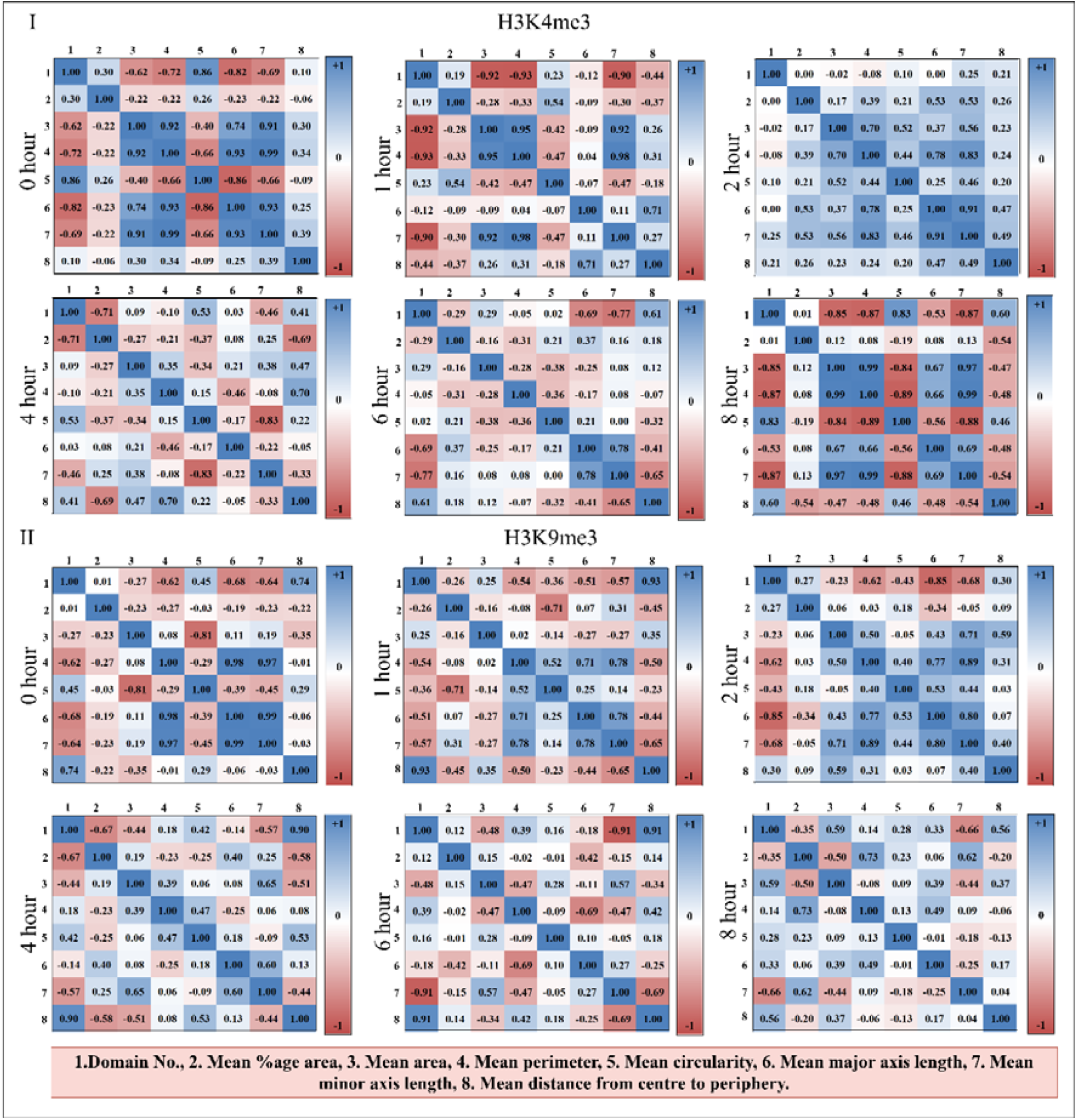
Correlation matrices for H3K4me3 (I) and H3K9me3 (II) modifications across different time points (0 hour, 1hr, 2hr, 4hr, 6hr, 8hr) post-anaesthesia. Blue indicates positive correlations, and red indicates negative correlations between chromatin parameters (Domain Number, Area, Perimeter, Circularity, Axis Lengths, and Distance from Centre). H3K4me3 shows early relaxation followed by condensation, while H3K9me3 exhibits progressive compaction, reflecting distinct chromatin remodeling dynamics under stress.

### Final Commitment to Cell Death

As anaesthesia-induced stress accumulated, it became increasingly evident that cells were unable to maintain homeostasis, eventually progressing toward irreversible damage. To find out how much cell death was caused by long-term lidocaine exposure, we used PI staining to see how many dead cells there were and DAPI staining to see how the nuclei were shaped ^45^. This dual approach enabled the precise identification of stress-induced cell death in the root apex cells of *Solanum lycopersicum* by linking plasma membrane integrity with nuclear structure **(Figure 7I)**.

**Figure 7:**
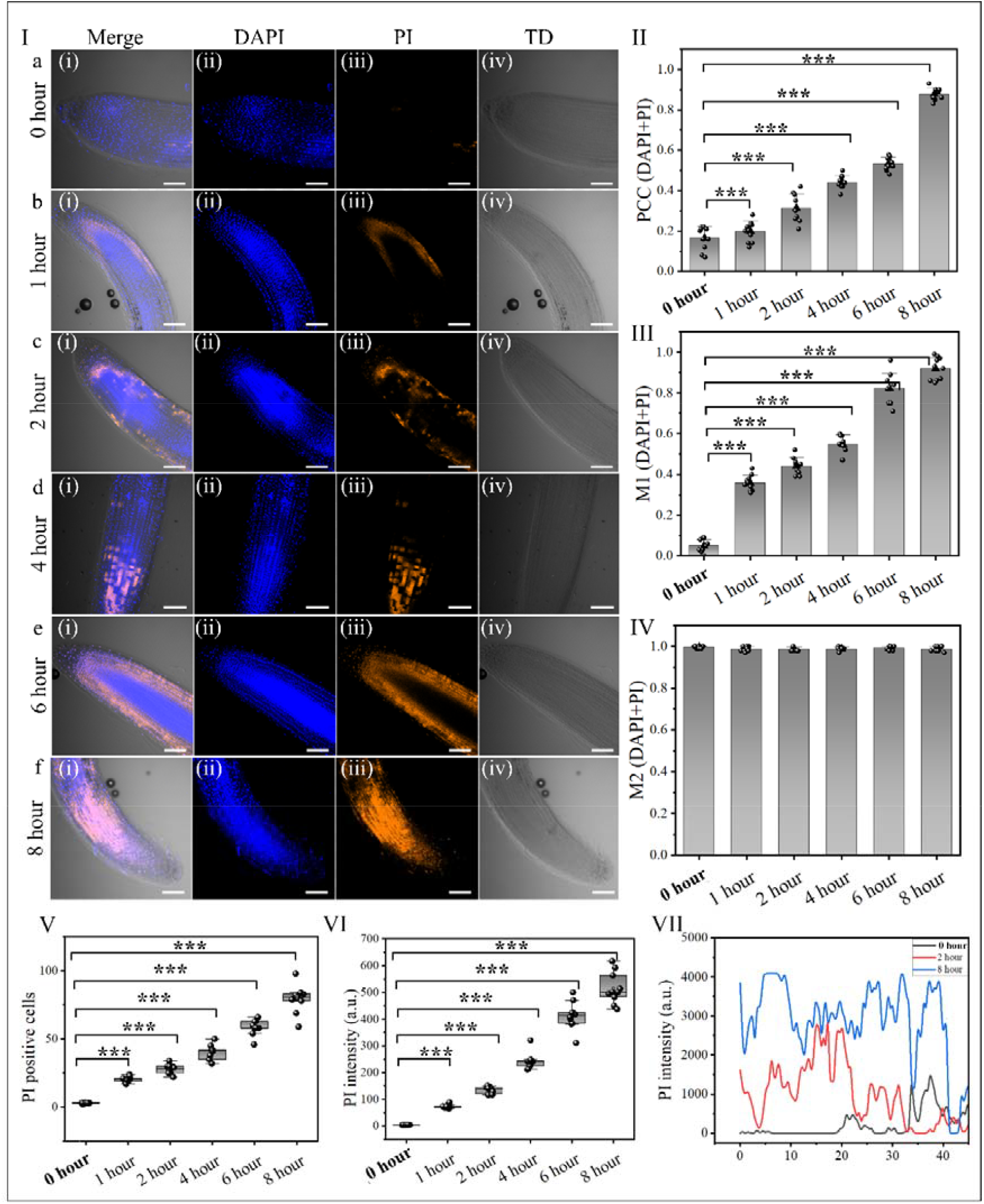
Assessment of nuclear integrity and cell death in plant root cells under anaesthesia exposure. **(I)** Representative images of root cells at different time points. (i) Merged images, (ii) DAPI-stained nuclei (blue), (iii) PI-stained dead cells (orange), and (iv) transmitted light (TD) images. PI fluorescence increases over time, indicating progressive cell death from 1 to 8 hours. **(II-IV)** Quantification of colocalization between DAPI and PI using Pearson’s correlation coefficient (PCC), Manders’ coefficient 1 (M1), and Manders’ coefficient 2 (M2), showing a significant increase in colocalization, suggesting increased nuclear membrane permeability and DNA fragmentation. **(V-VI)** Quantification of PI-positive cells and PI fluorescence intensity, confirming a time-dependent increase in cell death. **(VII)** Line plot showing PI intensity changes across different time points, highlighting nuclear membrane damage and increased permeability. Statistical significance determined by two-tailed paired t-test (\**p* < *0*.*05*, \*\**p* < *0*.*01*, \*\*\**p* < *0*.*001*). Scale bars = 20 μm.

In 0 hour cells, the nuclei were whole and clearly defined, and there was no PI signal to be seen. This means the cells were healthy and alive **(Figure 7Ia)**. However, with increasing anaesthetic exposure, PI-positive cells began to appear, marking the onset of membrane compromise and early cell death **(Figure 7Ib)**. As stress intensified, the frequency and fluorescence intensity of PI-positive nuclei increased significantly **(Figure 7Ic–d)**, indicating a correlation between nuclear damage and cell death ^6,26,46,47^. Pearson correlation and M1 analysis further validated this relationship **(Figure 7IV–V)**. By higher concentrations of anaesthesia, the lateral root cap (LRC), a region known for active developmental programmed cell death (dPCD), showed widespread PI staining **(Figure 7Ie)**, and by the final stage, most LRC cells exhibited nuclear fragmentation and a strong PI signal, confirming advanced cell death **(Figure 8If)**. While some degree of dPCD is typical in the LRC, anaesthesia appeared to accelerate and amplify this process. Statistical evaluation using a two-tailed paired t-test confirmed significant differences across conditions **(\**p* < *0*.*05, p* < *0*.*01*, \**p* < *0*.*001*) (Figure 7II– VII)**, underscoring the detrimental effects of sustained anaesthetic exposure on root cell viability.

**Figure 8:**
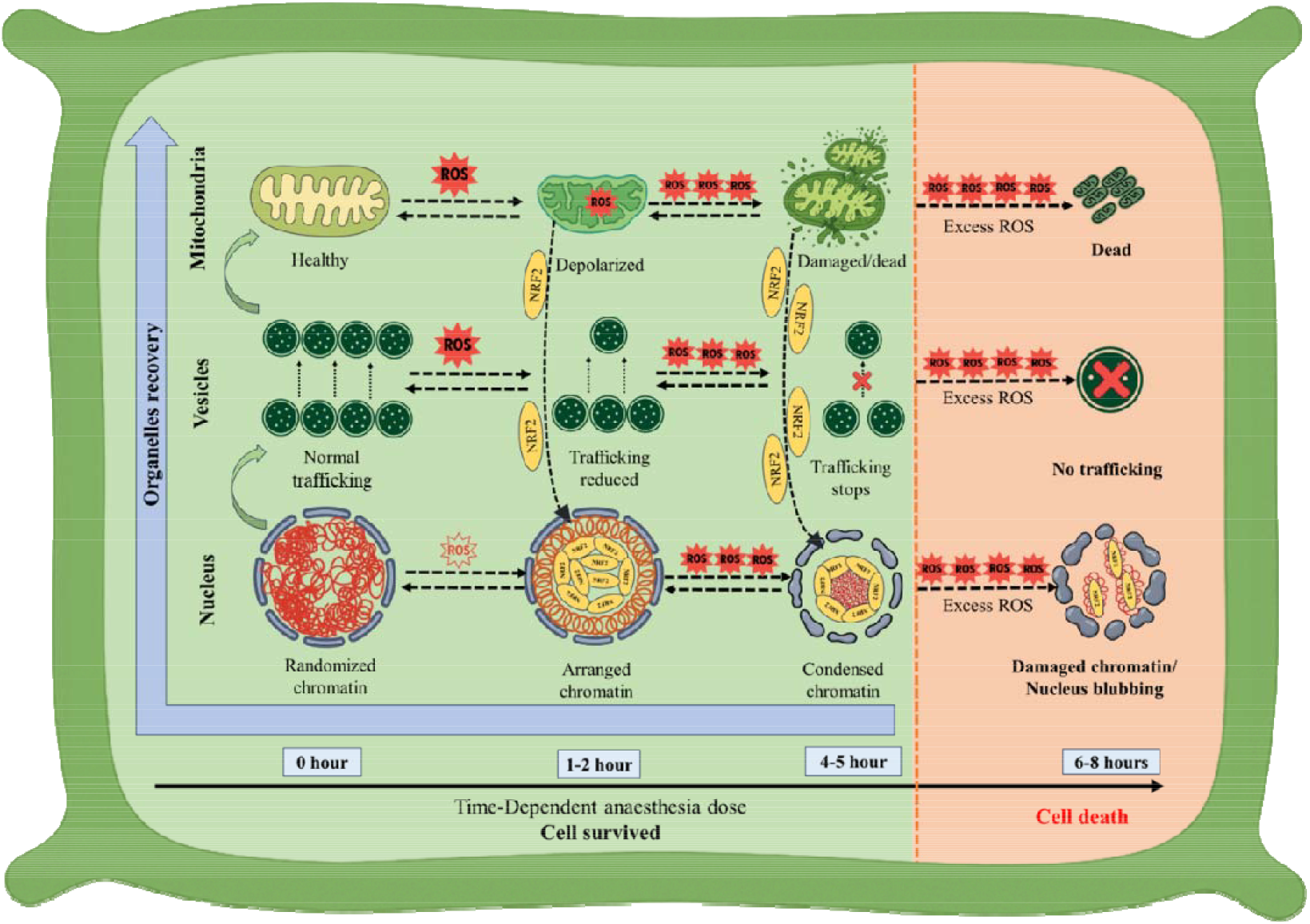
Illustrates the progressive cellular events observed during prolonged anaesthesia exposure, correlating cellular damage with the causal structure of cell death and its recovery.

### From Energy Failure to Genome Instability Defining the Cellular Point-of-No-Return

Our observations reveal a sequential cascade of events of cell death and recovery under lidocaine ^26^. Mitochondrial membrane depolarization is the earliest indicator of dysfunction, signifying an energy crisis that precedes cell-wide collapse ^5,23^. This depolarization leads to the excessive production of reactive oxygen species (ROS), hallmark indicators of redox imbalance and a trigger for downstream organelle and nuclear stress^13,27^. ROS accumulation interferes with vesicle trafficking, disrupting intracellular communication and transport ^10^. This separation makes organelles more stressed, reducing the ability of lysosomes to break down waste and causing a buildup of autophagic materials ^48^. A notable adaptive response is the translocation of NRF2, a master regulator of oxidative stress, to the nucleus ^28,29^. While the pattern suggests an initial attempt to counter oxidative damage via upregulation of detoxifying and antioxidant genes, the system becomes overwhelmed upon prolonged exposure. Nuclear responses fail beyond a critical exposure threshold of approximately 4–6 hours, leading to a progressive breakdown of nuclear architecture characterized by shrinkage, chromatin condensation, and fragmentation ^14,23,33,42–44^. As the nuclear arrangement seen in local anaesthesia same arrangement we have found with general anaesthesia.

Crucially, this study highlights that chromatin reorganization is not merely a terminal event in programmed cell death (PCD) but a key transition state in the progression toward cellular collapse. Immunostaining for euchromatin (H3K4me3) and heterochromatin (H3K9me3) shows a two-phase response: first, euchromatin relaxes to enable changes in gene activity due to stress, then it becomes compacted and broken apart; meanwhile, heterochromatin gradually tightens toward the centre of the nucleus. This result suggests that the chromatin landscape dynamically adapts to stress before structural destabilization seals the cell’s fate. The causal structure established in this study— **mitochondrial depolarization** → **ROS accumulation** → **vesicle trafficking inhibition** → **NRF2 translocation** → **chromatin reorganization** → **PCD** provides a hierarchical map of organelle vulnerability. Not only is this pathway consistent with known stress responses in both plants and animals, but it also opens the door to comparative frameworks for understanding “cell death” across biological kingdoms. The nucleus appears to serve as the final decision-making hub: its ability to preserve integrity under stress delays PCD, whereas its collapse marks the point of no return. During the post-stress recovery phase, a distinct temporal sequence of organelle and cellular restoration was observed. Mitochondria were the first to show signs of recovery, with restoration of membrane potential and network integrity. This was followed by lysosome-mediated clearance of damaged organelles via autophagy, indicating activation of intracellular housekeeping pathways. As ROS levels declined, detoxification mechanisms were re-established, likely under the influence of NRF2 signaling. Subsequently, vesicular trafficking resumed, enabling membrane repair and cargo redistribution. Finally, nuclear reorganization was evident, marked by chromatin decondensation and reassembly of nuclear architecture. This orchestrated sequence facilitated cellular survival and homeostasis re-establishment.

This understanding also has broader implications. Our data identify time-specific thresholds of damage and windows of reversibility. Mitochondrial and lysosomal functions are recoverable up to 4 hours post-exposure, but nuclear fragmentation beyond this duration leads to irreversible cell death. These thresholds define intervention windows and reveal the plasticity of the plant cell’s stress-adaptation machinery. These insights can help develop strategies for dealing with stress and can be applied in medicine, such as using NRF2 activators or antioxidants aimed at mitochondria like MitoQ and SS-31 to protect cells ^15^. Importantly, our findings offer a unique biological platform to explore anaesthesia induced cell death from a systems biology perspective. At the subcellular level, this capacity may be mirrored by the ability of organelles to interact and coordinate metabolic and genetic responses to stress. When cells are exposed to too much anaesthetic, the normal communication between their parts starts to break down. This loss of coordination makes it harder for the cell to manage stress and stay alive. By studying this process, we can learn more about how cells handle damage and what causes them to fail. This understanding could help future research on how different parts of the cell work together during stress in many types of living organisms.

## Conclusion

This study provides a comprehensive, time-resolved dissection of the cellular effects of prolonged anaesthesia in *Solanum lycopersicum* root apex cells, revealing a clear causal sequence of subcellular dysfunction leading to programmed cell death. By combining fluorescence imaging, immunostaining, and super-resolution microscopy, we demonstrate that mitochondrial depolarization acts as the earliest and most sensitive indicator of anaesthetic and eventual failure of nuclear stress responses mediated by NRF2. The process of changing the structure and arrangement of chromatin shows an important point where euchromatin tightens, heterochromatin becomes more compact, and the nucleus breaks apart. These results support a hierarchical model in which the nucleus integrates stress signals from organelles and governs the balance between recovery and collapse. This model captures the dynamics of stress progression, resilience, and fate decisions, whether driven by energy failure or chromatin instability. Beyond advancing our understanding of plant stress physiology under anaesthesia, this work also presents a simplified yet powerful model system with broader applicability. It offers a novel platform for evaluating cytotoxic effects of anaesthetics, screening stress-modulating compounds, and probing cross-kingdom parallels in stress integration. These findings open new directions for clinical anaesthesia research, plant resilience engineering, and bio-inspired studies on subcellular decision-making under extreme stress.

## Data and Materials Availability

All primary data supporting the findings of this study are included in the main manuscript and its supplementary information.

## Acknowledgements

The authors sincerely thank the Advanced Materials Research Centre (AMRC) and the Indian Knowledge System and Mental Health Applications (IKSMHA) Centre at the Indian Institute of Technology Mandi (IIT Mandi) for providing access to advanced facilities and sophisticated imaging and analytical instruments essential for this study. We gratefully acknowledge the Ministry of Education (MoE), Government of India, for providing research scholarships to the contributing authors. CKN, SC, and BM also acknowledge the financial support received from the IKSMHA Centre, IIT Mandi.

## Author Contributions

SC conceived and designed the experiments with inputs from CKN and LB, optimized experimental protocols, performed all plant-related experiments, imaging, and data analysis, and also wrote the manuscript. SCH contributed by assisting in statistical evaluation. BM supported the study by contributing to the design some experimental setups and manuscript inputs. AS provided assistance in high-resolution imaging and technical support. CKN and LB supervised the overall project, offered continuous guidance throughout the research, and contributed to the conceptual development and editing of the manuscript. All authors read, reviewed, and approved the final version of the manuscript.

## Ethical Statement

This research did not involve any human participants or animal subjects; therefore, ethical clearance was not required. The study was conducted entirely on plant materials (*Solanum lycopersicum* and *Solanum melongena*) under controlled laboratory conditions to investigate physiological responses to anaesthetic treatment. No genetically modified organisms (GMOs) were used in any part of the study. All experimental procedures complied with institutional and national guidelines for plant research. Pearson correlation coefficient (PCC) values for MTG and LTR **(VII)** initially increase, indicating enhanced mitophagy, but later decline as mitochondria are lost. M1 and M2 correlation coefficients **(VIII-IX)** reveal that mitochondria-lysosome interactions peak at 2 hours but decrease by 6 hours, suggesting lysosomal degradation fails to counteract prolonged mitochondrial damage. Statistical significance was determined using a two-tailed paired t-test (*p < 0.05, **p < 0.01, ***p < 0.001). Scale bars = 50 μm.

