## Supplementary Data for "A Hierarchical Cascade of Organellar Silencing and their Regeneration under Anaesthetic Stress in Plants"

### **Supplementary information**

#### **Contents**

##### A. **Methodology**

##### B. **Experimental Results**

##### C. **Statistical Analysis (Main manuscript analysis)**

**A. Methodology**

**1. Selected Plants and Growth Conditions**

The study utilized tomato plants (*Solanum lycopersicum*, F1 hybrid ‘Ashutosh’), sourced locally from Mandi, Himachal Pradesh, selected for their availability and responsiveness to environmental factors. The seeds were sterilized by immersion in 70% ethanol and rinsed thoroughly with sterile distilled water. The sterilized seeds were sown on ½-strength Murashige and Skoog (MS) agar plates. Seedlings were cultivated under 0 hourled conditions at 21°C with a 16-hour light and 8-hour dark photoperiod. Two-week-old seedlings were treated with 1% lidocaine anaesthesia in a time series of 0 hr, 1 hr, 2 hr, 4 hr, 6 hr, and 8 hr. Following anaesthetic exposure, seedlings underwent a recovery process involving three PBS washes and progressively increasing rest periods, with recovery assessed via confocal microscopy.

**2. Prolonged Effect of Anaesthesia on Mitochondria and Mitochondrial Membrane Potential**

To investigate the effect of anaesthesia on mitochondria and membrane potential, root cell mitochondria were stained with MitoTracker Green (Thermo Fisher Scientific) and washed with PBS. Seedlings were treated with 1% lidocaine for up to 8 hours. Mitochondrial membrane potential was visualized using tetramethylrhodamine ethyl ester (TMRE; Thermo Fisher). Recovery was facilitated by washing the samples with PBS three times, with increasing recovery periods.

**3. Autophagy Experiment for Mitochondria**

To study mitochondrial autophagy, lysosomes in root cells were stained with LysoTracker Red (Thermo Fisher Scientific). After staining, root cells were treated with 1% lidocaine over increasing time intervals. The reversibility of the anaesthetic effect was assessed after removing lidocaine and washing with PBS.

**4. ROS Generation Following Anaesthesia Treatment**

Reactive oxygen species (ROS) generation was evaluated in root cells using the H2DCFDA probe (10 mM; Thermo Fisher) after exposure to 1% lidocaine. Recovery was carried out by washing the cells with PBS three times, with progressively increasing recovery periods.

**5. Endocytic Vesicle Trafficking Under Anaesthesia**

Endocytic vesicle trafficking was examined by staining tomato seedling roots with FM4-64 dye (Thermo Fisher Scientific) and exposing them to 1% lidocaine for 0-8 hour. Vesicle recycling was inhibited by treating seedlings with Brefeldin A (BFA; Thermo Fisher) for an additional 30 minutes. Recovery was achieved by removing anaesthesia and washing with PBS three times, with increasing recovery periods.

**6. Nucleus and Chromatin Staining**

Nuclear integrity and chromatin organization were assessed by staining root cell nuclei with DAPI (1 mg/mL; Thermo Fisher) and washing with PBS. After lidocaine treatment for various durations, chromatin arrangement was evaluated via immunofluorescence staining using primary antibodies against H3K4Me3 and H3K9Me3 (1:100; Abclonal) and Cy3-conjugated secondary antibodies (1:600; Abclonal). The samples were mounted with glycerol and sealed before imaging. Recovery involved washing the cells with PBS three times, with increasing rest periods.

**7. Cell Death Staining Following Anaesthesia Exposure**

Cell death was monitored in root cells using propidium iodide staining (0.5 mg/2 mL; Thermo Fisher) after exposure to 1% lidocaine for increasing durations. The extent of cell death was observed using confocal microscopy.

**8. Nuclear factor erythroid 2-related factor 2 (NRF2) Staining Under Anaesthesia Treatment**

NRF2 movement was evaluated via immunofluorescence staining using primary antibodies against NRF2 (1:100; Abclonal) and Cy3-conjugated secondary antibodies (1:600; Abclonal). After lidocaine treatment for various durations, the samples were mounted with glycerol and sealed before imaging.

**9. Confocal and SRRF Microscopy and Image Processing**

**a. Confocal Microscopy**

Confocal imaging was performed using a Nikon Eclipse Ti inverted microscope. Images were acquired with Nikon NIS-Element software. Cell samples were excited with a 405, 561, 639 nm and etc laser, and emissions were collected using appropriate filter sets. Colocalization studies were conducted with a 60x (1.40 NA) oil immersion objective.

**b. SRRF Bioimaging of Heterochromatin and Heterochromatin**

Super-resolution radial fluctuations (SRRF) imaging was used to resolve Heterochromatin and heterochromatin structures. The Nikon Ti Eclipse microscope, equipped with a 100x Plan Apo λ oil immersion objective (NA 1.45) and 1.5x magnification, and an Andor iXon Ultra 897U EMCCD camera, was employed. Acquisition was managed via Andor Solis software, using a 50 ms exposure time, 17 MHz readout rate, and a gain setting of 3. Movies of 5,000 stacked images were saved in. fits format for SRRF processing.

**c. SRRF Image Reconstruction for Chromatin**

SRRF image reconstruction was conducted using the NanoJ-SRRF plugin in ImageJ with optimized parameters (ring radius 0.5, radiality magnification 5, and 6 axes in the ring). Drift correction was performed using NanoJ-core, and images were processed to achieve super-resolution by splitting each pixel into 25 (5x5) sub-pixels. Background artifacts were uniformly removed, and final images were scaled, color-coded, and merged in ImageJ.

**d. Image Processing and Analysis**

ImageJ (National Institutes of Health) was used for processing fluorescence intensity, colocalization, and other quantitative analyses. Custom MATLAB scripts were employed for statistical analysis and image quantification. For intensity measurement we have utilized normalised intensity of the area of 70nm^2 and also, we plot a intensity graph using a line plot of 50nm length.

**10. Statistical Analysis**

Statistical analyses were performed using SPSS Statistics software. Each experimental condition included 10 root replicas. Data were analyzed using paired Student’s *t*-tests. Statistical significance was determined at *p* < 0.05 (**), p < 0.01 (**), and p < 0.001 (****). These significance levels ensured robust evaluation and confidence in the experimental outcomes. Graph was drawn in Originpro software.

##### B. **Experimental Results**

1. **Figure S1** shows mitochondrial recovery after 1 hour of anaesthesia treatment.
2. **Figure S2** shows mitochondrial recovery after 2 hours of anaesthesia treatment at various time points.
3. **Figure S3** shows mitochondrial recovery after 4 hours of anaesthesia treatment at various time points.
4. **Figure S4** shows mitochondrial recovery after 6 hours of anaesthesia treatment at various time points.
5. **Figure S5** shows reactive oxygen species (ROS) levels after 1 hour of anaesthesia treatment at various recovery time points.
6. **Figure S6** shows reactive oxygen species (ROS) levels after 2 hours of anaesthesia treatment at various recovery time points.
7. **Figure S7** shows reactive oxygen species (ROS) levels after 4 hours of anaesthesia treatment at various recovery time points.
8. **Figure S8** shows reactive oxygen species (ROS) levels after 6 hours of anaesthesia treatment at various recovery time points.
9. **Figure S9** shows endocytic vesicle trafficking after 1 hour of anaesthesia treatment at various recovery time points.
10. **Figure S10** shows endocytic vesicle trafficking after 2 hour of anaesthesia treatment at various recovery time points.
11. **Figure S11** shows endocytic vesicle trafficking after 4 hour of anaesthesia treatment at various recovery time points.
12. **Figure S12** shows endocytic vesicle trafficking after 6 hour of anaesthesia treatment at various recovery time points.
13. **Figure S13** shows nuclear changes after 1 hour of anaesthesia treatment at various recovery time points.
14. **Figure S14** shows nuclear changes after 2 hour of anaesthesia treatment at various recovery time points.
15. **Figure S15** shows nuclear changes after 4 hour of anaesthesia treatment at various recovery time points.
16. **Figure S16** shows nuclear changes after 6 & 8 hour of anaesthesia treatment at various recovery time points.
17. **Figure S17** Time-dependent changes in nuclear morphology under general anaesthesia (etomidate) treatment.

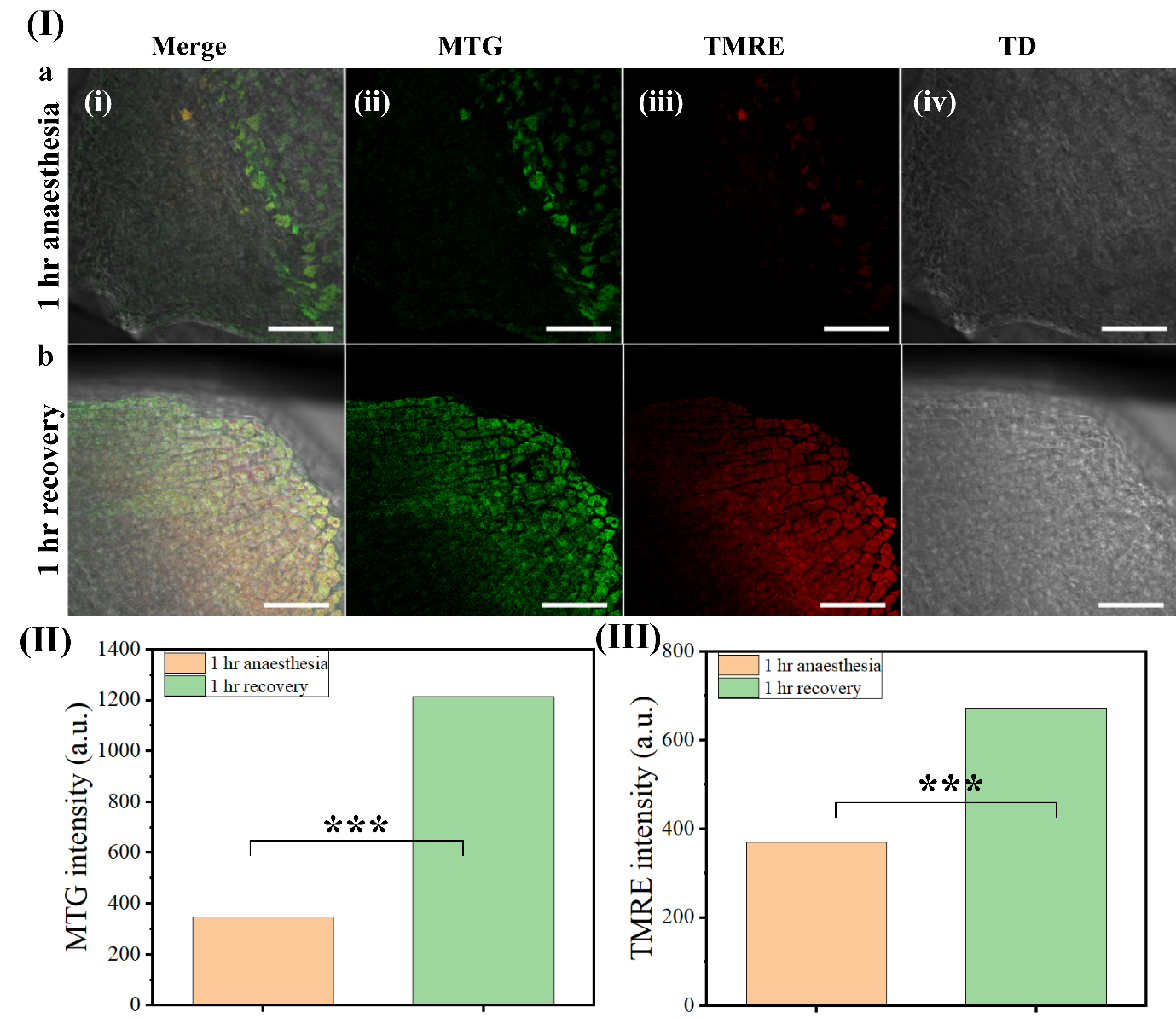
 **Figure S1: Mitochondrial recovery after 1-hour anaesthesia treatment.** **(I)** Representative images compare 1 hour anaesthesia treatment (a) and 1-hour recovery (b) groups. Merged images (i) show MitoTracker Green (MTG, green) and TMRE (red) signals, with individual MTG (ii), TMRE (iii), and Transmission Differential (TD, iv) images providing structural context (scale bar: 50 µm). **(II)** Quantification of MTG intensity shows a significant increase in mitochondrial mass in the recovery group (p < 0.001). **(III)** TMRE intensity indicates a significant rise in mitochondrial membrane potential post-recovery (p < 0.001).

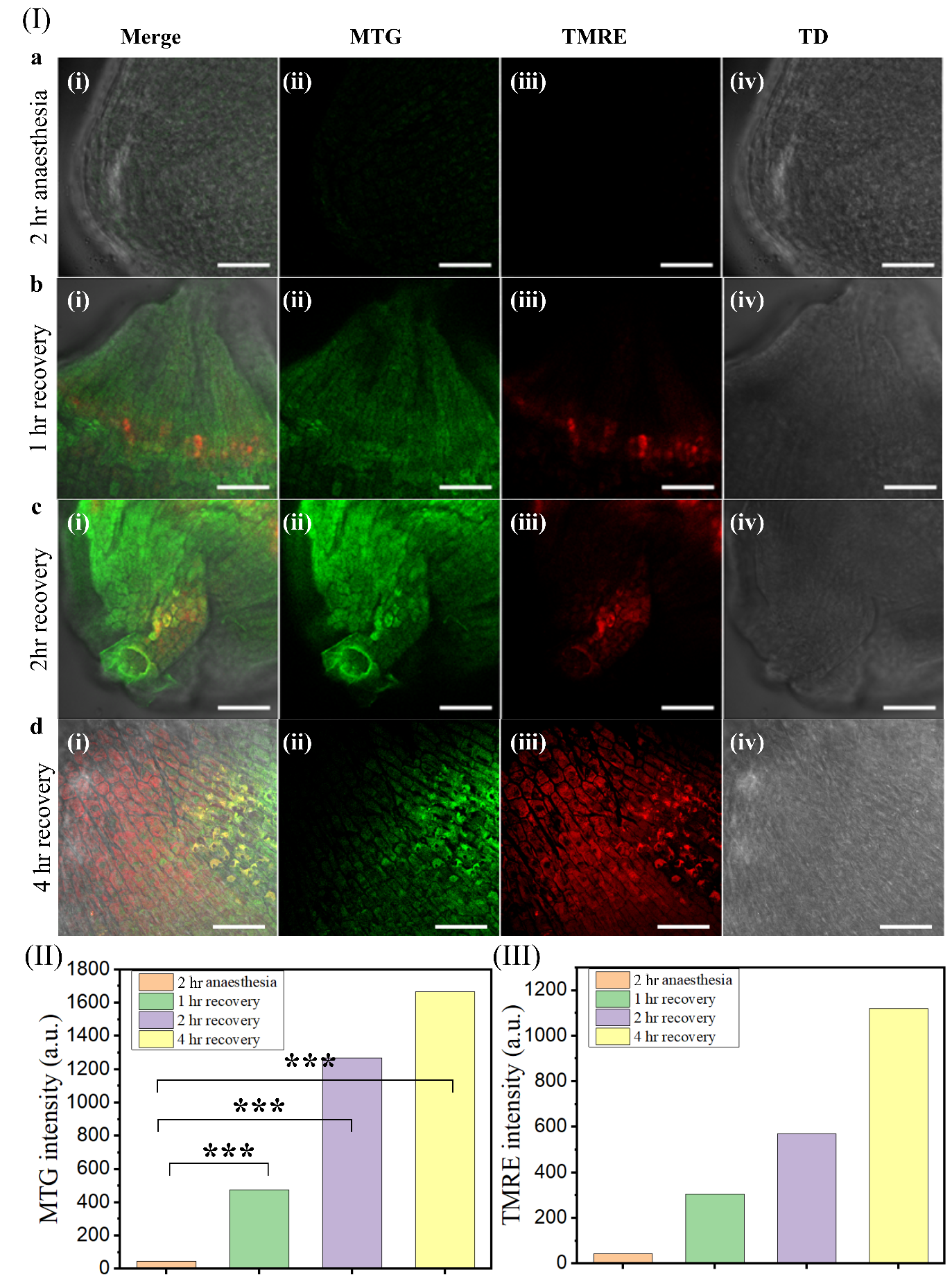

**Figure S2: Mitochondrial recovery after 2-hour anaesthesia treatment at different time points. (I)** Representative images compare 2-hour anaesthesia (a) with 1-hour (b), 2-hour (c), and 4-hour (d) recovery groups. Merged images (i) display MitoTracker Green (MTG, green) and TMRE (red) signals, with MTG (ii), TMRE (iii), and Transmission Differential (TD, iv) images providing structural context (scale bar: 50 µm). **(II)** Quantification of MTG intensity shows a significant increase in mitochondrial mass across recovery groups compared to 0 hour (p < 0.001). **(III)** TMRE intensity analysis reveals a progressive recovery of mitochondrial membrane potential, with a significant increase after 4 hours (p < 0.001).

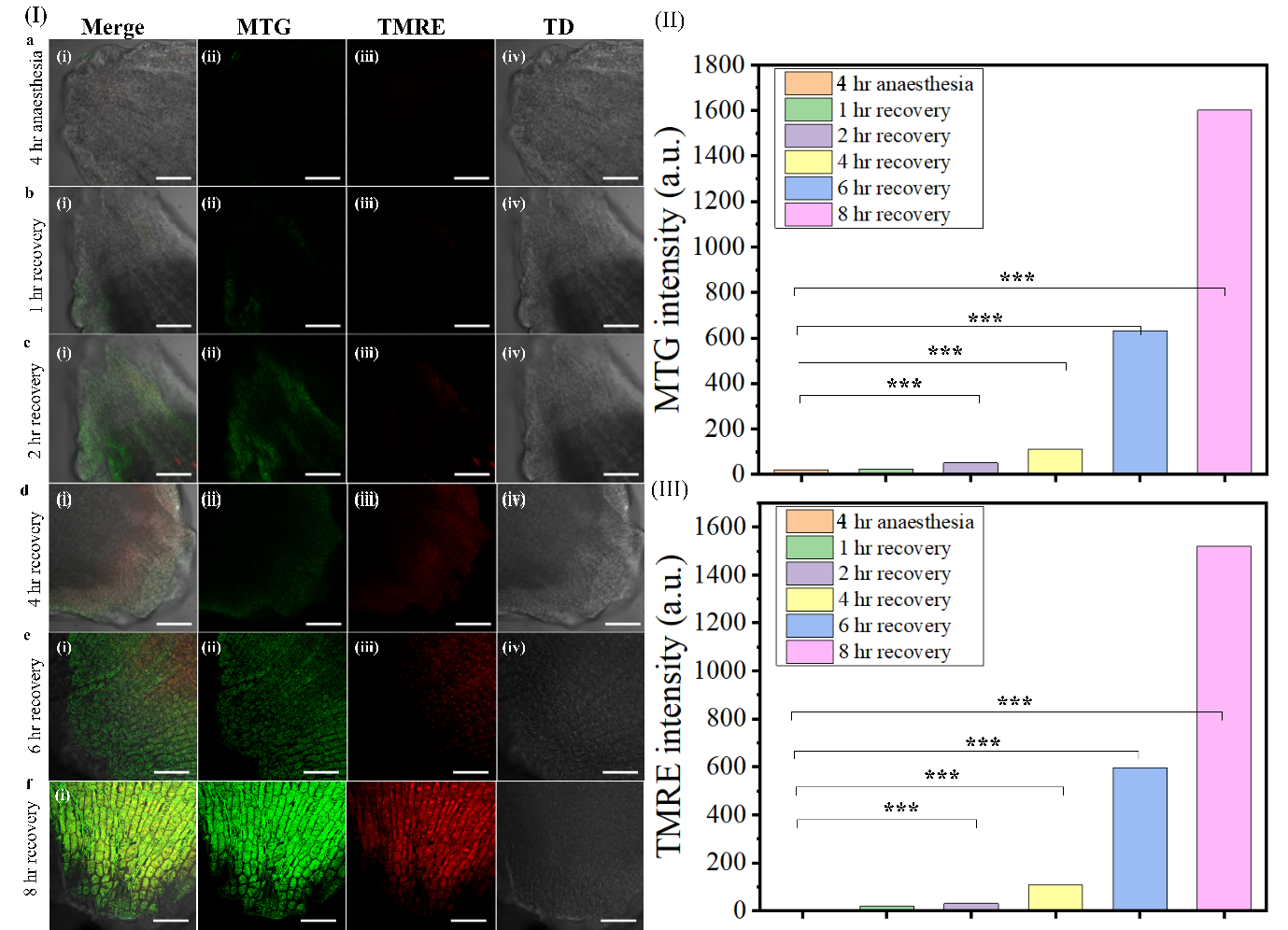

**Figure S3: Mitochondrial recovery after 4-hour anaesthesia treatment at various time points. (I)** Representative images compare 4-hour anaesthesia (a) with 1-hour (b), 2-hour (c), 4-hour (d), 6-hour (e), 8-hour (f) recovery groups. Merged images (i) display MitoTracker Green (MTG, green) and TMRE (red) signals, with MTG (ii), TMRE (iii), and Transmission Differential (TD, iv) images providing structural context (scale bar: 50 µm). **(II)** MTG intensity quantification shows a significant increase in mitochondrial mass across recovery groups, peaking at 8 hours (p < 0.001). **(III)** TMRE intensity analysis indicates a progressive restoration of mitochondrial membrane potential, with the highest recovery at 8 hours (p < 0.001).

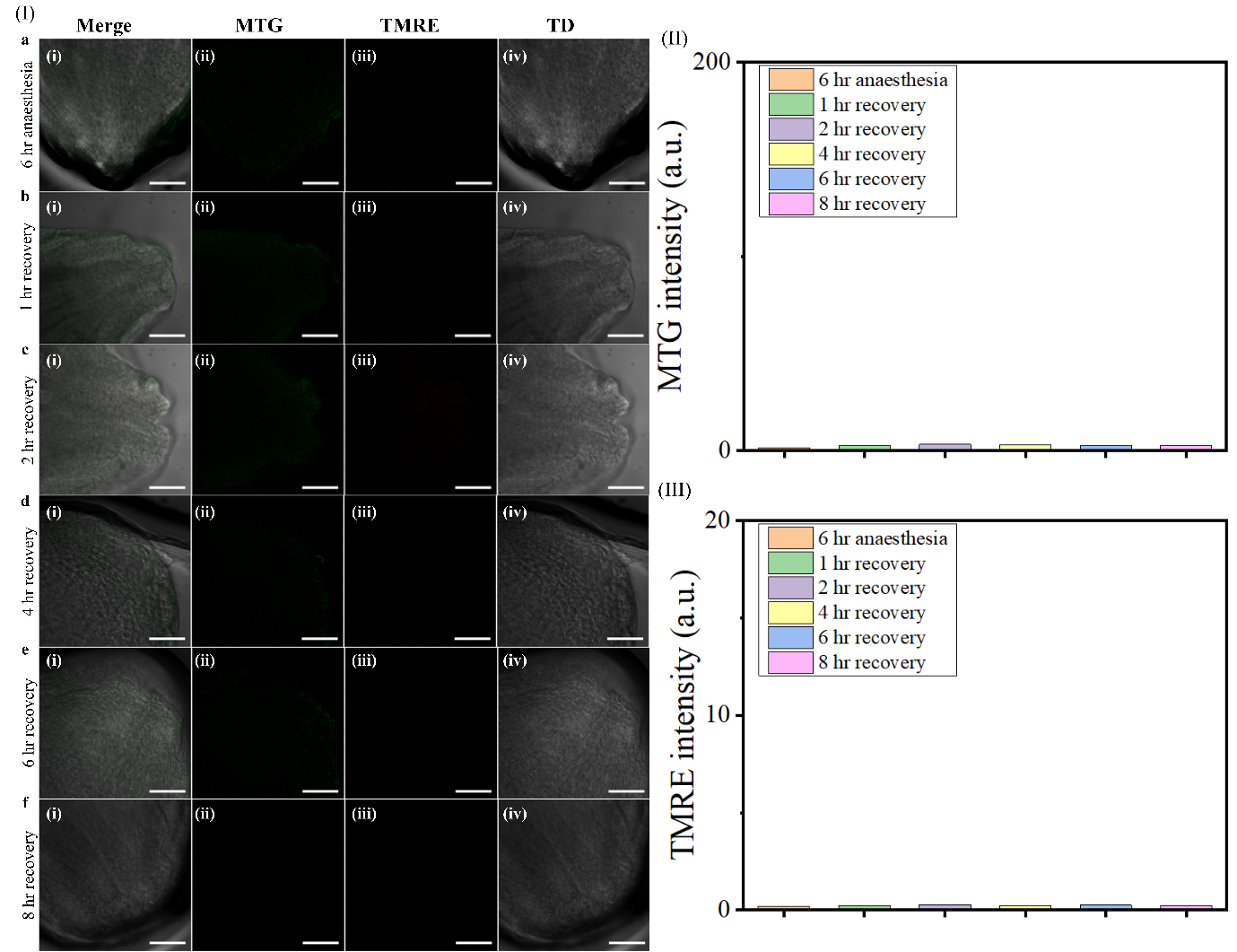

**Figure S4 Mitochondrial recovery after 6-hour anaesthesia treatment at various time points. (I)** Representative images compare 6-hour anaesthesia (a) with 1-hour (b), 2-hour (c), 4-hour (d), 6-hour (e), and 8-hour (f) recovery groups. Merged images (i) display MitoTracker Green (MTG, green) and TMRE (red) signals, with MTG (ii), TMRE (iii), and Transmission Differential (TD, iv) images providing structural context (scale bar: 50 µm). **(II)** MTG intensity quantification shows minimal mitochondrial mass across all groups, with no significant differences compared to the 0 hour. **(III)** TMRE intensity analysis indicates consistently low mitochondrial membrane potential across all recovery groups, with no significant changes from the 0-hour.

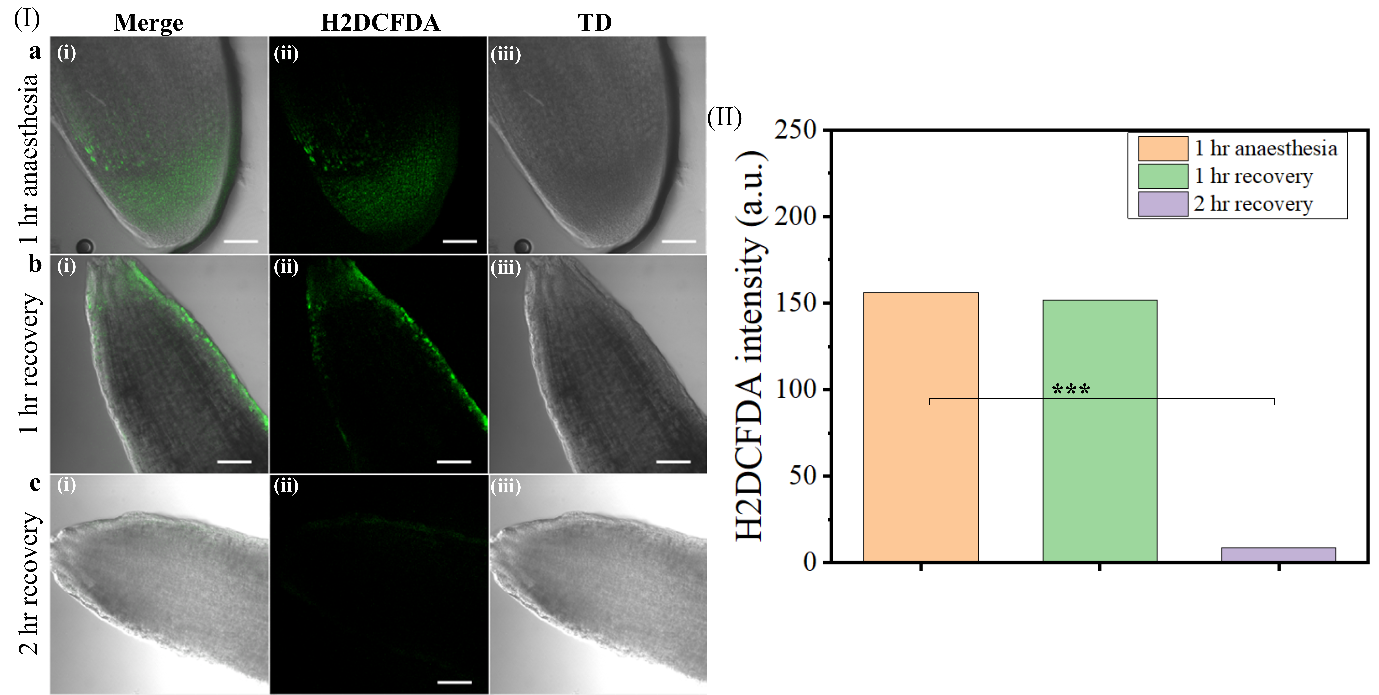

**Figure S5 shows reactive oxygen species (ROS) levels after 1 hour of anaesthesia treatment at various recovery time points.** In panel **(I)**, representative images compare the 1-hour anaesthesia (**a**) with the 1-hour (**b**) and 2-hour (**c**) recovery groups. Each row presents merged images (**i**) showing H2DCFDA staining (green) and root structure, H2DCFDA images (**ii**) highlighting ROS levels (green), and Transmission Differential (TD) images (**iii**) for structural context. Scale bars represent 100µm. **(II)** Quantification of H2DCFDA fluorescence intensity (a.u.) across the control, 1-hour, and 2-hour recovery groups. ROS levels remain high in the control and 1-hour recovery groups, with a significant decrease observed at the 2-hour recovery time point *(****p < 0.001****).*

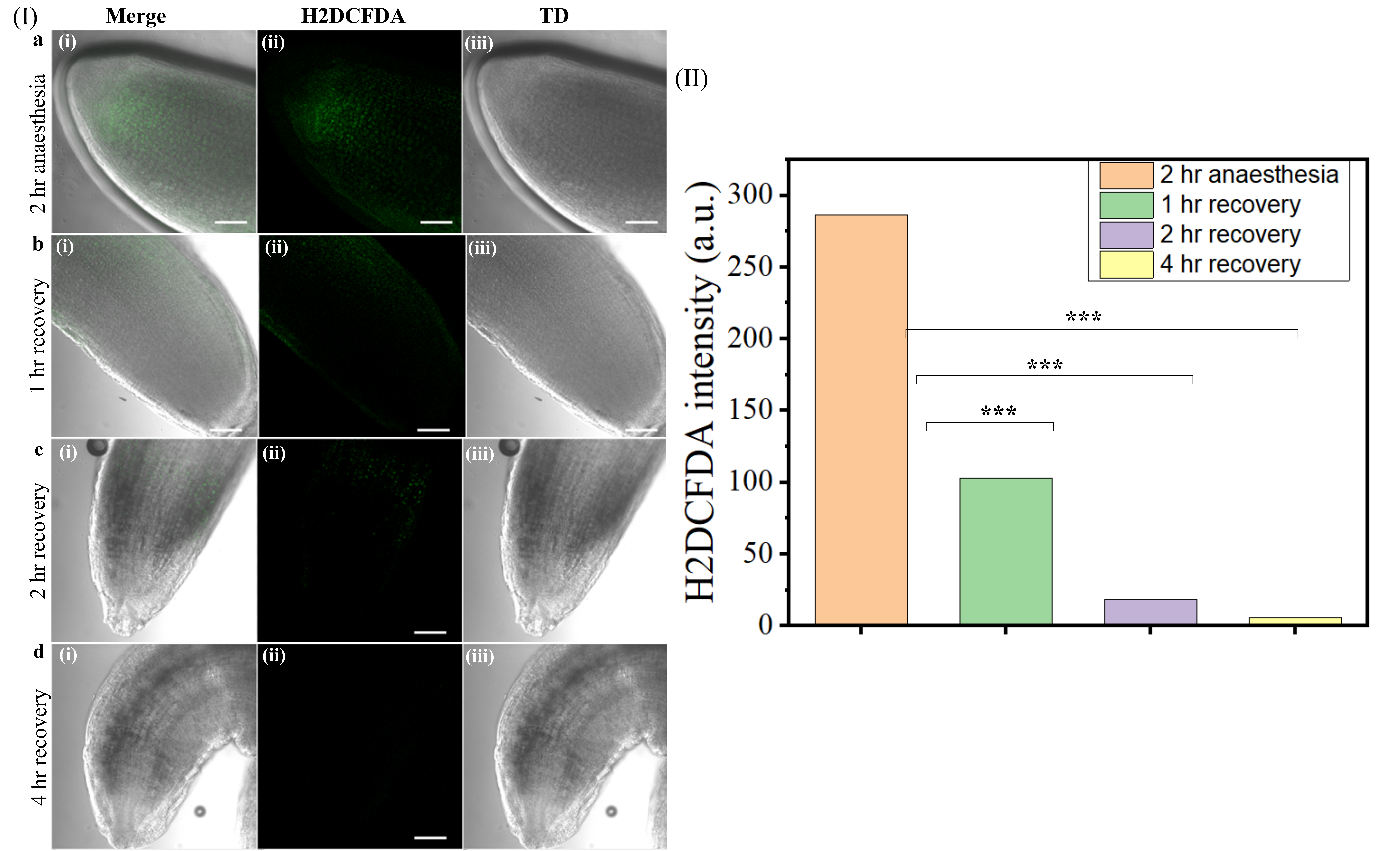

**Figure S6 shows reactive oxygen species (ROS) levels after 2 hours of anaesthesia treatment at various recovery time points.** In panel **(I)**, representative images compare the 2-hour anaesthesia (**a**) with the 1-hour (**b**), 2-hour (**c**), and 4-hour (**d**) recovery groups. Each row presents merged images (**i**) showing H2DCFDA staining (green) and root structure, H2DCFDA images (**ii**) highlighting ROS levels (green), and Transmission Differential (TD) images (**iii**) for structural context. Scale bars represent 100µm. **(II)** Quantification of H2DCFDA fluorescence intensity (a.u.) across the control, 1-hour, 2-hour, and 4-hour recovery groups. ROS levels are significantly reduced with recovery time. The control group shows the highest ROS levels, which decrease notably at the 1-hour recovery mark and continue to drop at 2-hour and 4-hour recovery points *(****p < 0.001****).*

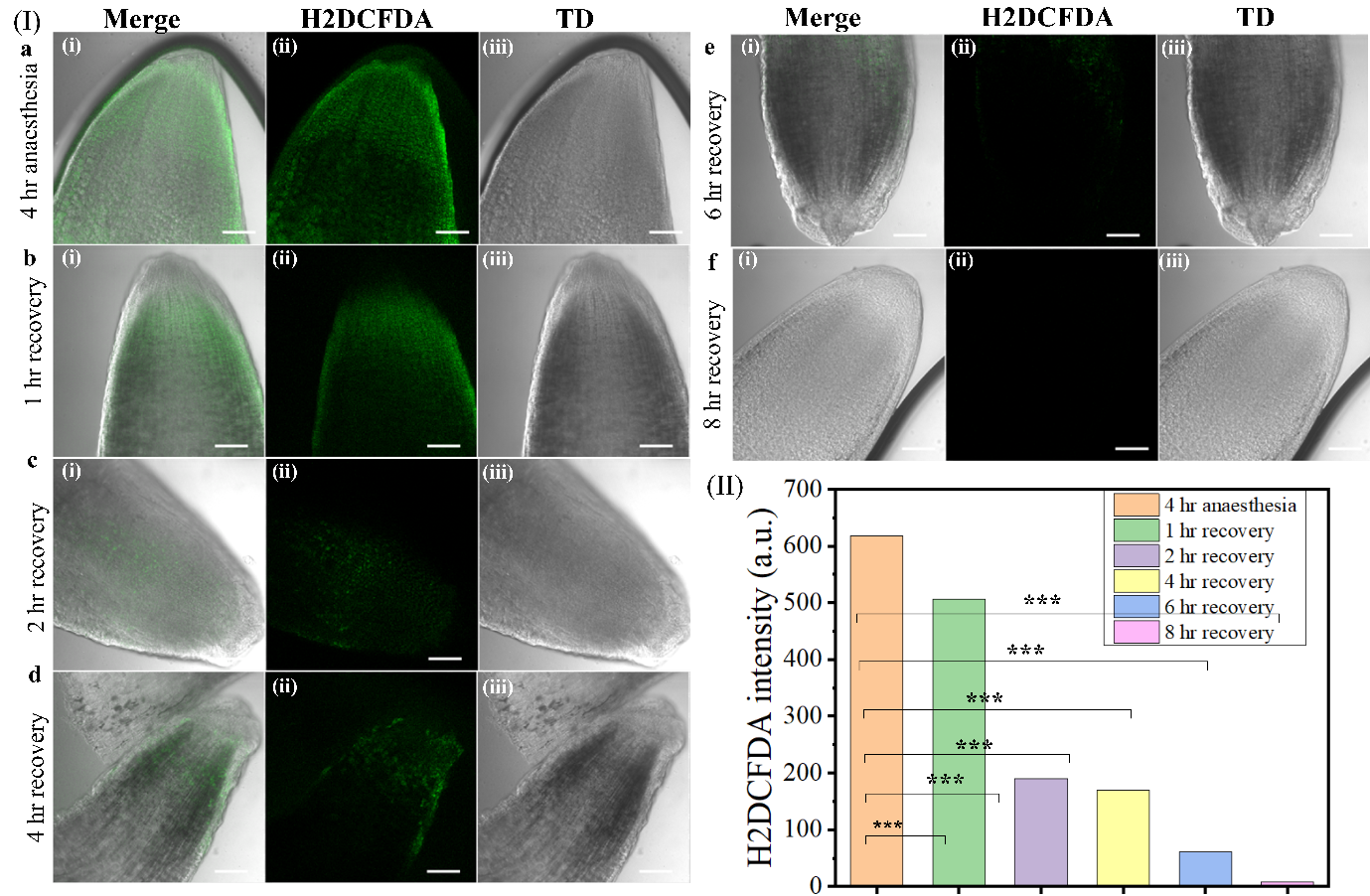

**Figure S7 shows reactive oxygen species (ROS) levels after 4 hours of anaesthesia treatment at various recovery time points.** In panel **(I)**, representative images compare the 4-hour anaesthesia (**a**) with the 1-hour (**b**), 2-hour (**c**), 4-hour (**d**), 6-hour (**e**), 8-hour (**f**) recovery groups. Each row presents merged images (**i**) showing H2DCFDA staining (green) and root structure, H2DCFDA images (**ii**) highlighting ROS levels (green), and Transmission Differential (TD) images (**iii**) for structural context. Scale bars represent 100µm. **(II)** Quantification of H2DCFDA fluorescence intensity (a.u.) across the control, 1- hour to 8-hour recovery groups. ROS levels are significantly elevated in the control and 1-hour recovery groups. However, ROS levels progressively decrease with extended recovery time, showing significant reductions at 2-hour, 4-hour, 6-hour, and 8-hour recovery points *(****p < 0.001****).*

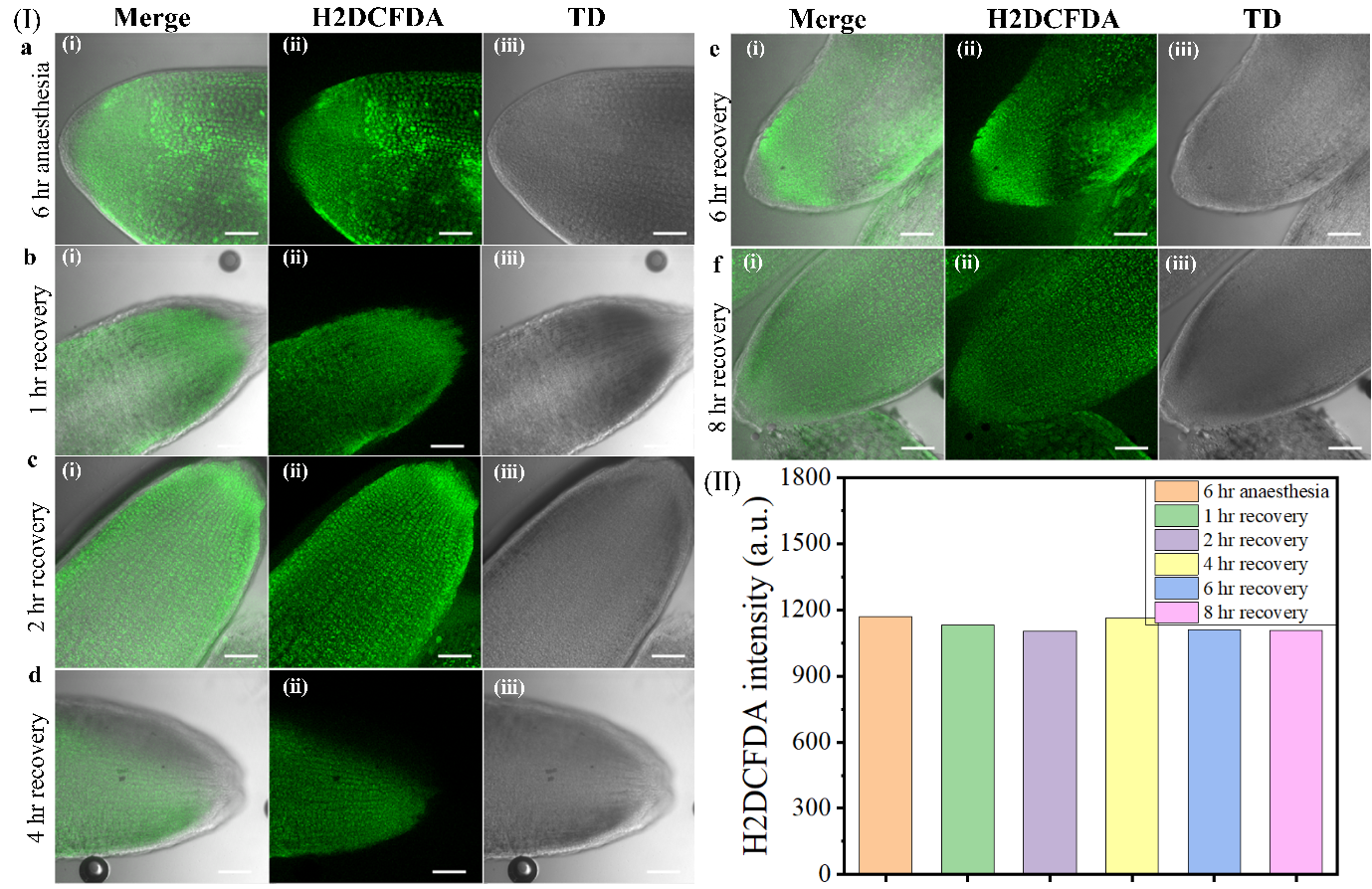

**Figure S8 shows reactive oxygen species (ROS) levels after 6 hours of anaesthesia treatment at various recovery time points.** In panel **(I)**, representative images compare the 6-hour anaesthesia (**a**) with the 1-hour (**b**), 2-hour (**c**), 4-hour (**d**), 6-hour (**e**), and 8-hour (**f**) recovery groups. Each row presents merged images (**i**) showing H2DCFDA staining (green) and root structure, H2DCFDA images (**ii**) highlighting ROS levels (green), and Transmission Differential (TD) images (**iii**) for structural context. **Scale bars** represent 100µm. **(II)** Quantification of H2DCFDA fluorescence intensity (a.u.) across the control, 1-hour, 2-hour, 4-hour, 6-hour, and 8-hour recovery groups. ROS levels remain consistently high across all recovery groups compared to the control, with no significant differences observed. These results suggest that after 6 hours of anaesthesia treatment, ROS levels remain elevated throughout the recovery period, indicating persistent oxidative stress.

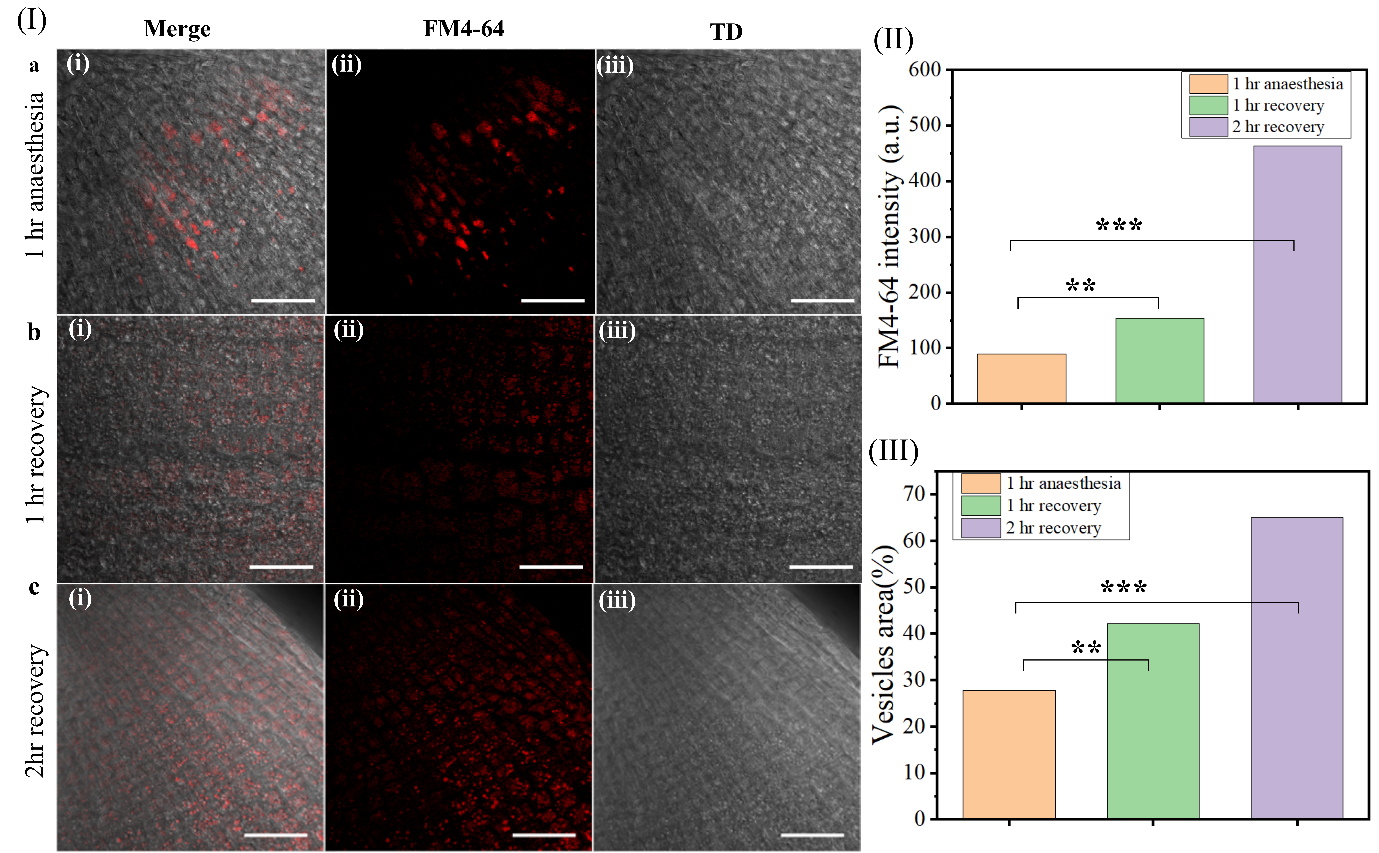

**Figure S9 shows endocytic vesicle trafficking after 1 hour of anaesthesia treatment at various recovery time points.** In panel **(I)**, representative images compare the 1-hour anaesthesia treatment(**a**) with the 1-hour (**b**) and 2-hour (**c**) recovery groups. Each row presents merged images (**i**) showing FM4-64 staining (red) and root structure, FM4-64 images (**ii**) highlighting endocytic vesicle trafficking (red), and Transmission Differential (TD) images (**iii**) for structural context. Scale bars represent 50 µm. **(II)** Quantification of FM4-64 fluorescence intensity (a.u.) across the control, 1-hour, and 2-hour recovery groups. There is a significant increase in FM4-64 intensity with recovery time, with the 2-hour recovery group showing the highest intensity *(****p < 0.001****).* **(III)** Quantification of vesicle area (%) across the control, 1-hour, and 2-hour recovery groups. Vesicle area significantly increases during recovery, with a notable increase observed in the 2-hour recovery group compared to the control *(****p < 0.001****).*

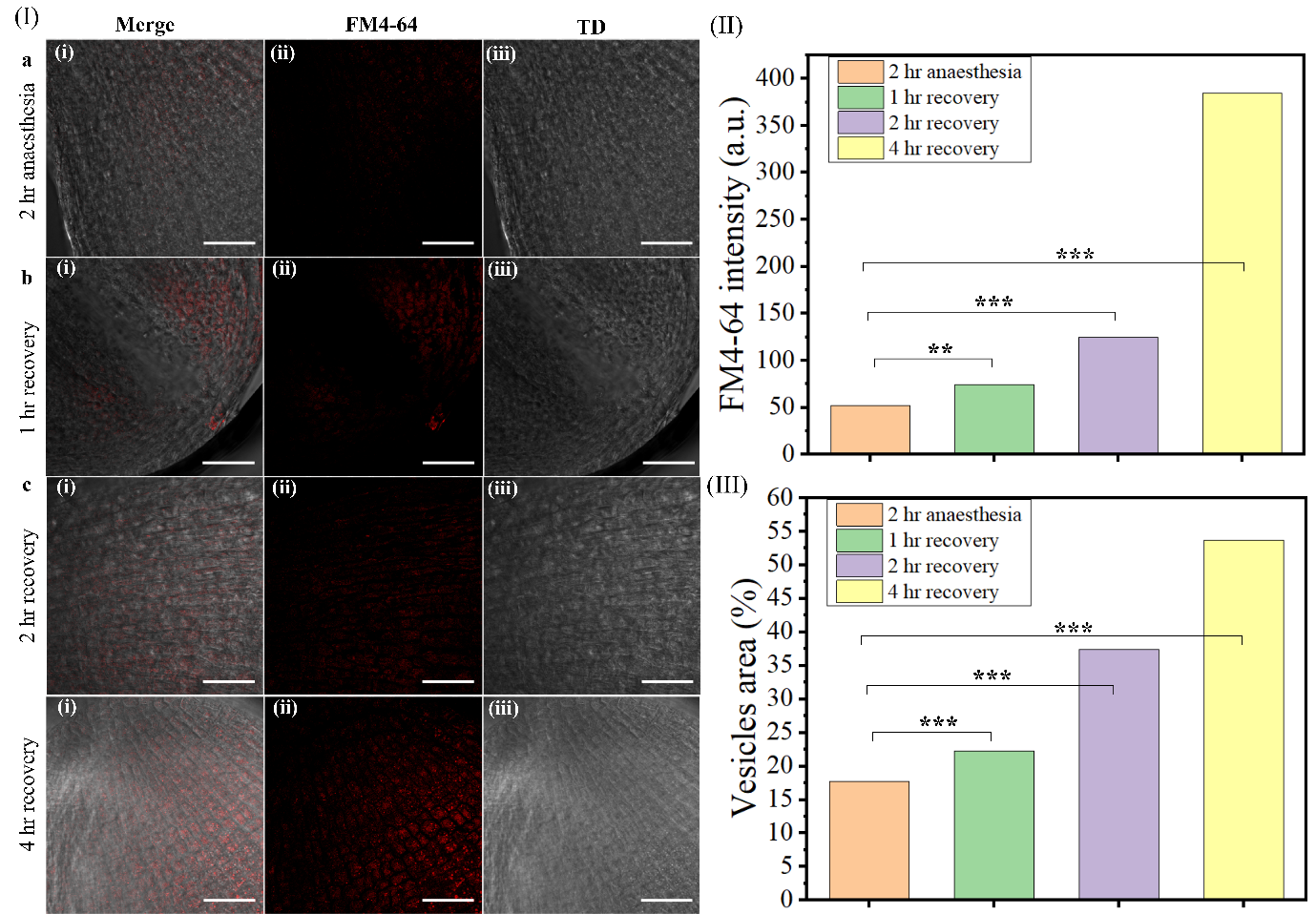

**Figure S10 shows endocytic vesicle trafficking after 2 hours of anaesthesia treatment at various recovery time points.** In panel **(I)**, representative images compare the 2-hour anaesthesia treatment (**a**) with the 1-hour (**b**), 2-hour (**c**), and 4-hour (**d**) recovery groups. Each row presents merged images (**i**) showing FM4-64 staining (red) and root structure, FM4-64 images (**ii**) highlighting endocytic vesicle trafficking (red), and Transmission Differential (TD) images (**iii**) for structural context. Scale bars represent 50 µm. **(II)** Quantification of FM4-64 fluorescence intensity (a.u.) across the control, 1-hour, 2-hour, and 4-hour recovery groups. There is a significant increase in FM4-64 intensity with recovery time, showing a progressive increase, with the highest levels observed at the 4-hour recovery point *(****p < 0.001****).* **(III)** Quantification of vesicle area (%) across the control, 1-hour, 2-hour, and 4-hour recovery groups. Vesicle area significantly increases during recovery, with a notable rise observed at the 2-hour and 4-hour recovery points compared to the control *(****p < 0.001****).*

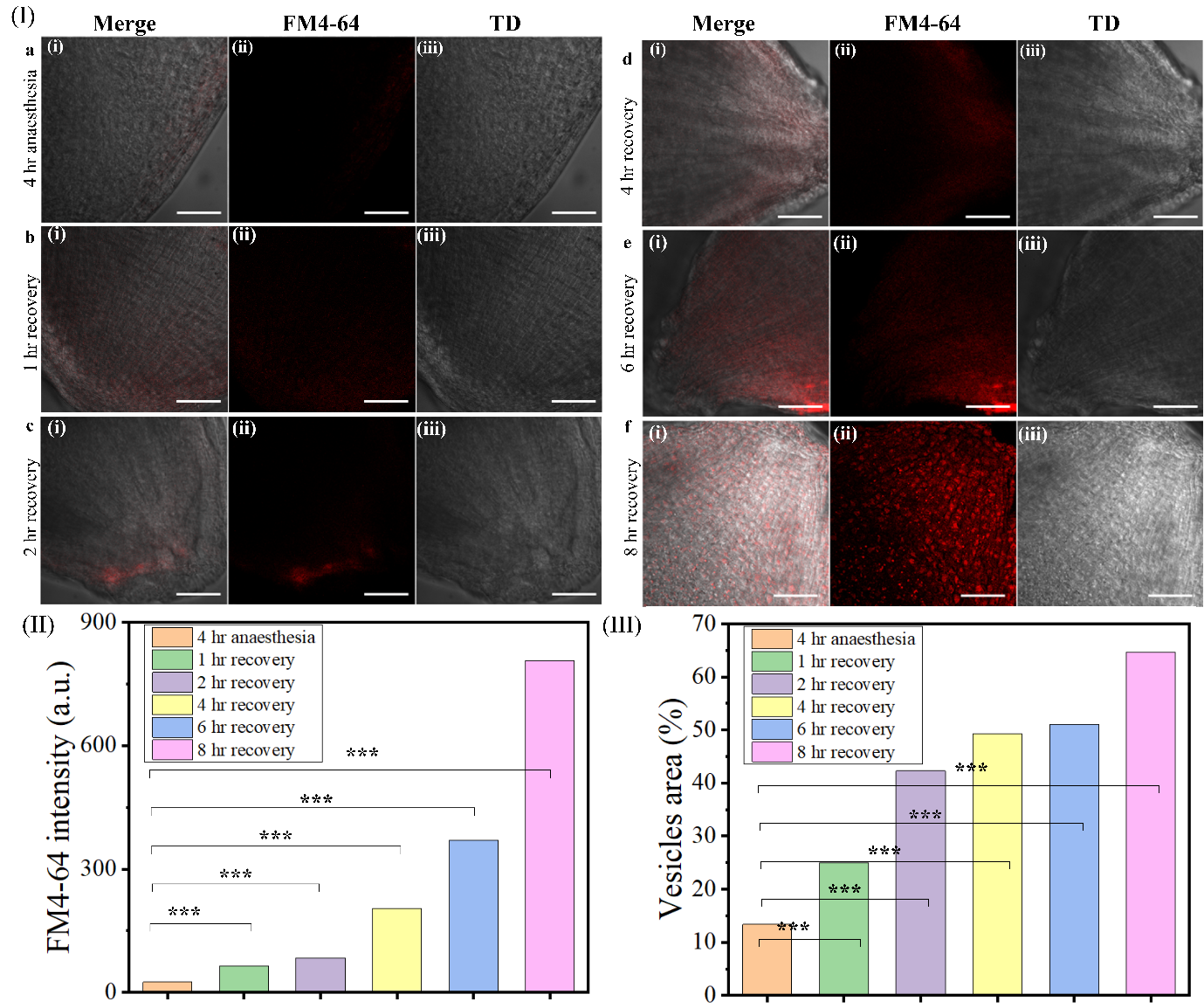

**Figure S11 shows endocytic vesicle trafficking after 4 hours of anaesthesia treatment at various recovery time points.** In panel **(I)**, representative images compare the 4-hour anaesthesia treatment (**a**) with the 1-hour (**b**), 2-hour (**c**), 4-hour (**d**), 6-hour (**e**), 8-hour (**f**) recovery groups. Each row presents merged images (**i**) showing FM4-64 staining (red) and root structure, FM4-64 images (**ii**) highlighting endocytic vesicle trafficking (red), and Transmission Differential (TD) images (**iii**) for structural context. Scale bars represent 50 µm. **(II)** Quantification of FM4-64 fluorescence intensity (a.u.) across the control, 1-hour, 2-hour, 4-hour, 6-hour, and 8-hour recovery groups. There is a significant increase in FM4-64 intensity with recovery time, showing a progressive increase, with the highest levels observed at the 8-hour recovery point *(****p < 0.001****).* **(III)** Quantification of vesicle area (%) across the control, 1-hour, 2-hour, 4-hour, 6-hour, and 8-hour recovery groups. Vesicle area significantly increases during recovery, with substantial increases observed from the 4-hour recovery point onward, reaching the highest levels at 8 hours *(****p < 0.001****).*

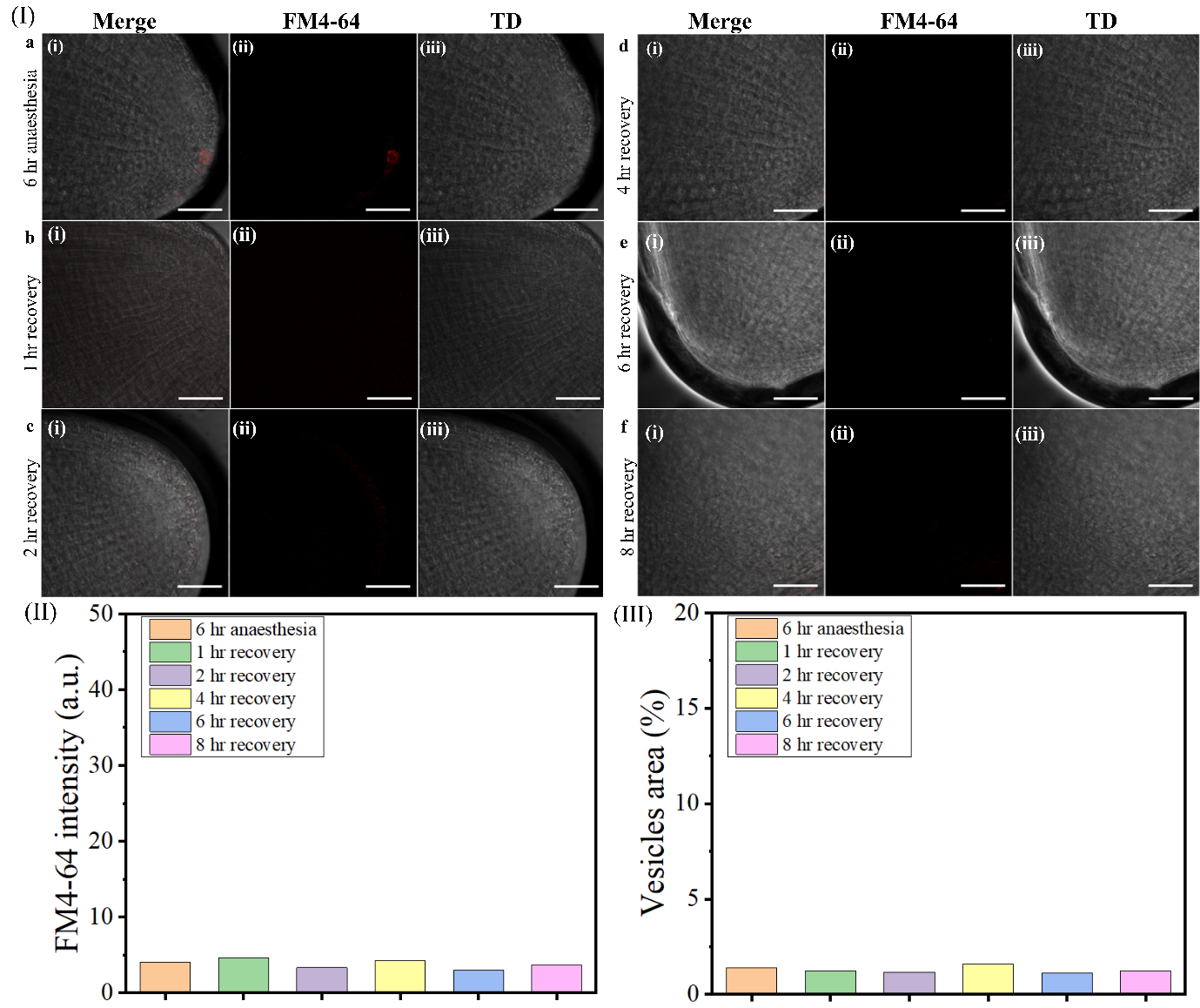

**Figure S12 shows endocytic vesicle trafficking after 6 hours of anaesthesia treatment at various recovery time points.** In panel **(I)**, representative images compare the 6-hour anaesthesia treatment (**a**) with the 1-hour (**b**), 2-hour (**c**), 4-hour (**d**), 6-hour (**e**), and 8-hour (**f**) recovery groups. Each row presents merged images (**i**) showing FM4-64 staining (red) and tissue structure, FM4-64 images (**ii**) highlighting endocytic vesicle trafficking (red), and Transmission Differential (TD) images (**iii**) for structural context. Scale bars represent 50 µm. **(II)** Quantification of FM4-64 fluorescence intensity (a.u.) across the control, 1-hour, 2-hour, 4-hour, 6-hour, and 8-hour recovery groups. FM4-64 intensity remains low across all recovery time points, showing no significant differences compared to the control group. **(III)** Quantification of vesicle area (%) across the control, 1-hour, 2-hour, 4-hour, 6-hour, and 8-hour recovery groups. Vesicle area remains minimal across all recovery time points, with no significant changes compared to the control group. These results indicate that endocytic vesicle trafficking does not recover following 6 hours of anaesthesia treatment, suggesting a prolonged or permanent disruption in vesicle formation and transport within the observed recovery period.

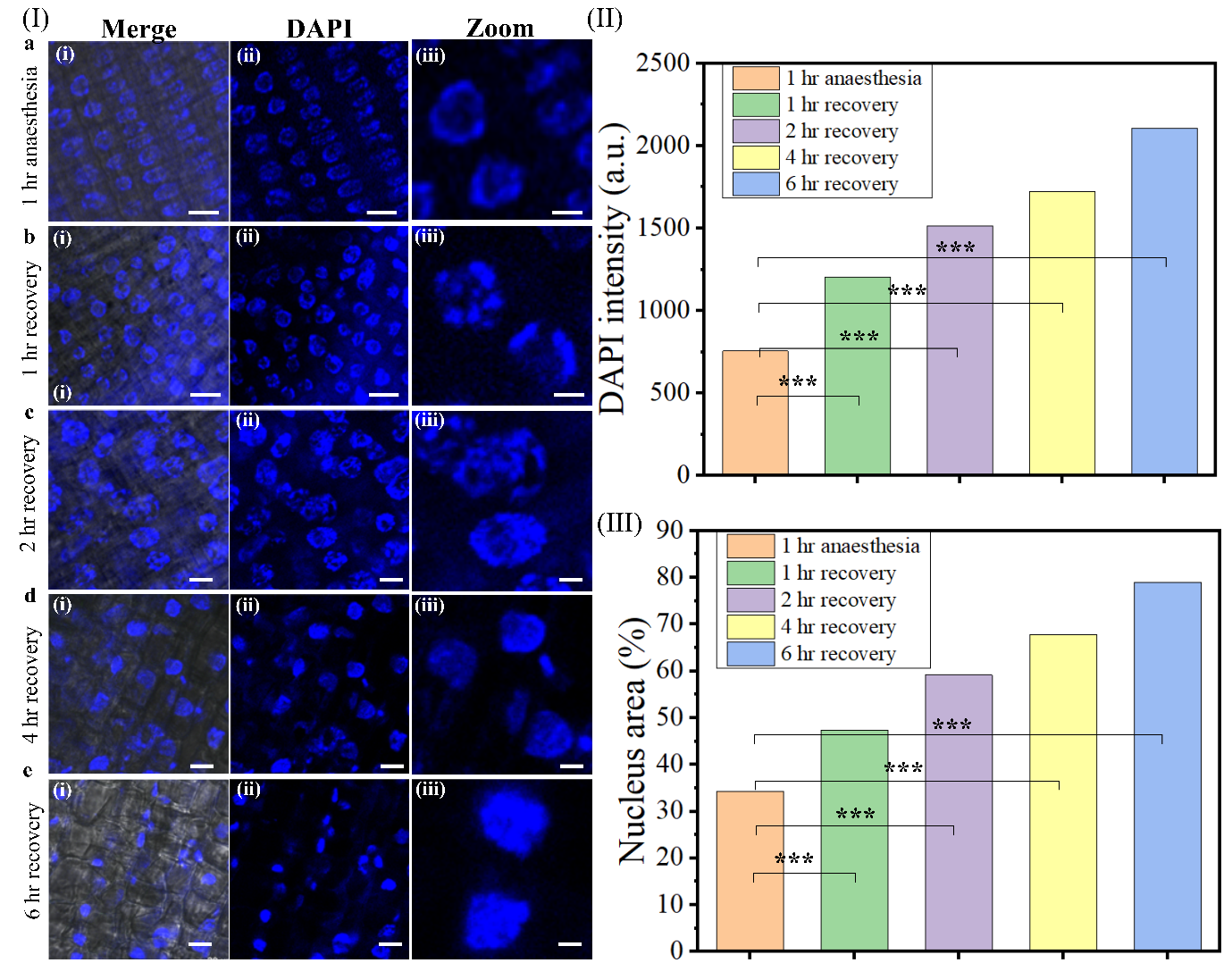

**Figure S13 shows nuclear changes after 1 hour of anaesthesia treatment at various recovery time points.** In panel **(I)**, representative images compare the 1-hour anaesthesia treatment (**a**) with the 1-hour (**b**), 2-hour (**c**), 4-hour (**d**), and 6-hour (**e**) recovery groups. Each row displays merged images (**i**) showing DAPI staining (blue) and tissue structure, DAPI images (**ii**) highlighting nuclear staining, and zoom images (**iii**) for structural context. Scale bars represent 10 µm. Scale bars represent 5 µm. **(II)** Quantification of DAPI fluorescence intensity (a.u.) in the nucleus across the control, 1-hour, 2-hour, 4-hour, and 6-hour recovery groups. DAPI intensity significantly increases with recovery time, indicating nuclear recovery and potential chromatin reorganization after anaesthesia treatment. **(III)** Quantification of nucleus area (%) across the control, 1-hour, 2-hour, 4-hour, and 6-hour recovery groups. Nuclear area progressively increases with recovery time, showing a significant difference between the control and 6-hour recovery group. The graph quantifies this increase, showing statistically significant differences *(****p < 0.001****)* at each time point.

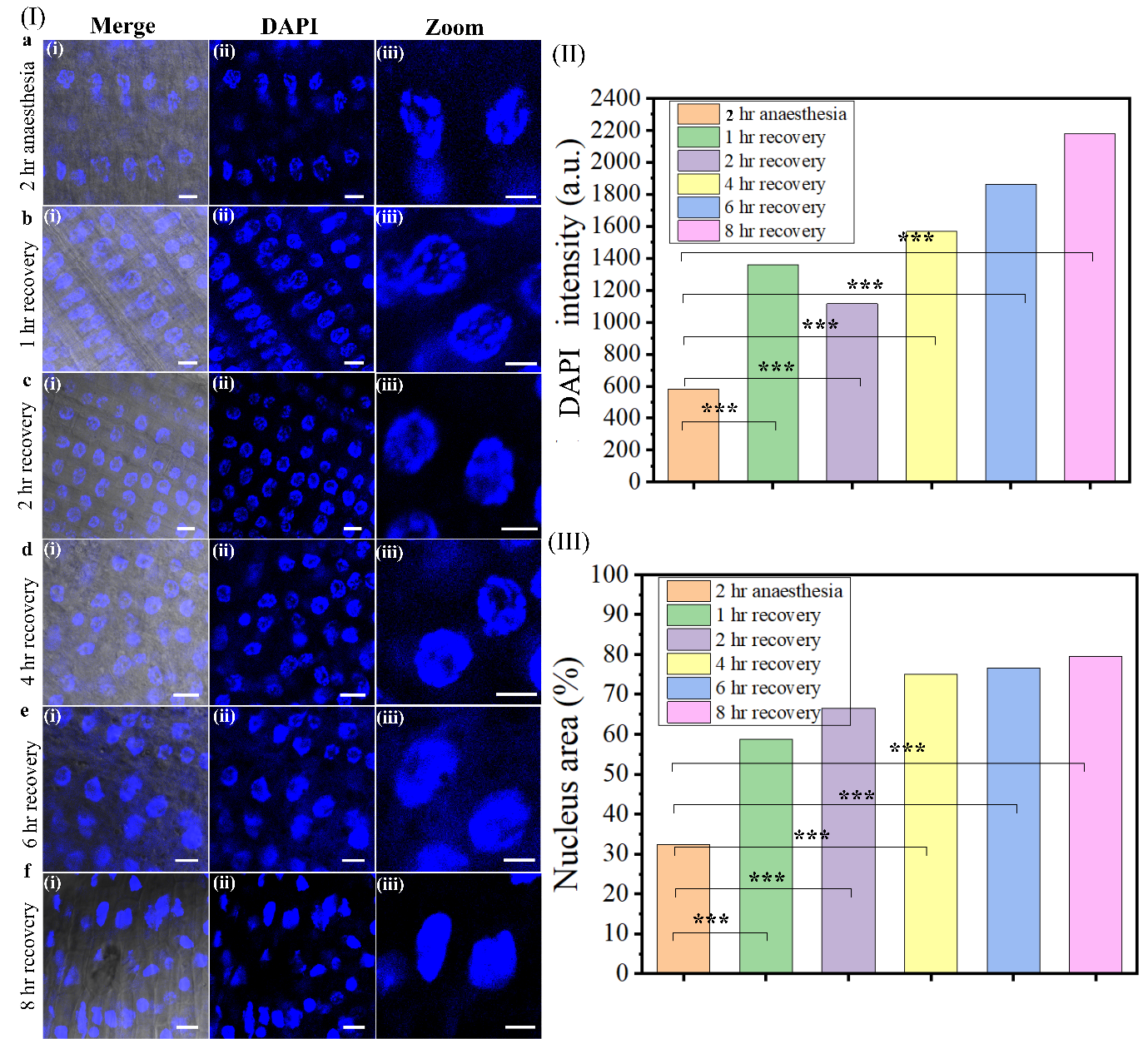

**Figure S14 shows nuclear changes after 2 hours of anaesthesia treatment at various recovery time points.** In panel **(I)**, representative images compare the 2-hour anaesthesia treatment (**a**) with the 1-hour (**b**), 2-hour (**c**), 4-hour (**d**), 6-hour (**e**), and 8-hour (**f**) recovery groups. Each row displays merged images (**i**) showing DAPI staining (blue) and tissue structure, DAPI images (**ii**) highlighting nuclear staining and zoom images (**iii**) for structural context. Scale bars represent 10 µm. Scale bars represent 5 µm. **(II)** Quantification of DAPI fluorescence intensity (a.u.) in the nucleus across the control, 1-hour, 2-hour, 4-hour, 6-hour and 8-hour recovery groups. DAPI intensity significantly increases with recovery time, indicating nuclear recovery and potential chromatin reorganization after anaesthesia treatment. **(III)** Quantification of nucleus area (%) across the control, 1-hour, 2-hour, 4-hour, 6-hour and 8-hour recovery groups. Nuclear area progressively increases with recovery time, showing a significant difference between the control and 6-hour recovery group\The graph quantifies this increase, showing statistically significant differences *(****p < 0.001****)* at each time point.

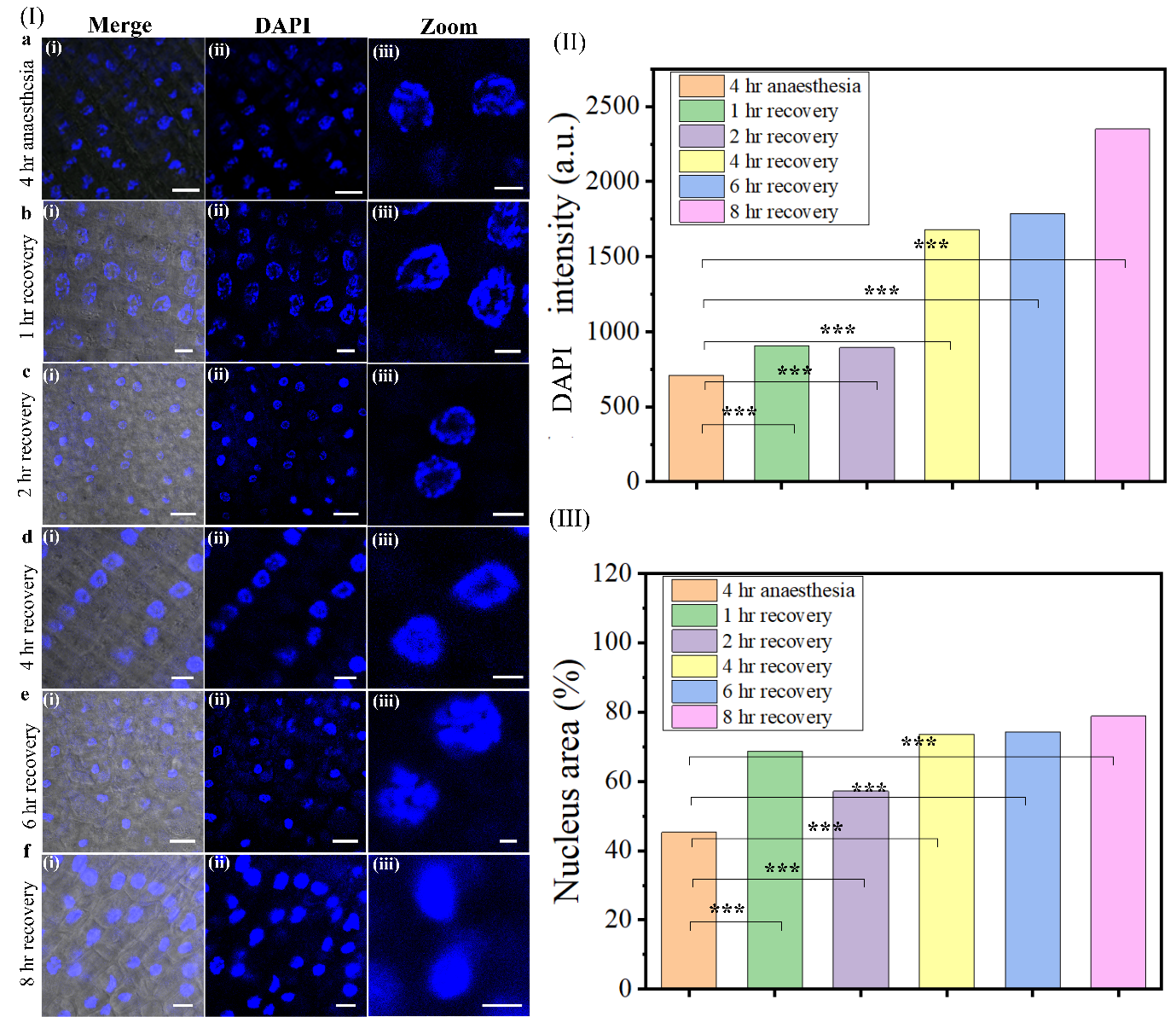

**Figure S15 shows nuclear changes after 4 hours of anaesthesia treatment at various recovery time points.** In panel **(I)**, representative images compare the 4-hour anaesthesia treatment (**a**) with the 1-hour (**b**), 2-hour (**c**), 4-hour (**d**), 6-hour (**e**), and 8-hour (**f**) recovery groups. Each row displays merged images (**i**) showing DAPI staining (blue) and tissue structure, DAPI images (**ii**) highlighting nuclear staining, and zoom images (**iii**) for structural context. Scale bars represent 10 µm. **(II)** Quantification of DAPI fluorescence intensity (a.u.) in the nucleus across the control, 1-hour, 2-hour, 4-hour, 6-hour, and 8-hour recovery groups. DAPI intensity significantly increases with recovery time, indicating nuclear recovery and potential chromatin reorganization after anaesthesia treatment. µm **(III)** Quantification of nucleus area (%) across the control, 1-hour, 2-hour, 4-hour, 6-hour, and 8-hour recovery groups. The nuclear area increases progressively with recovery time, showing a significant difference between the control and 8-hour recovery group. The graph shows statistically significant differences *(****p < 0.001****)* at each recovery time point.

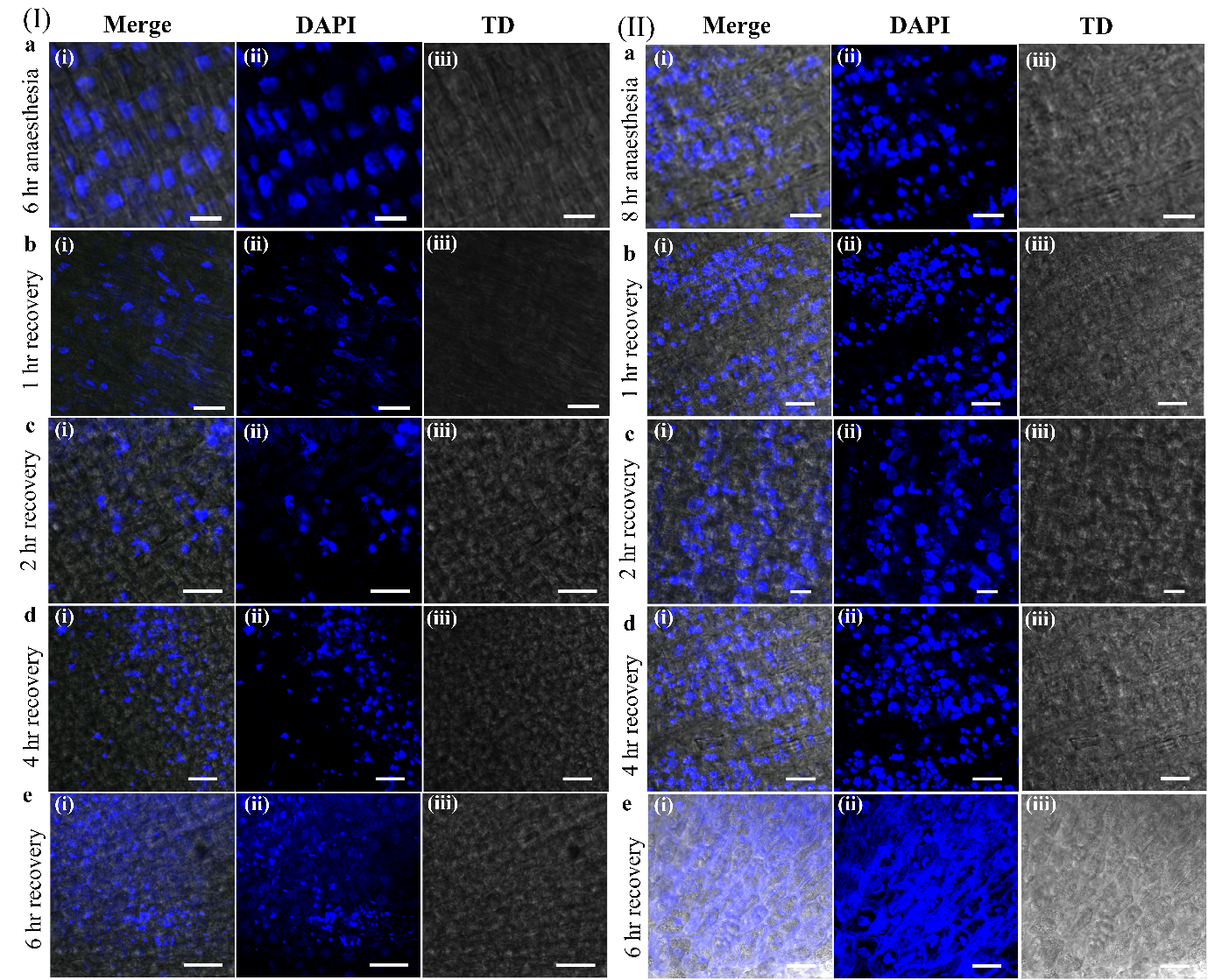

**Figure S16 shows nuclear changes after 6-hour and 8-hour anaesthesia treatments, demonstrating nuclear breakdown with no recovery.** In panel **(I)**, representative images show the effects of 6 to 8-hour anaesthesia treatment and subsequent recovery time points. The 1-hour anaesthesia treatment (**a**) is compared to the 1-hour (**b**), 2-hour (**c**), 4-hour (**d**), and 6-hour (**e**) recovery groups. Each row presents merged images (**i**) showing DAPI staining (blue) and tissue structure, DAPI images (**ii**) highlighting nuclear staining, and Transmission Differential (TD) images (**iii**) for structural context. Scale bars represent 10 µm. In panel **(II)**, representative images show the effects of 8-hour anaesthesia treatment and subsequent recovery time points. The control group (**a**) is compared to the 1-hour (**b**), 2-hour (**c**), 4-hour (**d**), and 6-hour (**e**) recovery groups. Each row displays merged images (**i**), DAPI images (**ii**), and TD images (**iii**). Scale bars represent 10 µm. The images in both panels reveal significant nuclear breakdown with no visible recovery, as indicated by fragmented or diffuse DAPI staining and disrupted nuclear morphology. These results suggest that prolonged anaesthesia treatment (6 hours and 8 hours) leads to severe nuclear damage that is not reversible within the observed recovery periods.

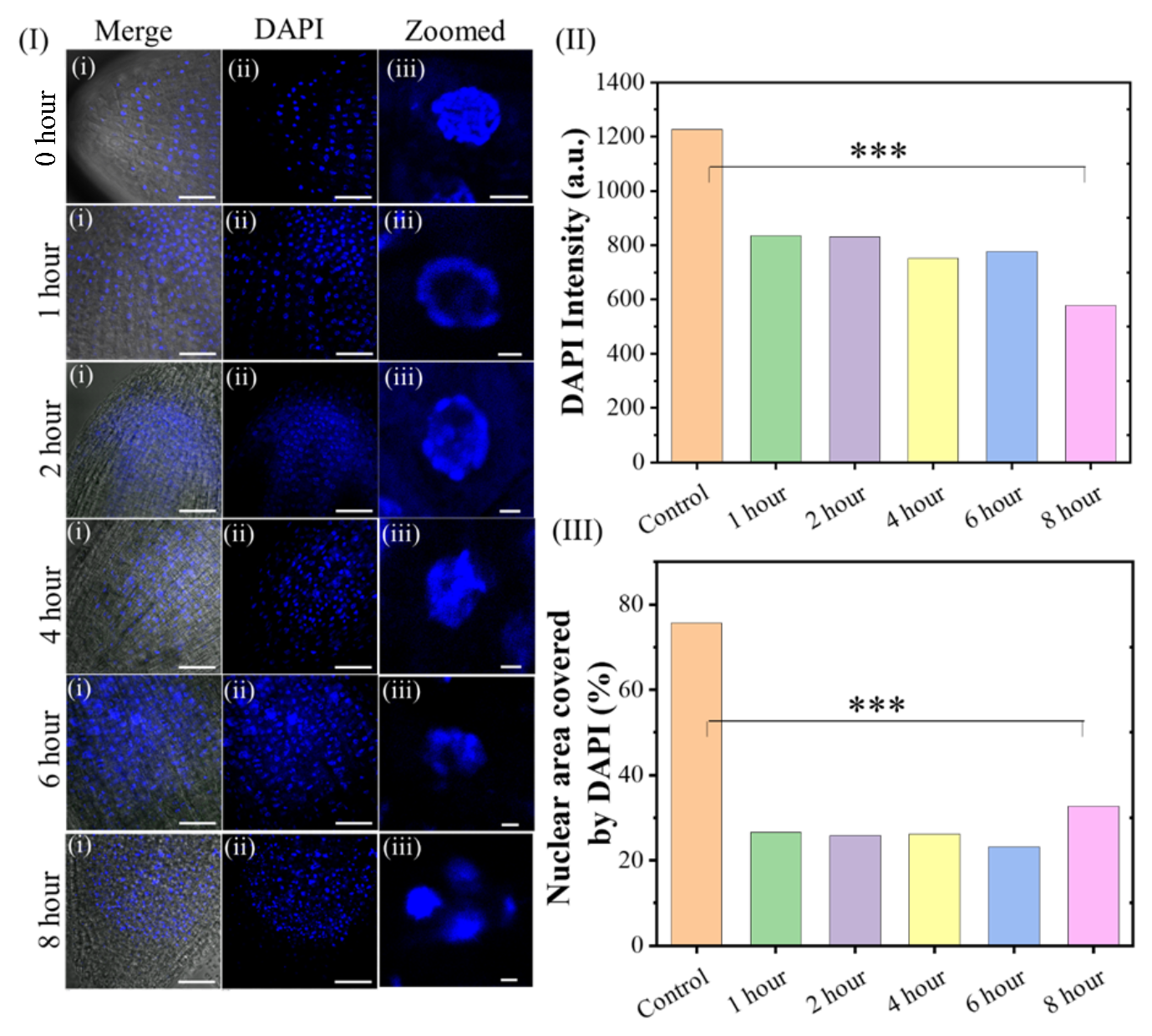

**Figure S17** **(I)** Time-dependent changes in nuclear morphology under general anaesthesia (etomidate) treatment. (i–iii) DAPI-stained root apex cells show progressive chromatin decondensation and nuclear disorganization from 1 h to 8 h. Right: **(II-III)** Quantification of DAPI intensity (top) and percentage of intact nuclei (bottom) confirms nuclear disruption. Similar patterns were observed with etomidate, indicating conserved effects of both anaesthetics on nuclear architecture. The graph shows statistically significant differences *(****p < 0.001****)* at each time point.

##### C. **Statistical Analysis (Main manuscript analysis)**

1. **Table 1:** Mean, standard deviation, and two-tailed paired t-test analysis of MitoTracker Green (MTG) intensity values measured at various time points during anaesthesia.
2. **Table 2:** Mean, standard deviation, and two-tailed paired t-test analysis of TMRE intensity values measured at various time points during anaesthesia.
3. **Table 3:** Mean, standard deviation, and two-tailed paired t-test analysis of MTG and LTR Pearson’s Correlation Coefficient (PCC) values measured at various time points during anaesthesia.
4. **Table 4:** Mean, standard deviation, and two-tailed paired t-test analysis of MTG and LTR Mander’s Coefficient 1 (M1) values measured at various time points during anaesthesia.
5. **Table 5:** Mean, standard deviation, and two-tailed paired t-test analysis of MTG and LTR Mander’s Coefficient 2 (M2) values measured at various time points during anaesthesia.
6. **Table 6:** Mean, standard deviation, and two-tailed paired t-test analysis of ROS generation using H2DCFDA probe intensity values measured at various time points during anaesthesia.
7. **Table 7:** Mean, standard deviation, and two-tailed paired t-test analysis of vesicle generation using FM4-64 probe intensity values measured at various time points during anaesthesia.
8. **Table 8:** Mean, standard deviation, and two-tailed paired t-test analysis of vesicle area in plants using FM4-64 probe values measured at various time points during anaesthesia.
9. **Table 9:** Mean, standard deviation, and two-tailed paired t-test analysis of NRF2 and DAPI Pearson’s Correlation Coefficient (PCC) values measured at various time points during anaesthesia.
10. **Table 10:** Mean, standard deviation, and two-tailed paired t-test analysis of NRF2 and DAPI Mander’s Coefficient 1 (M1) values measured at various time points during anaesthesia.
11. **Table 11:** Mean, standard deviation, and two-tailed paired t-test analysis of NRF2 and DAPI Mander’s Coefficient 2 (M2) values measured at various time points during anaesthesia.
12. **Table 12:** Mean, standard deviation, and two-tailed paired t-test analysis of DAPI intensity values measured at various time points during anaesthesia.
13. **Table 13:** Mean, standard deviation, and two-tailed paired t-test analysis of DAPI area (%) values measured at various time points during anaesthesia.
14. **Table 14:** Mean, standard deviation, and two-tailed paired t-test analysis of DAPI and PI Pearson’s Correlation Coefficient (PCC) values measured at various time points during anaesthesia.
15. **Table 15:** Mean, standard deviation, and two-tailed paired t-test analysis of DAPI and PI Mander’s Coefficient 1 (M1) values measured at various time points during anaesthesia.
16. **Table 16:** Mean, standard deviation, and two-tailed paired t-test analysis of DAPI and PI Mander’s Coefficient 2 (M2) values measured at various time points during anaesthesia.
17. **Table 17:** Mean, standard deviation, and two-tailed paired t-test analysis of PI-positive cell values measured at various time points during anaesthesia.
18. **Table 18:** Mean, standard deviation, and two-tailed paired t-test analysis of PI intensity values measured at various time points during anaesthesia.
19. **Table 19:** Mean, standard deviation, and two-tailed paired t-test analysis of euchromatin (H3K4me3) domain number values measured at various time points during anaesthesia.
20. **Table 20:** Mean, standard deviation, and two-tailed paired t-test analysis of euchromatin (H3K4me3) percentage area values measured at various time points during anaesthesia.
21. **Table 21:** Mean, standard deviation, and two-tailed paired t-test analysis of euchromatin (H3K4me3) mean domain area values measured at various time points during anaesthesia.
22. **Table 22:** Mean, standard deviation, and two-tailed paired t-test analysis of euchromatin (H3K4me3) mean domain perimeter values measured at various time points during anaesthesia.
23. **Table 23:** Mean, standard deviation, and two-tailed paired t-test analysis of euchromatin (H3K4me3) mean circularity values measured at various time points during anaesthesia.
24. **Table 24:** Mean, standard deviation, and two-tailed paired t-test analysis of euchromatin (H3K4me3) mean major axis length values measured at various time points during anaesthesia.
25. **Table 25:** Mean, standard deviation, and two-tailed paired t-test analysis of euchromatin (H3K4me3) mean minor axis length values measured at various time points during anaesthesia.
26. **Table 26:** Mean, standard deviation, and two-tailed paired t-test analysis of euchromatin (H3K4me3) distance from center to periphery values measured at various time points during anaesthesia.
27. **Table 27:** Mean, standard deviation, and two-tailed paired t-test analysis of heterochromatin (H3K9me3) domain number values measured at various time points during anaesthesia.
28. **Table 28:** Mean, standard deviation, and two-tailed paired t-test analysis of heterochromatin (H3K9me3) percentage area values measured at various time points during anaesthesia.
29. **Table 29:** Mean, standard deviation, and two-tailed paired t-test analysis of heterochromatin (H3K9me3) mean domain area values measured at various time points during anaesthesia.
30. **Table 30:** Mean, standard deviation, and two-tailed paired t-test analysis of heterochromatin (H3K9me3) mean domain perimeter values measured at various time points during anaesthesia.
31. **Table 31:** Mean, standard deviation, and two-tailed paired t-test analysis of heterochromatin (H3K9me3) mean circularity values measured at various time points during anaesthesia.
32. **Table 32:** Mean, standard deviation, and two-tailed paired t-test analysis of heterochromatin (H3K9me3) mean major axis length values measured at various time points during anaesthesia.
33. **Table 33:** Mean, standard deviation, and two-tailed paired t-test analysis of heterochromatin (H3K9me3) mean minor axis length values measured at various time points during anaesthesia.
34. **Table 34:** Mean, standard deviation, and two-tailed paired t-test analysis of heterochromatin (H3K9me3) distance from center to periphery values measured at various time points during anaesthesia.

**Table 1: This table presents the mean, standard deviation and statistical analysis of MTG intensity values measured at various time points during anaesthesia (n=10)**

| S.No. | Control | Treatments (MTG) | | | | |
| --- | --- | --- | --- | --- | --- | --- |
|  |  | **1 hour** | **2 hour** | **4 hour** | **6 hour** | **8 hour** |
| 1 | 2386.85733 | 297.841 | 129.12569 | 29.84911 | 19.19957 | 0.9 |
| 2 | 1819.42217 | 367.451 | 92.61721 | 13.03577 | 7.18426 | 0.07 |
| 3 | 1865.98967 | 320.3407 | 131.98497 | 95.29197 | 7.81128 | 0 |
| 4 | 2091.62603 | 282.1042 | 124.42149 | 25.49906 | 7.65767 | 0.01 |
| 5 | 1964.55162 | 382.0213 | 100.9453 | 22.32028 | 28.24769 | 0 |
| 6 | 2056.66872 | 289.2101 | 290.77411 | 122.36425 | 8.10403 | 0.06 |
| 7 | 2343.22906 | 347.6427 | 114.42802 | 25.60977 | 7.26441 | 0 |
| 8 | 2226.34982 | 337.3589 | 110.259 | 75.99674 | 8.2882 | 0.3 |
| 9 | 1991.74323 | 343.3454 | 109.88982 | 22.97968 | 18.11163 | 0 |
| 10 | 1680.95509 | 395.6148 | 129.83785 | 23.55626 | 8.51546 | 0 |
| Mean | **2042.739274** | **336.293** | **133.428346** | **45.650289** | **12.03842** | **0.134** |
| Std | **227.4791267** | **38.93999** | **56.8085391** | **37.9146789** | **7.274574663** | **0.284612797** |
| t-test |  | **8.88E-10** | **5.22604E-10** | **4.16124E-10** | **4.02764E-10** | **5.8885E-07** |

**Table 2: This table presents the mean, standard deviation and statistical analysis of TMRE intensity values measured at various time points during anaesthesia (n=10)**

| S.No. | Control | Treatments (TMRE) | | | | |
| --- | --- | --- | --- | --- | --- | --- |
|  |  | **1 hour** | **2 hour** | **4 hour** | **6 hour** | **8 hour** |
| 1 | 1337.628 | 115.8426 | 13.27991206 | 9.638002287 | 1.105206656 | 0 |
| 2 | 1367.334 | 89.40294 | 13.64486803 | 9.226435088 | 0.917714612 | 0 |
| 3 | 1112.081 | 97.0556 | 12.42683473 | 7.774817228 | 0.826579072 | 0.3 |
| 4 | 1346.986 | 110.1883 | 14.83344717 | 8.070947468 | 1.087967301 | 0 |
| 5 | 1214.024 | 101.1858 | 13.12627387 | 9.210841909 | 0.977714574 | 0 |
| 6 | 1384.448 | 122.441 | 13.42639072 | 11.14145861 | 1.073012891 | 0 |
| 7 | 1181.359 | 93.7056 | 14.51899118 | 9.830543993 | 0.813392586 | 0 |
| 8 | 1362.873 | 92.99864 | 14.2926972 | 10.94933266 | 1.073185448 | 0 |
| 9 | 1303.817 | 92.78179 | 17.1783564 | 9.943688832 | 0.963764299 | 0 |
| 10 | 974.6664 | 94.56141 | 12.68859281 | 10.59708664 | 0.991263081 | 0 |
| Mean | **1258.522** | **101.0164** | **13.94163642** | **9.638315472** | **0.982980052** | **0.03** |
| Std | **135.2672** | **11.25832** | **1.376358** | **1.120976736** | **0.105488769** | **0.09486833** |
| t-test |  | **4.82E-10** | **3.14567E-10** | **3.13749E-10** | **2.96458E-10** | **2.96214E-10** |

**Table 3: This table presents the mean, standard deviation and statistical analysis of MTG and LTR PCC (Pearson’s Correlation Coefficient) values measured at various time points during anaesthesia (n=10)**

| S.No. | Control | Treatments (PCC) | | | | |
| --- | --- | --- | --- | --- | --- | --- |
|  |  | **1 hour** | **2 hour** | **4 hour** | **6 hour** | **8 hour** |
| 1 | 0.17 | 0.65 | 0.3 | 0.23 | 0.2 | 0.07 |
| 2 | 0.16 | 0.72 | 0.22 | 0.28 | 0.2 | 0.06 |
| 3 | 0.18 | 0.63 | 0.31 | 0.25 | 0.34 | 0.02 |
| 4 | 0.14 | 0.59 | 0.37 | 0.3 | 0.25 | 0.16 |
| 5 | 0.24 | 0.7 | 0.3 | 0.23 | 0.27 | 0.03 |
| 6 | 0.2 | 0.75 | 0.35 | 0.27 | 0.21 | 0.02 |
| 7 | 0.07 | 0.67 | 0.28 | 0.17 | 0.21 | 0.09 |
| 8 | 0.17 | 0.6 | 0.32 | 0.24 | 0.15 | 0.09 |
| 9 | 0.21 | 0.67 | 0.29 | 0.35 | 0.23 | 0.07 |
| 10 | 0.19 | 0.66 | 0.3 | 0.29 | 0.28 | 0.09 |
| Mean | **0.173** | **0.664** | **0.304** | **0.261** | **0.234** | **0.07** |
| Std | **0.045717** | **0.050376361** | **0.040331956** | **0.048865** | **0.053166** | **0.042164** |
| t-test |  | **6.28121E-10** | **5.14764E-05** | **0.000252** | **0.01065** | **0.002136** |

**Table 4: This table presents the mean, standard deviation and statistical analysis of MTG and LTR M 1 (Mander’s Coefficient 1) values measured at various time points during anaesthesia (n=10)**

| S.No. | Control | Treatments (M1) | | | | |
| --- | --- | --- | --- | --- | --- | --- |
|  |  | **1 hour** | **2 hour** | **4 hour** | **6 hour** | **8 hour** |
| 1 | 0.14 | 0.95 | 0.87 | 1 | 1 | 0.96 |
| 2 | 0.14 | 0.99 | 0.91 | 0.98 | 1 | 0.91 |
| 3 | 0.19 | 0.91 | 0.94 | 1 | 0.91 | 0.91 |
| 4 | 0.14 | 0.93 | 0.97 | 0.98 | 1 | 1 |
| 5 | 0.13 | 0.84 | 1 | 0.98 | 1 | 0.99 |
| 6 | 0.18 | 0.94 | 0.92 | 0.98 | 1 | 0.94 |
| 7 | 0.07 | 0.96 | 1 | 0.92 | 1 | 0.99 |
| 8 | 0.18 | 1 | 0.98 | 0.99 | 1 | 0.93 |
| 9 | 0.12 | 0.95 | 1 | 1 | 0.99 | 0.99 |
| 10 | 0.23 | 0.88 | 1 | 1 | 0.97 | 1 |
| Mean | **0.152** | **0.935** | **0.959** | **0.983** | **0.987** | **0.962** |
| Std | **0.044422** | **0.04836206** | **0.04653553** | **0.02406011** | **0.02869379** | **0.036757463** |
| t-test |  | **8.18382E-11** | **3.28571E-11** | **3.73539E-14** | **1.3017E-11** | **2.21506E-11** |

**Table 5: This table presents the mean, standard deviation and statistical analysis of MTG and LTR M 2 (Mander’s Coefficient 2) values measured at various time points during anaesthesia (n=10)**

| S.No. | Control | Treatments (M2) | | | | |
| --- | --- | --- | --- | --- | --- | --- |
|  |  | **1 hour** | **2 hour** | **4 hour** | **6 hour** | **8 hour** |
| 1 | 0.83 | 0.57 | 0.23 | 0.24 | 0.13 | 0.04 |
| 2 | 0.8 | 0.73 | 0.44 | 0.23 | 0.21 | 0.07 |
| 3 | 0.88 | 0.61 | 0.36 | 0.28 | 0.05 | 0.01 |
| 4 | 0.81 | 0.53 | 0.34 | 0.2 | 0.08 | 0.04 |
| 5 | 0.74 | 0.61 | 0.4 | 0.22 | 0.06 | 0.05 |
| 6 | 0.79 | 0.52 | 0.31 | 0.15 | 0.05 | 0 |
| 7 | 0.83 | 0.63 | 0.31 | 0.23 | 0.17 | 0.01 |
| 8 | 0.84 | 0.61 | 0.24 | 0.25 | 0.13 | 0.09 |
| 9 | 0.85 | 0.54 | 0.31 | 0.19 | 0.13 | 0.03 |
| 10 | 0.82 | 0.61 | 0.31 | 0.27 | 0.12 | 0 |
| Mean | **0.819** | **0.596** | **0.325** | **0.226** | **0.113** | **0.034** |
| Std | **0.037845** | **0.061318839** | **0.064506675** | **0.03864367** | **0.053135048** | **0.030258149** |
| t-test |  | **5.39253E-06** | **2.42406E-08** | **5.1918E-12** | **4.28981E-11** | **3.97349E-12** |

**Table 6: This table presents the mean, standard deviation and statistical analysis of ROS generation using H2DCFDA probe intensity values measured at various time points during anaesthesia (n=10)**

| S.No. | Control | Treatments (H2DCFDA) | | | | |
| --- | --- | --- | --- | --- | --- | --- |
|  |  | **1 hour** | **2 hour** | **4 hour** | **6 hour** | **8 hour** |
| 1 | 44.01735 | 23.95755 | 485.1159882 | 975.742545 | 2043.276 | 3080.264532 |
| 2 | 7.420655 | 63.41274 | 304.6271604 | 1139.43457 | 2036.653 | 3407.333221 |
| 3 | 61.28412 | 171.0811 | 397.222253 | 1039.1326 | 2006.608 | 3199.298351 |
| 4 | 5.107463 | 240.4499 | 417.7853546 | 970.687807 | 2133.096 | 3082.733912 |
| 5 | 39.5844 | 319.8 | 387.0611157 | 867.23922 | 2086.029 | 3008.888704 |
| 6 | 9.21001 | 191.8026 | 256.813099 | 1093.74928 | 2143.661 | 3247.734941 |
| 7 | 7.303947 | 221.1639 | 210.9508475 | 932.76057 | 2005.109 | 3176.817703 |
| 8 | 0.059118 | 61.63844 | 281.5969468 | 1005.30584 | 2059.238 | 3191.368303 |
| 9 | 37.13059 | 61.88808 | 339.0653991 | 1075.77417 | 2101.317 | 3165.032912 |
| 10 | 7.787546 | 234.9127 | 236.3911247 | 1034.65002 | 1971.562 | 3032.586444 |
| Mean | **21.89052** | **159.0107** | **331.6629289** | **1013.44766** | **2058.655** | **3159.205902** |
| Std | **21.41365** | **99.79042** | **88.74110279** | **80.6353852** | **56.97874** | **116.9510647** |
| t-test |  | **0.002353** | **4.42215E-07** | **4.6924E-11** | **4.24E-15** | **3.71029E-14** |

**Table 7: This table presents the mean, standard deviation and statistical analysis of Vesicles in plants generation using FM4-64 probe intensity values measured at various time points during anaesthesia (n=10)**

| S.No. | Control | Treatments (FM4-64) | | | | |
| --- | --- | --- | --- | --- | --- | --- |
|  |  | **1 hour** | **2 hour** | **4 hour** | **6 hour** | **8 hour** |
| 1 | 776.318 | 37.578 | 45.252 | 11.715 | 3.948 | 5.53 |
| 2 | 594.785 | 103.975 | 50.084 | 28.185 | 9.06 | 3.53 |
| 3 | 496.74 | 27.048 | 73.901 | 13.451 | 21.687 | 1.66 |
| 4 | 483.022 | 38.485 | 124.664 | 12.461 | 3.948 | 6.66 |
| 5 | 493.452 | 105.235 | 43.327 | 24.251 | 0.333 | 3.034 |
| 6 | 463.536 | 73.779 | 50.084 | 36.1 | 31.653 | 3.034 |
| 7 | 179.74 | 157.743 | 16.32 | 20.408 | 15.793 | 6.66 |
| 8 | 1016.452 | 134.905 | 8.37 | 81.598 | 21.687 | 4.412 |
| 9 | 1016.452 | 153.959 | 46.715 | 11.715 | 0.333 | 2.412 |
| 10 | 681.122 | 60.502 | 57.321 | 12.461 | 15.793 | 3.66 |
| Mean | **620.1619** | **89.3209** | **51.6038** | **25.2345** | **12.4235** | **4.0592** |
| Std | **260.6478** | **49.00928** | **31.83136654** | **21.49573675** | **10.60108647** | **1.72649142** |
| t-test |  | **0.000109** | **9.04768E-05** | **4.05785E-05** | **4.53224E-05** | **3.85182E-05** |

**Table 8: This table presents the mean, standard deviation and statistical analysis of Vesicles area in plants generation using FM4-64 probe values measured at various time points during anaesthesia (n=10)**

| S.No. | Control | Treatments (FM4-64) | | | | |
| --- | --- | --- | --- | --- | --- | --- |
|  |  | **1 hour** | **2 hour** | **4 hour** | **6 hour** | **8 hour** |
| 1 | 83.183 | 14.176 | 18.344 | 8.779 | 3.888 | 2.16 |
| 2 | 64.963 | 25.845 | 19.973 | 16.748 | 4.754 | 1.16 |
| 3 | 75.949 | 14.627 | 21.055 | 9.145 | 16.875 | 0.25 |
| 4 | 77.964 | 16.384 | 37.328 | 6.541 | 3.888 | 0.25 |
| 5 | 79.695 | 34.448 | 18.194 | 13.164 | 0.429 | 1.577 |
| 6 | 64.965 | 22.157 | 19.973 | 23.631 | 22.333 | 3.577 |
| 7 | 34.759 | 49.279 | 8.036 | 8.95 | 11.964 | 1.25 |
| 8 | 89.748 | 40.371 | 3.3 | 31.563 | 16.875 | 1.749 |
| 9 | 89.748 | 42.17 | 15.014 | 8.779 | 0.429 | 0.749 |
| 10 | 78.724 | 18.801 | 15.106 | 6.541 | 11.964 | 1.25 |
| Mean | **73.9698** | **27.8258** | **17.6323** | **13.3841** | **9.3399** | **1.3972** |
| Std | **16.1855** | **12.80702** | **8.957658015** | **8.303525161** | **7.713142441** | **0.979874232** |
| t-test |  | **0.000153** | **3.19619E-06** | **1.46818E-06** | **2.60827E-06** | **1.97761E-07** |

**Table 9: This table presents the mean, standard deviation and statistical analysis of NRF2 and DAPI PCC (Pearson’s Correlation Coefficient) values measured at various time points during anaesthesia (n=10)**

| S.No. | Control | Treatments (NRF2-DAPI) | | | | |
| --- | --- | --- | --- | --- | --- | --- |
|  |  | **1 hour** | **2 hour** | **4 hour** | **6 hour** | **8 hour** |
| 1 | 0.63 | 0.7 | 0.72 | 0.75 | 0.64 | 0.67 |
| 2 | 0.47 | 0.7 | 0.52 | 0.57 | 0.68 | 0.76 |
| 3 | 0.51 | 0.77 | 0.59 | 0.65 | 0.63 | 0.62 |
| 4 | 0.5 | 0.65 | 0.71 | 0.69 | 0.74 | 0.66 |
| 5 | 0.35 | 0.54 | 0.76 | 0.62 | 0.74 | 0.77 |
| 6 | 0.22 | 0.59 | 0.54 | 0.75 | 0.62 | 0.68 |
| 7 | 0.15 | 0.45 | 0.61 | 0.61 | 0.78 | 0.69 |
| 8 | 0.3 | 0.48 | 0.63 | 0.58 | 0.6 | 0.75 |
| 9 | 0.18 | 0.47 | 0.73 | 0.58 | 0.8 | 0.67 |
| 10 | 0.07 | 0.59 | 0.6 | 0.71 | 0.65 | 0.79 |
| Mean | **0.338** | **0.594** | **0.641** | **0.651** | **0.688** | **0.706** |
| Std | **0.184439** | **0.109869** | **0.083858** | **0.069833** | **0.071461** | **0.056804** |
| t-test |  | **0.00012** | **0.000643** | **0.000493** | **0.00058** | **0.000404** |

**Table 10: This table presents the mean, standard deviation and statistical analysis of NRF2 and DAPI M 1 (Mander’s Coefficient 1) values measured at various time points during anaesthesia (n=10)**

| S.No. | Control | Treatments (NRF2+DAPI) | | | | |
| --- | --- | --- | --- | --- | --- | --- |
|  |  | **1 hour** | **2 hour** | **4 hour** | **6 hour** | **8 hour** |
| 1 | 0.872 | 0.892 | 0.849 | 0.865 | 0.755 | 0.898 |
| 2 | 0.841 | 0.857 | 0.747 | 0.85 | 0.774 | 0.56 |
| 3 | 0.681 | 0.968 | 0.857 | 0.719 | 0.792 | 0.993 |
| 4 | 0.821 | 0.822 | 0.921 | 0.748 | 0.761 | 0.99 |
| 5 | 0.782 | 0.907 | 0.874 | 0.821 | 0.994 | 0.97 |
| 6 | 0.872 | 0.702 | 0.877 | 0.874 | 0.981 | 0.87 |
| 7 | 0.753 | 0.841 | 0.8 | 0.966 | 0.993 | 0.719 |
| 8 | 0.892 | 0.772 | 0.776 | 0.877 | 0.996 | 0.936 |
| 9 | 0.923 | 0.729 | 0.873 | 0.766 | 0.699 | 0.863 |
| 10 | 0.837 | 0.774 | 0.723 | 0.733 | 0.911 | 0.966 |
| Mean | **0.8274** | **0.8264** | **0.8297** | **0.8219** | **0.8656** | **0.8765** |
| Std | **0.071986** | **0.083458** | **0.064565** | **0.079051** | **0.120109** | **0.138409** |
| t-test |  | **0.983168** | **0.94359** | **0.865168** | **0.439085** | **0.36664** |

**Table 11: This table presents the mean, standard deviation and statistical analysis of NRF2 and DAPI M 2 (Mander’s Coefficient 2) values measured at various time points during anaesthesia (n=10)**

| S.No. | Control | Treatments (NRF2+DAPI) | | | | |
| --- | --- | --- | --- | --- | --- | --- |
|  |  | **1 hour** | **2 hour** | **4 hour** | **6 hour** | **8 hour** |
| 1 | 0.874 | 0.942 | 0.855 | 0.996 | 0.942 | 0.978 |
| 2 | 0.921 | 0.94 | 0.993 | 0.992 | 0.991 | 0.97 |
| 3 | 0.955 | 0.944 | 0.918 | 0.991 | 0.955 | 0.944 |
| 4 | 0.817 | 0.958 | 0.991 | 0.968 | 0.929 | 0.993 |
| 5 | 0.84 | 0.8 | 1 | 0.976 | 0.997 | 0.968 |
| 6 | 0.867 | 0.863 | 0.971 | 0.899 | 0.964 | 0.98 |
| 7 | 0.998 | 0.951 | 0.979 | 0.976 | 0.925 | 0.998 |
| 8 | 0.855 | 0.983 | 0.97 | 0.951 | 0.935 | 0.95 |
| 9 | 0.875 | 0.911 | 0.967 | 0.974 | 0.999 | 0.991 |
| 10 | 0.245 | 0.803 | 0.958 | 0.938 | 0.997 | 0.998 |
| Mean | **0.8247** | **0.9095** | **0.9602** | **0.9661** | **0.9634** | **0.977** |
| Std | **0.210897** | **0.065107** | **0.043555** | **0.029752** | **0.030332** | **0.019125** |
| t-test |  | **0.166724** | **0.078739** | **0.053095** | **0.08326** | **0.055268** |

**Table 12: This table presents the mean, standard deviation and statistical analysis of DAPI intensity values measured at various time points during anaesthesia (n=10)**

| S.No. | Control | Treatments (DAPI) | | | | |
| --- | --- | --- | --- | --- | --- | --- |
|  |  | **1 hour** | **2 hour** | **4 hour** | **6 hour** | **8 hour** |
| 1 | 2382.57 | 717.03 | 500.81 | 691.62 | 296.62 | 310.7 |
| 2 | 1975.31 | 864.39 | 527.28 | 731.75 | 294.29 | 310.6 |
| 3 | 2170.07 | 769.66 | 551.78 | 717.42 | 351.17 | 393.84 |
| 4 | 2049.98 | 812.3 | 541.98 | 670.04 | 326.89 | 313.72 |
| 5 | 2290.02 | 759.16 | 698.37 | 792.2 | 311.92 | 340.95 |
| 6 | 2240.9 | 780.41 | 625.84 | 724 | 336.43 | 338.56 |
| 7 | 1956.87 | 798.27 | 608.29 | 766.11 | 276.24 | 325.88 |
| 8 | 2089.95 | 791.54 | 486.15 | 712.6 | 314.98 | 299.83 |
| 9 | 2459.62 | 690.14 | 481.78 | 717.49 | 327.65 | 284.04 |
| 10 | 2333.04 | 756.76 | 533.54 | 699.56 | 308.09 | 356.6 |
| Mean | **2194.833** | **773.966** | **555.582** | **722.279** | **314.428** | **327.472** |
| Std | **-174.182** | **-48.7503418** | **-69.0061** | **-35.3208** | **-22.0906** | **-31.5626** |
| t-test |  | **2.21733E-15** | **3.34E-16** | **8.71E-16** | **9.37E-18** | **1.23E-17** |

**Table 13: This table presents the mean, standard deviation and statistical analysis of DAPI area (%) values measured at various time points during anaesthesia (n=10)**

| S.No. | Control | Treatments (DAPI) | | | | |
| --- | --- | --- | --- | --- | --- | --- |
|  |  | **1 hour** | **2 hour** | **4 hour** | **6 hour** | **8 hour** |
| 1 | 75.82 | 48.89 | 39.97 | 48.38 | 50.93 | 37.4 |
| 2 | 74.1 | 36.34 | 31 | 52.19 | 47.28 | 32.07 |
| 3 | 73.46 | 25.96 | 52.6 | 34.32 | 49.34 | 31.65 |
| 4 | 71.06 | 66.5 | 37.67 | 51.11 | 40.65 | 40.34 |
| 5 | 74.85 | 31.74 | 32.3 | 55.14 | 39.61 | 30.96 |
| 6 | 79.23 | 16.92 | 38.35 | 47.04 | 44.37 | 26.61 |
| 7 | 76.24 | 47.06 | 42.21 | 41.59 | 38.86 | 32.82 |
| 8 | 74.86 | 45.19 | 41.82 | 41.48 | 47.74 | 39.58 |
| 9 | 79.52 | 45.25 | 48.89 | 51.97 | 44.95 | 36.89 |
| 10 | 78.58 | 49.62 | 43.99 | 36.15 | 28.08 | 26.81 |
| Mean | **75.772** | **41.347** | **40.88** | **45.937** | **43.181** | **33.513** |
| Std | **-2.71083** | **-14.0295554** | **-6.68835638** | **-7.17742147** | **-6.71630595** | **-4.8883309** |
| t-test |  | **4.87972E-07** | **9.3626E-12** | **3.40083E-10** | **3.10405E-11** | **4.3349E-15** |

**Table 14: This table presents the mean, standard deviation and statistical analysis of NRF2 and DAPI PCC (Pearson’s corelation coefficient) values measured at various time points during anaesthesia (n=10)**

| S.No. | Control | Treatments (DAPI +PI) | | | | |
| --- | --- | --- | --- | --- | --- | --- |
|  |  | **1 hour** | **2 hour** | **4 hour** | **6 hour** | **8 hour** |
| 1 | 0.22 | 0.22 | 0.3 | 0.47 | 0.55 | 0.86 |
| 2 | 0.16 | 0.2 | 0.27 | 0.42 | 0.56 | 0.86 |
| 3 | 0.08 | 0.23 | 0.21 | 0.44 | 0.52 | 0.83 |
| 4 | 0.16 | 0.28 | 0.42 | 0.43 | 0.54 | 0.85 |
| 5 | 0.2 | 0.14 | 0.35 | 0.42 | 0.58 | 0.89 |
| 6 | 0.07 | 0.18 | 0.39 | 0.43 | 0.48 | 0.93 |
| 7 | 0.22 | 0.24 | 0.26 | 0.38 | 0.5 | 0.9 |
| 8 | 0.21 | 0.23 | 0.38 | 0.47 | 0.51 | 0.86 |
| 9 | 0.12 | 0.14 | 0.25 | 0.45 | 0.52 | 0.9 |
| 10 | 0.21 | 0.12 | 0.3 | 0.5 | 0.57 | 0.9 |
| Mean | **0.165** | **0.198** | **0.313** | **0.441** | **0.533** | **0.878** |
| Std | **0.054452** | **0.049153** | **0.065429351** | **0.03176476** | **0.030675723** | **0.028913665** |
| t-test |  | **0.193876** | **5.83155E-05** | **1.16313E-10** | **8.12081E-13** | **6.10797E-18** |

**Table 15: This table presents the mean, standard deviation and statistical analysis of DAPI and PI M 1 (Mander’s Coefficient 1) values measured at various time points during anaesthesia (n=10)**

| S.No. | Control | Treatments (DAPI+PI) | | | | |
| --- | --- | --- | --- | --- | --- | --- |
|  |  | **1 hour** | **2 hour** | **4 hour** | **6 hour** | **8 hour** |
| 1 | 0.04 | 0.37 | 0.44 | 0.59 | 0.96 | 0.99 |
| 2 | 0.03 | 0.35 | 0.42 | 0.56 | 0.75 | 0.92 |
| 3 | 0.02 | 0.4 | 0.41 | 0.52 | 0.84 | 0.93 |
| 4 | 0.06 | 0.32 | 0.45 | 0.6 | 0.71 | 0.86 |
| 5 | 0.03 | 0.36 | 0.45 | 0.47 | 0.82 | 0.97 |
| 6 | 0 | 0.32 | 0.48 | 0.55 | 0.89 | 0.86 |
| 7 | 0.09 | 0.31 | 0.39 | 0.59 | 0.82 | 0.98 |
| 8 | 0.08 | 0.34 | 0.52 | 0.47 | 0.86 | 0.85 |
| 9 | 0.09 | 0.43 | 0.39 | 0.59 | 0.83 | 0.96 |
| 10 | 0.05 | 0.38 | 0.46 | 0.54 | 0.75 | 0.87 |
| Mean | **0.049** | **0.358** | **0.441** | **0.548** | **0.823** | **0.919** |
| Std | **0.029138** | **0.03627671** | **0.03858756** | **0.046** | **0.069577295** | **0.052239832** |
| t-test |  | **1.0297E-13** | **3.2113E-15** | **3.7373E-16** | **5.08558E-17** | **1.03281E-19** |

**Table 16: This table presents the mean, standard deviation and statistical analysis of DAPI and PI M 2 (Mander’s Coefficient 2) values measured at various time points during anaesthesia (n=10)**

| S.No. | Control | Treatments (DAPI+PI) | | | | |
| --- | --- | --- | --- | --- | --- | --- |
|  |  | **1 hour** | **2 hour** | **4 hour** | **6 hour** | **8 hour** |
| 1 | 1 | 1 | 0.99 | 0.99 | 0.99 | 1 |
| 2 | 0.99 | 0.98 | 0.98 | 0.99 | 0.99 | 0.97 |
| 3 | 0.99 | 0.99 | 0.98 | 0.99 | 1 | 0.98 |
| 4 | 1 | 1 | 0.98 | 1 | 0.98 | 1 |
| 5 | 1 | 0.98 | 1 | 0.99 | 1 | 1 |
| 6 | 1 | 1 | 0.98 | 0.99 | 0.99 | 0.98 |
| 7 | 1 | 0.99 | 1 | 0.98 | 0.99 | 0.98 |
| 8 | 1 | 0.99 | 0.98 | 0.99 | 0.99 | 0.98 |
| 9 | 0.99 | 0.98 | 0.98 | 0.97 | 0.98 | 0.98 |
| 10 | 0.99 | 0.97 | 1 | 0.99 | 1 | 1 |
| Mean | **0.996** | **0.988** | **0.987** | **0.988** | **0.991** | **0.987** |
| Std | **0.004899** | **0.009798** | **0.009** | **0.007483** | **0.007** | **0.011** |
| t-test |  | **0.041861** | **0.016816** | **0.015181** | **0.096157** | **0.03778** |

**Table 17: This table presents the mean, standard deviation and statistical analysis of PI positive cell values measured at various time points during anaesthesia (n=10)**

| S.No. | Control | Treatments (PI) | | | | |
| --- | --- | --- | --- | --- | --- | --- |
|  |  | **1 hour** | **2 hour** | **4 hour** | **6 hour** | **8 hour** |
| 1 | 2 | 19 | 27 | 42 | 63 | 98 |
| 2 | 3 | 21 | 25 | 42 | 46 | 83 |
| 3 | 3 | 24 | 30 | 50 | 58 | 82 |
| 4 | 3 | 21 | 29 | 35 | 66 | 81 |
| 5 | 3 | 17 | 27 | 32 | 63 | 84 |
| 6 | 3 | 21 | 30 | 35 | 58 | 79 |
| 7 | 3 | 21 | 34 | 45 | 60 | 59 |
| 8 | 3 | 19 | 25 | 42 | 54 | 69 |
| 9 | 3 | 23 | 22 | 38 | 58 | 80 |
| 10 | 3 | 20 | 30 | 41 | 58 | 78 |
| Mean | **2.9** | **20.6** | **27.9** | **40.2** | **58.4** | **79.3** |
| Std | **0.3** | **1.907878403** | **3.238826948** | **5.055689864** | **5.257375771** | **9.5713113** |
| t-test |  | **3.73266E-16** | **8.15367E-15** | **1.71439E-14** | **3.16633E-17** | **4.24918E-15** |

**Table 18: This table presents the mean, standard deviation and statistical analysis of PI intensity values measured at various time points during anaesthesia (n=10)**

| S.No. | Control | Treatments (PI) | | | | |
| --- | --- | --- | --- | --- | --- | --- |
|  |  | **1 hour** | **2 hour** | **4 hour** | **6 hour** | **8 hour** |
| 1 | 4.5 | 68.34 | 144.68 | 244.98 | 424.94 | 617.03 |
| 2 | 4.86 | 73.72 | 147.77 | 238.22 | 382.22 | 513.82 |
| 3 | 4.37 | 66.11 | 140.01 | 240.74 | 311.04 | 501.28 |
| 4 | 4.67 | 70.03 | 129.38 | 240.4 | 417.93 | 449.89 |
| 5 | 4.14 | 68.97 | 120.17 | 225.7 | 383.48 | 437.78 |
| 6 | 4.6 | 69.61 | 144.18 | 211.94 | 500.06 | 495.56 |
| 7 | 4.02 | 81.17 | 116.75 | 241.26 | 401.65 | 592.38 |
| 8 | 4.04 | 66.47 | 151.06 | 321 | 408.9 | 563.64 |
| 9 | 4.83 | 88.88 | 117.38 | 213.3 | 424.32 | 482.47 |
| 10 | 4.25 | 72.97 | 140.85 | 248.79 | 470.99 | 500.29 |
| Mean | **4.428** | **72.627** | **135.223** | **242.633** | **412.553** | **515.414** |
| Std | **0.296068** | **6.829595962** | **12.46128729** | **28.81656609** | **48.57531205** | **55.50341507** |
| t-test |  | **8.35816E-17** | **3.42292E-17** | **2.28697E-15** | **1.71848E-15** | **3.44699E-16** |

**Table 19: This table presents the mean, standard deviation and statistical analysis of Euchromatin (H3K4me3) domain no. values measured at various time points during anaesthesia (n=10)**

| S.No. | Control | Treatments (H3K4me3) | | | | |
| --- | --- | --- | --- | --- | --- | --- |
|  |  | **1 hour** | **2 hour** | **4 hour** | **6 hour** | **8 hour** |
| 1 | 52 | 18 | 22 | 28 | 14 | 41 |
| 2 | 165 | 17 | 39 | 22 | 21 | 48 |
| 3 | 64 | 22 | 39 | 39 | 15 | 46 |
| 4 | 66 | 49 | 32 | 35 | 21 | 66 |
| 5 | 100 | 52 | 29 | 18 | 29 | 17 |
| 6 | 48 | 49 | 99 | 30 | 12 | 56 |
| 7 | 174 | 46 | 87 | 35 | 22 | 31 |
| 8 | 134 | 31 | 24 | 13 | 29 | 35 |
| 9 | 64 | 12 | 66 | 29 | 6 | 21 |
| 10 | 76 | 9 | 57 | 14 | 34 | 28 |
| Mean | **94.3** | **30.5** | **49.4** | **26.3** | **20.3** | **38.9** |
| Std | **44.56915974** | **16.1322658** | **25.57029527** | **8.672369918** | **8.271033793** | **14.7** |
| t-test |  | **0.000771576** | **0.017299644** | **0.000281377** | **0.000115982** | **0.002332171** |

**Table 20: This table presents the mean, standard deviation and statistical analysis of Euchromatin (H3K4me3) %age area values measured at various time points during anaesthesia (n=10)**

| S.No. | Control | Treatments (H3K4me3) | | | | |
| --- | --- | --- | --- | --- | --- | --- |
|  |  | **1 hour** | **2 hour** | **4 hour** | **6 hour** | **8 hour** |
| 1 | 28.16436927 | 25.41172596 | 29.08964651 | 21.51815524 | 27.69190659 | 9.783414531 |
| 2 | 42.09219136 | 25.14473764 | 19.82584755 | 22.59060403 | 22.18301576 | 4.340720335 |
| 3 | 35.57130281 | 24.66857415 | 36.75368716 | 22.24422653 | 21.51268785 | 7.058005235 |
| 4 | 29.93084801 | 23.06293398 | 23.21733554 | 20.09708034 | 15.36392225 | 8.983329142 |
| 5 | 32.36813375 | 22.72151312 | 23.9840904 | 28.33681571 | 13.25966851 | 6.749725174 |
| 6 | 32.00048139 | 25.81026807 | 20.69263974 | 19.31114309 | 26.26532537 | 5.568348958 |
| 7 | 29.14054541 | 25.28629051 | 31.44071339 | 19.94969677 | 16.75635089 | 6.748571103 |
| 8 | 28.42805455 | 17.45484825 | 25.46797866 | 23.87660485 | 22.52502781 | 7.154725833 |
| 9 | 31.19360809 | 18.21674336 | 32.67243787 | 21.30921244 | 18.02750951 | 7.621736062 |
| 10 | 27.88816936 | 23.34899504 | 22.13163532 | 24.77809159 | 21.30128141 | 7.469875446 |
| Mean | **31.6777704** | **23.11266301** | **26.52760121** | **22.40116306** | **20.4886696** | **7.147845182** |
| Std | **4.137509887** | **2.828954363** | **5.383952992** | **2.568228447** | **4.387389817** | **1.460968834** |
| t-test |  | **7.06511E-05** | **0.035339126** | **2.03164E-05** | **2.77312E-05** | **1.96434E-12** |

**Table 21: This table presents the mean, standard deviation and statistical analysis of Euchromatin (H3K4me3) mean domain area values measured at various time points during anaesthesia (n=10)**

| S.No. | Control | Treatments (H3K4me3) | | | | |
| --- | --- | --- | --- | --- | --- | --- |
|  |  | **1 hour** | **2 hour** | **4 hour** | **6 hour** | **8 hour** |
| 1 | 281816.2393 | 285740.7407 | 444292.9293 | 193611.1111 | 249714.2857 | 144195.122 |
| 2 | 143865.3199 | 269673.2026 | 176438.7464 | 623333.333 | 231085.7143 | 140766.6667 |
| 3 | 358038.1944 | 293737.3737 | 177350.4274 | 285702.5641 | 265706.6667 | 134121.7391 |
| 4 | 518080.8081 | 94897.95918 | 889166.6667 | 264190.4762 | 308952.381 | 147224.2424 |
| 5 | 318711.1111 | 157094.0171 | 144137.931 | 118580.2469 | 252873.5632 | 260047.0588 |
| 6 | 738611.1111 | 187981.8594 | 211975.3086 | 228629.6296 | 168400 | 107085.7143 |
| 7 | 185957.8544 | 131739.1304 | 589934.6405 | 387977.1429 | 189527.2727 | 183432.2581 |
| 8 | 274510.7794 | 217885.3047 | 355879.6296 | 329600 | 268137.931 | 167314.2857 |
| 9 | 184895.8333 | 399537.037 | 487500 | 121422.2222 | 385600 | 230400 |
| 10 | 404839.1813 | 341728.3951 | 144990.2534 | 126520 | 504564.7059 | 252571.4286 |
| Mean | **340932.6432** | **238001.502** | **362166.6533** | **267956.6726** | **282456.2521** | **176715.8516** |
| Std | **170149.0207** | **91904.6131** | **231950.7308** | **146864.0043** | **93482.15531** | **50592.44727** |
| t-test |  | **0.127718204** | **0.827239266** | **0.342952452** | **0.378126138** | **0.01247894** |

**Table 22: This table presents the mean, standard deviation and statistical analysis of Euchromatin (H3K4me3) mean domain perimeter values measured at various time points during anaesthesia (n=10)**

| S.No. | Control | Treatments (H3K4me3) | | | | |
| --- | --- | --- | --- | --- | --- | --- |
|  |  | **1 hour** | **2 hour** | **4 hour** | **6 hour** | **8 hour** |
| 1 | 281816.2393 | 285740.7407 | 444292.9293 | 193611.1111 | 249714.2857 | 144195.122 |
| 2 | 143865.3199 | 269673.2026 | 176438.7464 | 623333.333 | 231085.7143 | 140766.6667 |
| 3 | 358038.1944 | 293737.3737 | 177350.4274 | 285702.5641 | 265706.6667 | 134121.7391 |
| 4 | 518080.8081 | 94897.95918 | 889166.6667 | 264190.4762 | 308952.381 | 147224.2424 |
| 5 | 318711.1111 | 157094.0171 | 144137.931 | 118580.2469 | 252873.5632 | 260047.0588 |
| 6 | 738611.1111 | 187981.8594 | 211975.3086 | 228629.6296 | 168400 | 107085.7143 |
| 7 | 185957.8544 | 131739.1304 | 589934.6405 | 387977.1429 | 189527.2727 | 183432.2581 |
| 8 | 274510.7794 | 217885.3047 | 355879.6296 | 329600 | 268137.931 | 167314.2857 |
| 9 | 184895.8333 | 399537.037 | 487500 | 121422.2222 | 385600 | 230400 |
| 10 | 404839.1813 | 341728.3951 | 144990.2534 | 126520 | 504564.7059 | 252571.4286 |
| Mean | **340932.6432** | **238001.502** | **362166.6533** | **267956.6726** | **282456.2521** | **176715.8516** |
| Std | **170149.0207** | **91904.6131** | **231950.7308** | **146864.0043** | **93482.15531** | **50592.44727** |
| t-test |  | **0.127718204** | **0.827239266** | **0.342952452** | **0.378126138** | **0.01247894** |

**Table 23: This table presents the mean, standard deviation and statistical analysis of Euchromatin (H3K4me3) mean circularity values measured at various time points during anaesthesia (n=10)**

| S.No. | Control | Treatments (H3K4me3) | | | | |
| --- | --- | --- | --- | --- | --- | --- |
|  |  | **1 hour** | **2 hour** | **4 hour** | **6 hour** | **8 hour** |
| 1 | 0.589298045 | 0.460529701 | 0.386973928 | 0.803955798 | 0.594197323 | 0.740819801 |
| 2 | 0.721408682 | 0.583013909 | 0.339215339 | 0.439841122 | 0.412204097 | 0.775583273 |
| 3 | 0.640484691 | 0.481533917 | 0.410324838 | 0.65036408 | 0.524913585 | 0.800645693 |
| 4 | 0.573897757 | 0.576365588 | 0.576790532 | 0.63443035 | 0.391520779 | 0.773045302 |
| 5 | 0.628726315 | 0.44279687 | 0.603927978 | 0.611029059 | 0.636561354 | 0.639718273 |
| 6 | 0.630535929 | 0.346975632 | 0.40752129 | 0.627820877 | 0.569595718 | 0.828245366 |
| 7 | 0.713667643 | 0.323117971 | 0.609193205 | 0.727176044 | 0.489965154 | 0.701280032 |
| 8 | 0.73660512 | 0.28368242 | 0.499889812 | 0.598772883 | 0.665584115 | 0.637745053 |
| 9 | 0.593081641 | 0.194638981 | 0.567138441 | 0.634802439 | 0.500528364 | 0.681250073 |
| 10 | 0.654709644 | 0.268967892 | 0.44494686 | 0.496034546 | 0.426234142 | 0.661198252 |
| Mean | **0.648241547** | **0.396162288** | **0.484592222** | **0.62242272** | **0.521130463** | **0.723953112** |
| Std | **0.055029459** | **0.125773446** | **0.094342963** | **0.097485362** | **0.089976533** | **0.06551485** |
| t-test |  | **3.13097E-05** | **0.000280028** | **0.497818531** | **0.001977607** | **0.016127952** |

**Table 24: This table presents the mean, standard deviation and statistical analysis of Euchromatin (H3K4me3) mean major axis length values measured at various time points during anaesthesia (n=10)**

| S.No. | Control | Treatments (H3K4me3) | | | | |
| --- | --- | --- | --- | --- | --- | --- |
|  |  | **1 hour** | **2 hour** | **4 hour** | **6 hour** | **8 hour** |
| 1 | 928.0888436 | 1108.255886 | 1241.87332 | 538.2504778 | 954.3626104 | 495.241482 |
| 2 | 625.7092706 | 1012.511522 | 781.6821755 | 917.2030536 | 850.0331754 | 530.8924063 |
| 3 | 893.0628952 | 587.8858283 | 763.4400168 | 917.1341207 | 783.2699556 | 520.670446 |
| 4 | 1127.889111 | 1368.031942 | 685.2291141 | 855.6611506 | 904.6946462 | 552.6419999 |
| 5 | 925.5582918 | 495.2673858 | 701.033588 | 978.5532752 | 854.0608796 | 857.7748797 |
| 6 | 993.2005686 | 551.2643339 | 670.8020208 | 865.330848 | 1195.895058 | 462.6792035 |
| 7 | 632.8836247 | 1489.5593 | 1301.599558 | 770.6064902 | 705.2307217 | 284.7687837 |
| 8 | 630.7497645 | 935.7339811 | 1234.179358 | 1080.078286 | 834.7721653 | 648.0611698 |
| 9 | 793.252299 | 1180.452407 | 1252.572737 | 1045.171541 | 1051.17313 | 783.2130736 |
| 10 | 846.2218682 | 1008.29025 | 580.6075664 | 466.1627733 | 725.9258512 | 645.086739 |
| Mean | **839.6616537** | **973.7252836** | **921.3019455** | **843.4152016** | **885.9418194** | **578.1030184** |
| Std | **166.7946806** | **337.0950051** | **272.5040089** | **171.191882** | **147.9230993** | **161.3993632** |
| t-test |  | **0.346636285** | **0.627747336** | **0.581944795** | **0.546971203** | **0.009303551** |

**Table 25: This table presents the mean, standard deviation and statistical analysis of Euchromatin (H3K4me3) mean minor axis length values measured at various time points during anaesthesia (n=10)**

| S.No. | Control | Treatments (H3K4me3) | | | | |
| --- | --- | --- | --- | --- | --- | --- |
|  |  | **1 hour** | **2 hour** | **4 hour** | **6 hour** | **8 hour** |
| 1 | 414.20607 | 473.9790867 | 491.152468 | 367.0400205 | 388.5053897 | 292.453527 |
| 2 | 297.7458766 | 431.0501328 | 338.9733641 | 1411.7888 | 348.3173861 | 272.1343176 |
| 3 | 412.7422705 | 337.2244802 | 327.6675235 | 393.2067251 | 510.2022936 | 301.4113677 |
| 4 | 584.1620368 | 204.729429 | 389.7319254 | 451.0440827 | 434.9559155 | 292.612303 |
| 5 | 455.0402904 | 281.7296836 | 309.696849 | 468.5304596 | 369.9449256 | 436.731246 |
| 6 | 579.2674217 | 258.4473271 | 337.3771279 | 350.3165524 | 640.7551049 | 266.277576 |
| 7 | 317.892145 | 253.4050141 | 652.9596091 | 420.4702716 | 304.6618275 | 342.6214393 |
| 8 | 325.6110846 | 352.4070627 | 482.4124668 | 443.8542891 | 340.5171035 | 357.8113682 |
| 9 | 346.606807 | 610.4300811 | 651.6309576 | 675.3329952 | 691.4007496 | 400.0439565 |
| 10 | 417.0643346 | 645.9389954 | 280.4291595 | 1215.084664 | 372.3623656 | 423.2094744 |
| Mean | **415.0338337** | **384.9341293** | **426.2031451** | **619.6668861** | **440.1623062** | **338.5306576** |
| Std | **101.6849676** | **149.0620979** | **135.9285982** | **368.7344816** | **130.9142161** | **61.34206671** |
| t-test |  | **0.538647484** | **0.949506352** | **0.104699825** | **0.606696152** | **0.108033366** |

**Table 26: This table presents the mean, standard deviation and statistical analysis of Euchromatin (H3K4me3) distance from centre to periphery values measured at various time points during anaesthesia (n=10)**

| S.No. | Control | Treatments (H3K4me3) | | | | |
| --- | --- | --- | --- | --- | --- | --- |
|  |  | **1 hour** | **2 hour** | **4 hour** | **6 hour** | **8 hour** |
| 1 | 2676.005684 | 4782.387448 | 5142.550646 | 2148.872985 | 2041.972594 | 2141.151957 |
| 2 | 3480.700387 | 4741.051603 | 2409.901892 | 2180.199529 | 2945.033373 | 5453.254082 |
| 3 | 3341.335558 | 3872.595919 | 2475.411903 | 2311.168414 | 1597.288025 | 3461.976118 |
| 4 | 5541.808922 | 3912.992392 | 2799.453784 | 2425.715305 | 1781.584918 | 4379.676811 |
| 5 | 3055.490727 | 2331.421967 | 3881.991801 | 1667.287571 | 2033.557179 | 2044.535531 |
| 6 | 3893.193202 | 2411.781562 | 3735.137399 | 2645.436581 | 1841.388361 | 3682.351023 |
| 7 | 4173.078131 | 5270.810687 | 4169.492269 | 2618.694604 | 2001.549457 | 3907.398031 |
| 8 | 3832.731999 | 5291.066478 | 2202.483959 | 2421.073398 | 1972.697892 | 3500.265534 |
| 9 | 3597.35778 | 5213.202038 | 3927.678465 | 1713.701885 | 1295.969451 | 2658.346453 |
| 10 | 2822.346443 | 3433.789526 | 2392.110973 | 1974.270286 | 2694.912288 | 3420.948254 |
| Mean | **3641.404883** | **4126.109962** | **3313.621309** | **2210.642056** | **2020.595354** | **3464.99038** |
| Std | **747.040094** | **1101.935327** | **753.2239555** | **342.3374906** | **483.6867735** | **913.4577748** |
| t-test |  | **0.526822437** | **0.108154465** | **7.6203E-05** | **4.84872E-05** | **0.747618342** |

**Table 27: This table presents the mean, standard deviation and statistical analysis of Heterochromatin (H3K9me3) domain no. values measured at various time points during anaesthesia (n=10)**

| S.No. | Control | Treatments (H3K9me3) | | | | |
| --- | --- | --- | --- | --- | --- | --- |
|  |  | **1 hour** | **2 hour** | **4 hour** | **6 hour** | **8 hour** |
| 1 | 138 | 73 | 41 | 82 | 38 | 26 |
| 2 | 94 | 82 | 39 | 43 | 62 | 23 |
| 3 | 122 | 94 | 65 | 28 | 53 | 13 |
| 4 | 216 | 87 | 47 | 31 | 33 | 20 |
| 5 | 59 | 82 | 54 | 59 | 50 | 32 |
| 6 | 65 | 48 | 117 | 40 | 40 | 19 |
| 7 | 42 | 60 | 149 | 44 | 14 | 29 |
| 8 | 117 | 74 | 63 | 38 | 18 | 8 |
| 9 | 102 | 72 | 80 | 44 | 14 | 19 |
| 10 | 127 | 188 | 35 | 44 | 29 | 6 |
| Mean | **108.2** | **86** | **69** | **45.3** | **35.1** | **19.5** |
| Std | **46.94635236** | **36.23534186** | **35.25053191** | **14.59486211** | **15.88363938** | **8.114801291** |
| t-test |  | **0.276181926** | **0.060453589** | **0.001204166** | **0.000326967** | **2.66355E-05** |

**Table 28: This table presents the mean, standard deviation and statistical analysis of Heterochromatin (H3K9me3) %age area values measured at various time points during anaesthesia (n=10)**

| S.No. | Control | Treatments (H3K9me3) | | | | |
| --- | --- | --- | --- | --- | --- | --- |
|  |  | **1 hour** | **2 hour** | **4 hour** | **6 hour** | **8 hour** |
| 1 | 26.36737 | 27.23815 | 26.35638 | 13.01311 | 11.36819 | 13.8413 |
| 2 | 38.9371 | 23.96621 | 18.52706 | 19.37001 | 16.73767 | 12.94778 |
| 3 | 23.25051 | 29.60334 | 21.48247 | 22.28248 | 16.35436 | 9.86572 |
| 4 | 32.2982 | 29.72251 | 22.4457 | 21.01409 | 19.5372 | 12.88222 |
| 5 | 31.29455 | 22.52252 | 27.41492 | 19.25011 | 14.77955 | 16.69028 |
| 6 | 34.82637 | 27.95003 | 22.21744 | 20.12879 | 18.63787 | 10.97318 |
| 7 | 25.66282 | 27.48692 | 25.59553 | 21.3334 | 16.10363 | 10.8221 |
| 8 | 39.8927 | 22.21051 | 21.79454 | 17.40376 | 14.35375 | 18.32632 |
| 9 | 39.73012 | 30.82315 | 25.50014 | 22.09237 | 13.32248 | 15.41611 |
| 10 | 33.69127112 | 24.17996459 | 21.35677417 | 14.22309845 | 20.85900101 | 18.28865439 |
| Mean | **32.59510111** | **26.57033046** | **23.26909542** | **19.01112185** | **16.2053701** | **14.00536644** |
| Std | **5.714076183** | **2.971525588** | **2.65336053** | **3.041722472** | **2.759730888** | **2.916968132** |
| t-test |  | **0.011677152** | **0.000315601** | **6.18882E-06** | **3.85049E-07** | **7.36318E-08** |

**Table 29: This table presents the mean, standard deviation and statistical analysis of Heterochromatin (H3K9me3) mean domain area values measured at various time points during anaesthesia (n=10)**

| S.No. | Control | Treatments (H3K9me3) | | | | |
| --- | --- | --- | --- | --- | --- | --- |
|  |  | **1 hour** | **2 hour** | **4 hour** | **6 hour** | **8 hour** |
| 1 | 399304.3478 | 262465.7534 | 236565.8537 | 218848.7805 | 150189.4737 | 191323.0769 |
| 2 | 327336.1702 | 416273.1707 | 263589.7436 | 377079.0698 | 265625.8065 | 190622.4281 |
| 3 | 347803.2787 | 506365.9574 | 546769.2308 | 340914.2857 | 248724.5283 | 258276.9231 |
| 4 | 335518.5185 | 379843.6782 | 484765.9574 | 400929.0323 | 240183.0808 | 335920 |
| 5 | 116691.5254 | 456097.561 | 483644.4444 | 382847.4576 | 233952 | 323650 |
| 6 | 514756.9231 | 316100 | 439329.9145 | 373840 | 289640 | 270273.6842 |
| 7 | 665942.8571 | 322720 | 28622.81879 | 204763.6364 | 310285.7143 | 376220.6897 |
| 8 | 420335.0427 | 455740.5405 | 249244.4444 | 347789.4737 | 242311.1111 | 197200 |
| 9 | 306415.6863 | 446222.2222 | 634080 | 214400 | 437371.429 | 191705.2632 |
| 10 | 171779.5276 | 403131.9149 | 251565.7143 | 293818.1818 | 303420.6897 | 150933.333 |
| Mean | **360588.3877** | **396496.0798** | **361817.8122** | **315522.9918** | **272170.3833** | **248612.5398** |
| Std | **149011.641** | **72242.20375** | **174780.9559** | **72810.12257** | **69909.38671** | **72227.07863** |
| t-test |  | **0.523585699** | **0.987364521** | **0.425625701** | **0.124456015** | **0.057555307** |

**Table 30: This table presents the mean, standard deviation and statistical analysis of Heterochromatin (H3K9me3) mean domain perimeter values measured at various time points during anaesthesia (n=10)**

| S.No. | Control | Treatments (H3K9me3) | | | | |
| --- | --- | --- | --- | --- | --- | --- |
|  |  | **1 hour** | **2 hour** | **4 hour** | **6 hour** | **8 hour** |
| 1 | 2889.114783 | 2284.173699 | 2135.334634 | 1790.569756 | 3225.661053 | 1244.861538 |
| 2 | 2395.530213 | 2007.059512 | 2389.025641 | 2912.869767 | 2283.993548 | 866.3235592 |
| 3 | 2731.483934 | 2564.099574 | 2128.108308 | 1407.692857 | 2199.906415 | 606.5661538 |
| 4 | 2657.097778 | 2709.865747 | 3567.372766 | 1357.341935 | 2363.281566 | 1266.604 |
| 5 | 6722.47661 | 2575.287805 | 3536.154815 | 3245.67322 | 1847.5344 | 2558.82 |
| 6 | 3825.479385 | 2713.024167 | 2008.024957 | 2876.863 | 2342.613 | 311.5705263 |
| 7 | 5071.412381 | 2786.976 | 1053.896107 | 1874.84 | 1176.385714 | 1112.355862 |
| 8 | 3105.902564 | 3173.816757 | 2131.145397 | 2686.611579 | 1773.797778 | 1954.65 |
| 9 | 2556.655294 | 2120.2 | 1937.063 | 1886.462727 | 1966.005714 | 2521.389474 |
| 10 | 1683.624882 | 1913.738085 | 2140.221714 | 2475.130909 | 2572.342069 | 1364.746667 |
| Mean | **3363.877782** | **2484.824135** | **2302.634734** | **2251.405575** | **2175.152126** | **1380.788778** |
| Std | **1414.669664** | **375.6066431** | **708.7043543** | **636.8133807** | **514.4821834** | **716.1839826** |
| t-test |  | **0.088362435** | **0.059416756** | **0.045285978** | **0.029221682** | **0.001459429** |

**Table 31: This table presents the mean, standard deviation and statistical analysis of Heterochromatin (H3K9me3) mean circularity values measured at various time points during anaesthesia (n=10)**

| S.No. | Control | Treatments (H3K9me3) | | | | |
| --- | --- | --- | --- | --- | --- | --- |
|  |  | **1 hour** | **2 hour** | **4 hour** | **6 hour** | **8 hour** |
| 1 | 0.701745619 | 0.637513686 | 0.770097397 | 0.550014655 | 0.484354806 | 0.670898878 |
| 2 | 0.704009204 | 0.674396944 | 0.607702486 | 0.560541821 | 0.696365213 | 0.696872282 |
| 3 | 0.673912624 | 0.592843903 | 0.449473131 | 0.419169544 | 0.645182181 | 0.537646075 |
| 4 | 0.685107869 | 0.62193363 | 0.615137415 | 0.385665117 | 0.636451019 | 0.777446198 |
| 5 | 0.702147621 | 0.712858347 | 0.606960767 | 0.666572125 | 0.341127644 | 0.726126021 |
| 6 | 0.571015811 | 0.674599481 | 0.510656336 | 0.658162133 | 0.6653281 | 0.601192442 |
| 7 | 0.597054318 | 0.706825239 | 0.428788995 | 0.514189187 | 0.608585 | 0.611471015 |
| 8 | 0.622330204 | 0.717689205 | 0.676808962 | 0.235478197 | 0.384848479 | 0.524999246 |
| 9 | 0.668359946 | 0.601843602 | 0.499825587 | 0.392536609 | 0.661474136 | 0.607212933 |
| 10 | 0.722644979 | 0.624449594 | 0.393203149 | 0.618101141 | 0.275870538 | 0.745586194 |
| Mean | **0.66483282** | **0.656495363** | **0.555865423** | **0.500043053** | **0.539958712** | **0.649945128** |
| Std | **0.048311462** | **0.044410434** | **0.113333464** | **0.131986249** | **0.147127877** | **0.082312912** |
| t-test |  | **0.707548566** | **0.016172014** | **0.002459897** | **0.026368216** | **0.645433474** |

**Table 32: This table presents the mean, standard deviation and statistical analysis of Heterochromatin (H3K9me3) mean major axis length values measured at various time points during anaesthesia (n=10)**

| S.No. | Control | Treatments (H3K9me3) | | | | |
| --- | --- | --- | --- | --- | --- | --- |
|  |  | **1 hour** | **2 hour** | **4 hour** | **6 hour** | **8 hour** |
| 1 | 962.2561574 | 857.8016426 | 861.4828521 | 720.7088966 | 685.9816134 | 848.7405664 |
| 2 | 871.3105054 | 798.418163 | 918.6716234 | 618.1933406 | 773.35923 | 465.5958648 |
| 3 | 918.5135663 | 847.274433 | 800.752652 | 1253.563859 | 840.1254756 | 621.1495949 |
| 4 | 840.4860233 | 843.2526537 | 938.2976245 | 661.0329029 | 482.7537453 | 917.9750336 |
| 5 | 1633.710806 | 707.8052343 | 830.1267911 | 944.6214959 | 780.7068182 | 718.4133607 |
| 6 | 1263.06714 | 933.6517023 | 734.923203 | 780.8386912 | 613.5637858 | 313.5274438 |
| 7 | 1278.074954 | 840.6611421 | 427.7486761 | 724.915193 | 1234.454672 | 989.4554437 |
| 8 | 1017.966569 | 965.954348 | 730.6352931 | 532.2233806 | 1034.069375 | 793.2220038 |
| 9 | 899.5758634 | 750.448703 | 693.9433665 | 757.864488 | 588.128083 | 875.1513279 |
| 10 | 695.3743168 | 727.1424423 | 763.9331165 | 678.4177866 | 377.008067 | 488.2802985 |
| Mean | **1038.03359** | **827.2410464** | **770.0515198** | **767.2380034** | **741.0150865** | **703.1510938** |
| Std | **274.9845543** | **83.2073107** | **141.3452002** | **201.7743488** | **253.9224876** | **217.2624957** |
| t-test |  | **0.043552999** | **0.018514882** | **0.037192682** | **0.03797835** | **0.010414025** |

**Table 33: This table presents the mean, standard deviation and statistical analysis of Heterochromatin (H3K9me3) mean minor axis length values measured at various time points during anaesthesia (n=10)**

| S.No. | Control | Treatments (H3K9me3) | | | | |
| --- | --- | --- | --- | --- | --- | --- |
|  |  | **1 hour** | **2 hour** | **4 hour** | **6 hour** | **8 hour** |
| 1 | 470.8485003 | 407.1599831 | 375.8172677 | 349.9140267 | 445.9016111 | 402.7905508 |
| 2 | 377.9839383 | 323.4632124 | 417.3394912 | 442.0831123 | 426.0565546 | 530.9865769 |
| 3 | 429.500865 | 411.9742811 | 417.5901351 | 618.769883 | 367.8136818 | 355.4738778 |
| 4 | 421.1939559 | 426.3083265 | 478.4911769 | 443.0782913 | 489.4068116 | 431.7631943 |
| 5 | 753.6769035 | 379.6064316 | 505.4424057 | 474.5108602 | 351.6270342 | 415.9742951 |
| 6 | 599.0561326 | 474.4379881 | 368.5975164 | 477.7588139 | 388.49685 | 585.7225088 |
| 7 | 629.0054511 | 398.5589688 | 224.7025736 | 373.667198 | 675.5598129 | 513.9003505 |
| 8 | 480.0417065 | 434.7400025 | 422.3893789 | 488.1357808 | 652.8624018 | 871.3208939 |
| 9 | 450.2131933 | 372.8253045 | 417.9284245 | 348.5971412 | 693.6992997 | 477.7437909 |
| 10 | 311.1080951 | 336.7268305 | 358.8387751 | 457.6894695 | 549.1797568 | 768.4775911 |
| Mean | **492.2628742** | **396.5801329** | **398.7137145** | **447.4204577** | **504.0603815** | **535.415363** |
| Std | **131.3070181** | **45.22922557** | **76.08990417** | **72.24546504** | **129.3703938** | **159.4868036** |
| t-test |  | **0.060329535** | **0.100973702** | **0.502085165** | **0.810583609** | **0.458311227** |

**Table 34: This table presents the mean, standard deviation and statistical analysis of Heterochromatin (H3K9me3) distance from centre to periphery values measured at various time points during anaesthesia (n=10)**

| S.No. | Control | Treatments (H3K9me3) | | | | |
| --- | --- | --- | --- | --- | --- | --- |
|  |  | **1 hour** | **2 hour** | **4 hour** | **6 hour** | **8 hour** |
| 1 | 5481.628188 | 3794.534476 | 2930.155082 | 4788.597348 | 3041.342678 | 2303.260098 |
| 2 | 4281.521841 | 4445.703183 | 2920.383229 | 3413.72237 | 4065.549536 | 2717.085977 |
| 3 | 5654.418392 | 3966.787468 | 4795.158932 | 3168.294605 | 3594.392082 | 1728.847217 |
| 4 | 6598.448705 | 4091.40849 | 4230.351293 | 2603.980776 | 2988.592722 | 2207.646178 |
| 5 | 5650.86589 | 3982.432225 | 4682.610845 | 4028.564859 | 3129.930724 | 2337.816069 |
| 6 | 4757.130536 | 3073.733333 | 5308.878234 | 2970.222178 | 2723.366941 | 2421.183563 |
| 7 | 4314.095064 | 3269.754884 | 3229.583096 | 3364.109513 | 2309.474423 | 2704.593456 |
| 8 | 5591.342024 | 4322.620676 | 4144.179692 | 2650.41156 | 2488.243427 | 2554.398006 |
| 9 | 4714.635441 | 3816.135665 | 3684.295243 | 3554.956931 | 2559.996416 | 1983.711506 |
| 10 | 5000.955835 | 5867.928915 | 2279.265466 | 3724.964458 | 3028.252596 | 1525.378762 |
| Mean | **5204.504192** | **4063.103931** | **3820.486111** | **3426.78246** | **2992.914154** | **2248.392083** |
| Std | **715.8619232** | **756.3763103** | **921.7440163** | **449.7578842** | **528.5156012** | **397.679868** |
| t-test |  | **0.009703088** | **0.007803135** | **9.62615E-06** | **3.27632E-06** | **2.3107E-08** |
